# Influence of F Repeats and Terminal Residue Polarity on Peptide–VIM-2 Metallo-β-Lactamase Interactions

**DOI:** 10.1101/2025.06.15.659760

**Authors:** Ananya Anurag Anand, Sarfraz Anwar, Rajat Kumar Mondal, Sintu Kumar Samanta

## Abstract

The global surge in antimicrobial resistance is a major public health concern, largely driven by the dissemination of beta-lactamases. Among them, VIM-2 metallo-beta-lactamase (MBL) poses a significant therapeutic challenge. In this study, we explore the role of phenylalanine and lysine residues, especially their repeat patterns, in modulating peptide binding affinity and structural interactions with VIM-2. Docking, molecular dynamics simulations, free energy decomposition analysis and residue analysis were conducted on gut metagenome-derived AMPs, PolyF and a control peptide (PolyR). Binding affinity with VIM-2 decreased with declining F content, yet PolyF alone showed unexpectedly low binding. However, PolyF exhibited most negative polar solvation energy in free energy decomposition analysis. This indicates that F repeats play a role in stabilizing a VIM-2-peptide complex in presence of a solvent. PolyF peptide lacks favorable electrostatic or dynamic interactions, likely due to the absence of K residues. Interestingly, loop1 and loop2 analysis of VIM-2 revealed that F repeats interfere with the loops that play a role in functioning of substrate binding at the active site. Additionally, F repeats were found to bind to residues lying adjacent to or nearby active site residues, which points towards the fact that F helps other residues like K to bind more stably with active site residues of VIM-2. Finally, analog designing and analysis reveals that a balance of aromatic (F) and positively charged (K) residues could enhance peptide binding with MBLs like VIM-2 when having hydrophobic on N-terminal and positively charged on C-terminal.

## Introduction

The alarming rise of antimicrobial resistance (AMR) has emerged as one of the most critical global health concerns **(Queenan and Bush, 2007; Mondal et al., 2023).** Among the most formidable contributors to this crisis are metallo-β-lactamases (MBLs), enzymes capable of hydrolyzing a wide spectrum of β-lactam antibiotics, including carbapenems—often considered the last line of defense against multidrug-resistant bacterial infections **(Mojica et al., 2021).** Verona integron-encoded metallo-β-lactamase-2 (VIM-2) is a clinically significant subclass B1 MBL that has been increasingly reported in Gram-negative pathogens, especially *Pseudomonas aeruginosa* and members of the Enterobacteriaceae family, which are a part of ESKAPE group **(Anurag Anand et al., 2025a; Anurag Anand et al., 2025b).** These infections pose severe therapeutic challenges due to the limited efficacy of available inhibitors against MBL producers, necessitating the urgent development of novel ones to restore β-lactam antibiotic potency.

Peptide-based inhibitors have gained considerable interest due to their multi-traget nature, target specificity, and less tendency to induce resistance in bacteria against it **(Rossino et al., 2023; Anurag Anand et al., 2023).** Among the wide array of amino acid sequences explored, the repetitive presence of particular residues such as arginine (R) and lysine (K) has shown promise in modulating binding interactions with enzymatic targets **(Rotondo et al., 2015; Anurag Anand et al., 2024)**. Lysine and arginine, with their positively charged side chain, play a pivotal role in electrostatic interactions and hydrogen bonding, potentially enhancing binding affinity through complementary charge distribution with negatively charged pockets on protein surfaces.

This study aims to elucidate the contribution of other repeats, particularly F repeats, through a combination of molecular docking, MD simulations, residue analysis and binding affinity predictions **(Pingali et al., 2023; Singh et al., 2023; Xiao et al., 2015)**. By systematically analyzing peptides with varying F and K compositions, we aim have tried to uncover sequence-specific features that help in binding and stabilizing the complex of of peptide-based inhibitors with VIM-2.

Preliminary analyses of peptide inhibitors interacting with VIM-2 revealed that sequences enriched in F and K residues tend to exhibit improved binding affinities. Notably, AMP13— containing 14 phenylalanine residues and several lysine repeats—demonstrated higher affinity towards VIM-2 compared to AMP18 and AMP10, which contain 7 and 4 F residues respectively. Intriguingly, a control peptide composed of 30 continuous F residues (PolyF) failed to show enhanced binding despite its high aromatic content, suggesting that the functional contribution of F residues is context-dependent and synergistic with other residues such as K. This observation was further supported by molecular dynamics (MD) simulations, where PolyF exhibited a stable RMSD and constant solvent-accessible surface area (SASA), yet lacked high binding affinity for the MBL.

These findings underscore a crucial insight that it is not merely the abundance of F or K residues but their spatial arrangement and interplay with surrounding residues that governs inhibitory potency. The dual contribution of hydrophobicity from F and electrostatic complementarity from K may together establish a robust framework for designing potent peptide inhibitors against MBLs like VIM-2. Our study shows that the structural flexibility endowed by a mixed amino acid composition might enable conformational adaptability necessary for effective enzyme inhibition. Finally, analog designing and analysis reveals that a balance of aromatic (F) and positively charged (K) residues could enhance peptide binding with MBLs like VIM-2 when having hydrophobic on N-terminal and positively charged on C-terminal.

Through these insights, we hope to contribute to the rational design of next-generation peptide inhibitors that are not only potent but also capable of overcoming the adaptive resistance mechanisms of VIM-2 and related β-lactamases (**Livermore, 2009; Anurag Anand et al., 2025**). As the global antibiotic pipeline continues to shrink, such molecular investigations are vital in steering the future of antimicrobial therapy toward precision-based, resistance-counteracting solutions.

### Methodology

#### Literature survey and Target Dossier Study

Target Dossier Study is the comprehensive analysis of a specific biological target, typically crucial for drug discovery and development purposes **(Anurag Anand et al., 2023)**. The objectives of a target dossier study are to: 1) Identify and validate potential drug targets, 2) Understand target structure, function, and regulation, 3) Characterize target-ligand interactions, and 4) Inform drug design and optimization. Target dossier study offers several benefits, like: 1) Informed drug discovery decisions, 2) Reduced risk of target-related failures, 3) Enhanced understanding of target biology and 4) Improved drug efficacy and safety

We performed a detailed Target Dossier Study, focusing on target selection, validation, and characterization. Our study consisted of four parts: 1) Literature review 2) Target identification 3) Comparative study of target structures, 4) Target structure and functional analysis, and 5) Ligand binding site analysis

Literature review was undertaken for providing context and identifying gaps in existing research **(Rotondo et al., 2015)**. After reviewing the literature, identification of target structures was performed. With targets identified, a comparative study of target structures available on PDB was done in order to explore the similarities and differences among them. It played as a crucial step in understanding their relationships and relevance. Once a strong foundation was laid, we proceeded to target structure and functional analysis, which involved examining how these structures function within their respective contexts. An in-depth ligand-binding site analysis was then carried out. The PDB database was utilized to obtain publicly available structures of the enzyme (https://www.rcsb.org/). X-ray crystallography reports were studied in detail so as to obtain the details regarding the same.

#### Selection of sequences and peptide modeling

In one of our ongoing works (data confidential; not yet published), we have found three lead candidates (gut-derived antimicrobial peptides (AMPs)) that have the potency to bind very tightly with VIM-2. These peptides were found to possess F and K repeats occurring in them naturally. In one of our previous studies we had already studied the role of K repeats in binding with VIM-2. Therefore, here our main focus is the F repeats. Thus, the three-dimensional structures of our candidate AMPs (AMP13, AMP18, AMP10) as well as PolyF (30 F residues) were predicted using the PEP-FOLD3 server (http://bioserv.rpbs.univ-paris-diderot.fr/services/PEP-FOLD3/). PolyR was kept as a control **(Anurag Anand et al., 2024; Rotondo et al., 2015)**. This *ab initio* approach predicts peptide conformations based on sequence information and fragment library assembly using Hidden Markov Models. Each peptide sequence was submitted in linear format, and a series of 200 simulations was run. The top-ranked model based on sOPEP energy scoring was selected for further studies. Structural quality was visually inspected using PyMOL, and models were minimized before docking.

#### Protein–Peptide Docking

Protein–peptide docking was performed using the HawkDock server (http://www.cadd.zju.edu.cn/hawkdock/), a hybrid computational platform combining molecular docking and MM/GBSA binding free energy prediction. The crystal structure of VIM-2 (PDB ID: 1KO3) was retrieved from the RCSB Protein Data Bank and prepared by removing water molecules, ligands, and non-standard residues using PyMOL, and then subjected to Protein-peptide docking. Protein-peptide blind docking between VIM-2 and peptides was performed using the HAWKDOCK server **(Weng et al. 2019).** The HAWKDOCK server utilizes the ATTRACT docking algorithm which is based on minimization of energy in rotational and translational degrees of freedom of one protein with respect to peptide.

#### MMGBSA free energy and residue decomposition analysis

The models with high binding affinity in docking were subjected to MM/GBSA using HAWKDOCK itself so as to obtain top complexes with the least binding free energy. In addition to this, MM/GBSA free energy decomposition analysis and residue analysis were performed to extract the contribution of each residue to the total binding free energy **(Dalal et al. 2021; Kumari and Dalal 2022).** The complexes were selected for MD Simulation studies based on the more negative MM/GBSA scores which represent tight binding with the enzyme.

#### Molecular Dynamics (MD) Simulation of Protein-Peptide Complexes

The protein-peptide complexes were prepared in the aqueous region of the TIP3P model, using Gromacs version 2019.**4** (https://www.gromacs.org/). The complex was prepared in two different states i.e., unbound protein and peptide bound protein. Protein structures were taken from the docked complex (**Singh et al., 2022**). charmm36-mar2019.ff force field was selected for generating the topology. In case of the unbound protein, charmm36-mar2019.ff force field was selected for generating protein topology. A cubic box with a size of 1 nm was set. Conjugate gradient energy minimization method of GROMACS was employed to minimize all the systems in order to remove any initial stress. Equilibration of the system was performed at 300 K temperature for a duration of 100 ps in case of NVT ensemble. This was followed by another equilibration of 100 ps for NPT ensemble. Nosé–Hoover thermostat was employed to control the system’s tempera ture at a coupling constant of 0.1 ps **(Evans et al. 1985).** On the other hand, Parrinello–Rahman barostat was used to control the pressure of the system at an isotropic coupling constant of 2 ps **(Parrinello& Rahman 1981).** In the end, a long production trajectory in the NVT ensemble was generated for each of the systems for a duration of 100 ns. For further analysis, all the trajectories were saved at an interval of 10 ps. Periodic boundary conditions (PBC) was applied to minimize the boundary effects. Calculation of long-range electrostatic interaction was done by using the Particle mesh Ewald (PME) method **(Essmann et al. 1995).**

#### Deriving structural order parameters and essential dynamics

In order to perform the MD analysis, GROMACS utilities were employed to obtain the trajectories **(Anurag Anand et al., 2024).** Hydrogen bond interactions, the radius of gyration (Rg), root mean square deviation (RMSD), and root mean square fluctuation (RMSF) were measured. PC1 and PC2 projections of the protein structure were considered for principal component analysis (PCA). gmx-sham was utilized to generate free energy landscape (FEL) plots **(Prakash et al. 2018).** Snap shots at different intervals were extracted to check the binding stability of peptide to the protein **(Dalal and Kumari 2022; Dalal et al. 2021).**

#### Studying the instability index

The instability index of the peptides was extracted using Protparam software, to correlate the interaction dynamics with the stability of the peptide in each case.

#### Protein-Peptide Interaction Analysis

The protein-peptide interaction plots showing non-covalent interactions like H-bonds and salt-bridges were extracted using Ligplus **(Dalal and Kumari 2022).**

#### Measuring the effect of smaller F repeats and comparison with repeats of other amino acids

Amino acid repeats were subjected to HAWKDOCK for docking using the protocol mentioned above. Subsequent MMGBSA analysis was performed as mentioned above. CABSDOCK’s coarse-grained model was further used to simulate the repeats in complex with VIM-2. CABS-dock utilizes a coarse-grained protein model called CABS for flexible protein-peptide docking. Instead of using all atoms of a protein, CABS represents each amino acid with a few pseudo-atoms, such as the alpha-carbon, beta-carbon, and side-chain center. The reduced complexity of the CABS model enables faster and more efficient simulations compared to all-atom models, especially when dealing with flexible proteins and peptides. Once the CABS algorithm has performed the fliexible docking, it simulates the complex to obtain all possible conformations upto 10,000. Further, 1000 most probable conformations are selected and subjected to clustering. Finally, 10 representative models are obtained. Interaction energy of all models (10 models) and model 1 (topmost model) were studied. Model 1 was given priority when interpreting the results. The interaction energy points were plotted for model 1 for all 1000 trajectory (replica) frame index. Similarly, when studying for all models, interaction energy of all 10,000 trajectory (replica) frame index were plotted and studied. A total of 50 simulation cycles were run to obtain reliable results.

#### Deriving ultra-short peptides and studying the effect of F monopeptide and F repeats in presence of other amino acids

From the parent AMPs, AMP10, AMP13 and AMP18, the binding domains were identified using Ligplot analysis and residue analysis based on MMGBSA Binding Free Energy. These binding domains were taken separately to form a library of ultra-short and short peptides (Library-I). The analog library I consisted of three sets each for each of the parent peptides (that is 3 parents lead to total 9 sets of peptides, meaning 3 each). The three sets were constructed as follows: set 1: Where single F (F monopeptide) is present in the sequence, set 2: Where F dipeptide is a part of the peptide sequence, set 3: where F tripeptide is a part of the sequence. Some random sets were also constructed with F in the form of larger repeats. The effect of F monopeptide and F repeats (di, tri or larger) in presence of other amino acids was studied using docking and simulation analysis performed using CABSDOCK itself, using the same protocol as mentioned above. Consequently, Library II was formed by identifying probable efficient analogs within analogs (AWA). The effect of F monopeptide and F repeats (di, tri or larger) was again studied in presence of other amino acids in the AWA sequences. Finally, Library-III was prepared by designing analogs based on the insights from the previous libraries. The effect of F monopeptide and F repeats (di, tri or larger) was finally studied once again in presence of other amino acids, for the Library-III sequences this time.

## Results and Discussion

### Selection of the representative MBL

Identifying the representative of MBLs was crucial when developing inhibitors. One of the key beta-lactamase enzymes, is the Verona integron-encoded metallo-beta-lactamase-2 (VIM-2) enzyme **(Poirel et al., 2000).** VIM-2 belongs to a class of enzymes (Ambler Class B) that can break down a wide range of beta-lactam antibiotics, such as penicillins, cephalosporins, and carbapenems. The spread of VIM-2-producing strains of *P. aeruginosa* in hospitals is a growing concern because it makes many common antibiotics ineffective. The gene that codes for VIM-2, known as blaVIM-2, is located on mobile genetic elements like integrons, which enable it to be transferred easily between different bacterial species **(Bhat et al., 2023).** This ability to move between bacteria has allowed the resistance gene to spread quickly in both clinical and environmental settings, making VIM-2 a prime target for new therapies aimed at overcoming beta-lactam resistance. The main reason behind choosing VIM-2 as a representative of other MBLs was that: 1) it shares characteristics of both IMP and NDM, two other most common MBLs, and 2) it is a growing concern in ESKAPE **(Rotondo et al., 2015)**. In a previous study, genomic and phenotypic tests were conducted across various clinical isolates of *P. aeruginosa,* to confirm VIM-2 as a strong target. The blaVIM-2 gene showed minimal changes between different strains, highlighting its significance as a target for broad-spectrum inhibitors **(Toleman et al., 2007).** Structural studies using X-ray crystallography have provided detailed images of VIM-2’s active site **(Yamaguchi et al., 2007).** These studies have identified key, conserved motifs around the zinc-binding site that are critical for the enzyme’s function. Because of this structural consistency, inhibitors designed to target these motifs could have wide-ranging effectiveness in combating beta-lactam resistance.

*P. aeruginosa* has been our pathogen of interest because it is an increasing problem in patients of burn and cystic fibrosis (**Anurag Anand et el., 2023**).

### Target Structure and Functional Analysis

The enzyme VIM-2 exists in two states: oxidized and reduced (native). The structural complexity of the VIM-2 enzyme, especially when comparing its native and oxidized forms, offers important insights into how it works **(Garcia-Saez et al., 2008).** By examining the enzyme’s crystal structures, we get a clear picture of its intricate details.

In its native, reduced form, VIM-2 is made up of 231 amino acids, which fold into a classic αβ/βα sandwich structure, a typical feature of the metallo-β-lactamase (MBL) family **(Garcia-Saez et al., 2008)**. This structure contains two critical metal-binding sites known as the His and Cys sites. Each of these sites holds a zinc ion (Zn1 and Zn2), positioned about 4.2 Å apart. In the reduced form, Zn2 is less stable than Zn1, leading to some functional differences. Additionally, near the active site, a chloride ion is bound to a water molecule, closely interacting with Zn2.

However, in the oxidized form of VIM-2, the biggest difference is that Zn2 is no longer present, leaving only Zn1 in the His site **(Garcia-Saez et al., 2008; Rotondo et al., 2015)**. Meanwhile, the cysteine residue in the Cys site becomes oxidized, turning into a cysteine-sulfonic form. This oxidation of cysteine (Cys221) leads to subtle but significant shifts in the enzyme’s active site. Neighboring amino acids like His263, Tyr67, Tyr224, and Arg228 also change their positions to make room for the larger cysteine-sulfonic group. These shifts could affect how the enzyme performs. For example, His263 rotates 60° to interact with the sulfonic group, while Arg228 and Tyr224 move to new positions, creating van der Waals interactions that stabilize this new form.

These structural differences between the oxidized and reduced forms of VIM-2 aren’t just minor adjustments—they suggest potential changes in function. Specifically, the oxidation of Cys221 might reduce the enzyme’s activity. The lack of Zn2 in the Cys site implies that the oxidized form could either resemble an enzyme-substrate complex or represent an inactive state.

Study by Rotondo and colleagues shows that PolyR which is a peptide containing R repeats (composed of 30 Arginine or R residues) binds most strongly with VIM-2 MBL (**Rotondo et al., 2015)**. We have therefore taken PolyR as a control in our study.

### Analyzing and Comparing Different PDB Structures for VIM-2

12 different structures of VIM-2 from *P. aeruginosa* were studied against each other. A comparison of different PDB structures of VIM-2 from *P. aeruginosa* has been given in detail in **Table S1 (PDB structures retrieved from** https://www.rcsb.org). Except 1KO2, 7DZ0 and 7DZ1, all structures show a resolution of less than 2 Å. 1KO3 and 4NQ2 showed no Ramachandran outlier, indicating that all residues lie in the allowed regions. Additionally, 4NQ2, 1KO3 an 1KO2 are the only candidates with no extra ligand attached apart from small molecules like Zn, Cl, Na, OH, etc. which are usually naturally present around VIM-2. Presence of extra ligands might modify the original functional structure of an enzyme.

We further studied the fraction of modeled and unmodeled regions of different VIM-2 structures **(Table S2)**. 4NQ2 had a drawback that more than 10% of the region in it was unmodelled. 1KO3 seemed to be an optimal candidate having 0% region not modelled. Out of 1KO3 and 1KO2, we prefer 1KO2 as it lies in its native/reduced state which is crucial for its functioning. Although 1KO3 and 7DUY have 0 % unmodelled region and 93% modelled region which is much higher than that of 1KO3, yet 1KO3 seems to be a better representative of the native conformation of the enzyme since it consists of reduced Cys, that is lies in its native state, alongside having no extra ligands attached to it (**Garcia-Saez et al., 2008).**

### Ligand binding-site analysis of 1KO3 structure of VIM-2

The 1KO3 protein structure contains several small molecules and ions that are already bound to specific regions of the protein. These pre-attached ligands include acetate ions, chloride ions, hydroxide ions, and zinc ions. All of these ligands are located in Chain A of the protein.

Zinc ions in the protein serve as vital anchors, forming multiple coordination bonds with surrounding residues and other ligands.

VIM-2 requires zinc ions (Zn²LJ) in its active site to function **(Christopeit et al., 2016).** These zinc ions are necessary for breaking the amide bond in the beta-lactam ring, which deactivates the antibiotic **(Page et al., 2008; Aitha et al., 2014).** The zinc ions interact with important amino acids, like histidine, aspartate, and cysteine, in the active site. This reliance on zinc gives researchers an opportunity to develop inhibitors that can block the enzyme’s activity by disrupting the zinc coordination, restoring the effectiveness of beta-lactam antibiotics.

Zn1 of VIM-2 in 1KO3 forms five metal coordination bonds: 1 with Water molecule, 3 with Histidine (His116, His118 and His196), and 1 with Hydroxide ion (**Garcia-Saez et al., 2008).** These interactions help stabilize the protein’s local structure, particularly around the active site, ensuring the protein remains in the proper configuration for its function. Zn2 also forms five metal coordination bonds: 1 with Hydroxide ion, 1 with Aspartate (Asp 120), 1 with Chloride ion, 1 with Histidine (His 263), and 1 with Cysteine (Cys 221). The coordination between Zn2 and these amino acids and ligands helps stabilize a different region of the protein, where charge balance and structure are essential for maintaining proper function. Zn3 forms four metal coordination bonds, 1 with Histidine (His 170), and 2 with Acetate ions. These bonds play a key role in anchoring the acetate ions, which likely contribute to the local electrostatic environment, impacting the overall folding of this protein region. In the sections that follow we have studied the interaction of our peptides with these residues.

### Additional ligand binding site analysis (via literature survey)

In one of the studies, *rac*-2-ω-phenylpropyl-3-mercaptopropionic acid, phenylC3SH, was found to be a potent inhibitor of VIM-2. The ligand binding site of the VIM-2 enzyme reveals some fascinating details when it interacts with this compound **(Yamaguchi et al., 2007).** Two zinc ions, Zn1 and Zn2, play important roles in this interaction. Zn1 is tightly connected to three histidine residues (His116, His118, and His196), forming a stable structure with distances between the zinc and the residues ranging from 2.0 to 2.2 Å. This creates a stable environment, and the thiol group of PhenylC3SH links both zinc ions. The connection distances are about 2.5/2.2 Å and 2.1/2.2 Å between Zn1 and Zn2. This creates a well-organized binding site, which is crucial for the enzyme’s function.

Zn2 is also connected in a similar way by other residues—Asp120, Cys221, and His263. When combined with the thiol group of the inhibitor, it forms a structure that is very similar to other enzymes like IMP-1. The zinc ions remain consistently spaced apart at 3.7 to 3.8 Å, indicating that VIM-2 uses the same method as other metallo-beta-lactamases (MBLs) to bind inhibitors with thiol groups. This specific arrangement helps the enzyme to work efficiently against beta-lactam antibiotics, with Zn1 acting as the key player in breaking down these drugs.

When the inhibitor binds, we also see some significant changes in the VIM-2 enzyme structure. Loop 1 (Phe61-Ala64) and loop 2 (Ile223-Trp242), which are important for recognizing substrates, undergo major shifts. Loop 1, including residues like Tyr67 and Phe61, moves to create stronger connections with the inhibitor. Tyr67 rotates by about 27° to form a close interaction with the phenyl ring of the inhibitor, stabilizing the entire enzyme-inhibitor complex.

Loop 2 also helps stabilize the inhibitor. The carboxyl group of PhenylC3SH forms a hydrogen bond with Asn233, which is a conserved residue in most MBLs. This bond not only helps recognize the inhibitor but also causes changes in nearby residues. For example, Gly232 changes its angle, and Arg228 takes on a new shape. These shifts are crucial for how VIM-2 holds and stabilizes the inhibitor in its active site.

So we can say that when PhenylC3SH binds to VIM-2, the enzyme’s active site changes from an open space to a tunnel-like structure. This change in structure, along with the detailed interactions between the inhibitor and the enzyme, shows how VIM-2 can adjust and recognize different molecules that bind to it. This flexibility is key to its function and could help in designing future inhibitors to fight antibiotic resistance.

In the study performed by Rotondo et al., it was found that PolyR binds to active site residues like Asp87 of our protein structure (corresponding to Asp120 of original 1KO3 structure; the original structure was renumbered during docking by the software used) which are conserved in nature (**Rotondo et al., 2015**). **Table S3-S4** of the supplementary file show the details of residues in our VIM-2 structure versus residues in the original structure (PDB ID: 1KO3). All active site residues have been discussed in **Table S3.**

We aim to see how our peptides affect regions of Loop1, loop 2 and active site residues of VIM-2 (**Yamaguchi et al., 2007; Rotondo et al., 2015)**. We also aim to see if F repeats play any role in affecting the functioning of either of these.

### Identifying patterns across AMPs

Across peptides screened against VIM-2, obtained from human gut metagenome in one of our studies, we found that F repeats occur in all of them. This led us to check what role F repeats play in binding of peptides with VIM-2. We thus designed F repeats and modeled a peptide containing 30 F residues for our study. We call it PolyF. The binding of PolyF and gut-derived AMPs (AMP10, AMP13 and AMP18) to VIM-2 were studied in detail. PolyR (peptide consisting of 30 R residues) was kept as a control in our study based on the study performed by Rotondo and colleagues, as well as one of our previous studies (**Rotondo et al., 2015; Anurag Anand et al., 2024**).

### Docking and MMGBSA results for VIM-2-peptide complexes

Docking and MMGBSA results shown in **Table 1** show that PolyF shows least binding affinity with VIM-2. We also compared it against the K repeats which exist as a part of our sequence and found that K repeats bind very effectively with VIM-2 but F repeats do not.

**Table 1.**
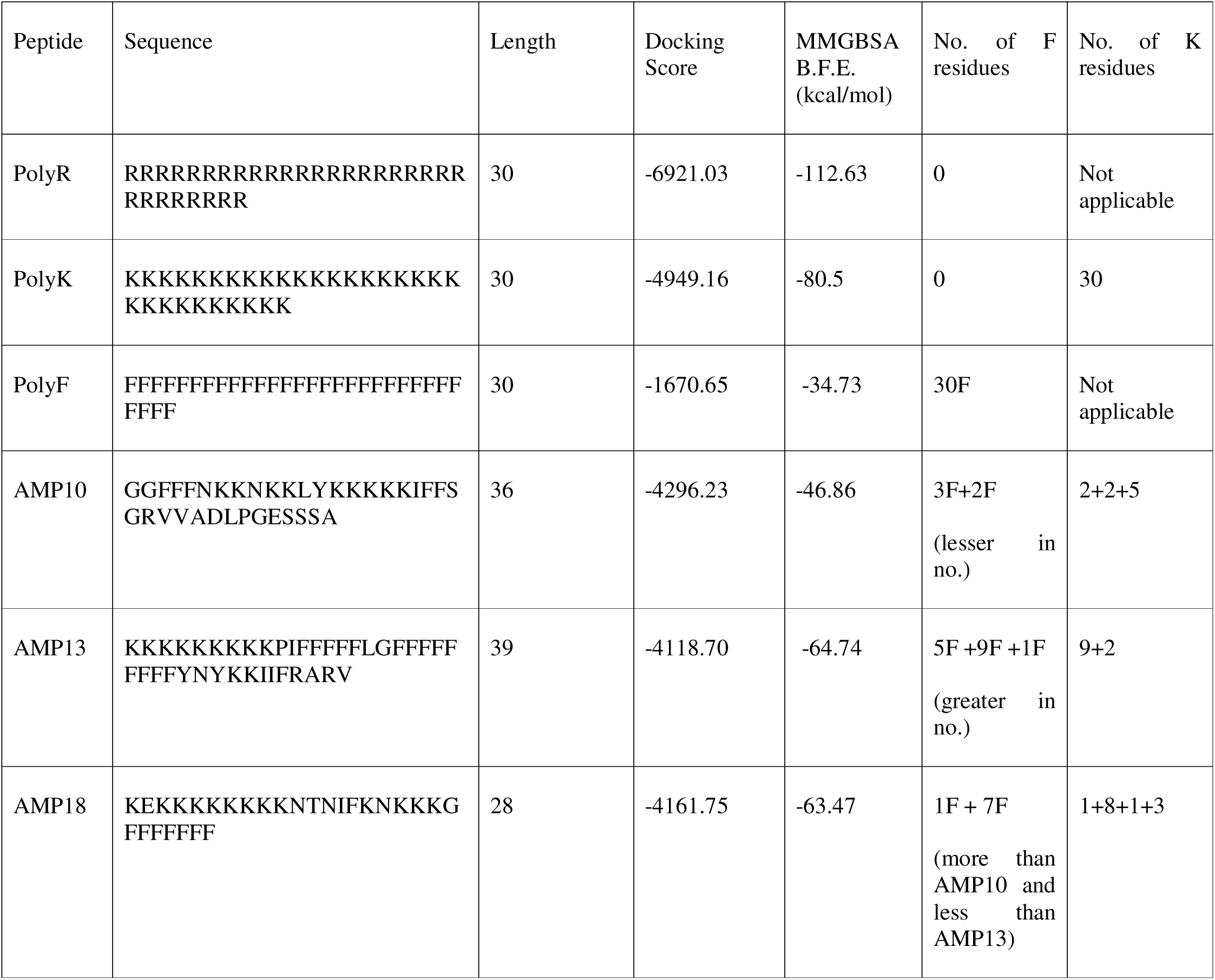
Docking and MMGBSA results for peptides in complex with VIM-2.

The docking and BFE results indicate that F repeats do not tend to bind very effectively with VIM-2, and especially do not directly aid much in binding of a peptide to VIM-2. This can be seen by more positive docking and MMGBSA BFE scores as compared to other peptides.

However, when comparing AMP10, AMP13 and AMP18, the MMGBSA BFE values represent that with an increase in the length of F repeats, BFE decreases. Since AMP10 has the least number of F residues, the BFE is most positive out of the three natural peptides being studied. On the other hand, AMP13 having the greatest number of F residues sequentially, shows the most negative BFE out of the three. However, it was surprising to note that F repeats alone show very positive BFE and docking scores as compared to the peptides containing the repeats as a part of them. We therefore, compared the docking score of PolyF and the three natural peptides with the other repeats found in our AMPs, that is PolyK. We found that K repeats have a much greater tendency to bind to VIM-2 as indicated by a more negative docking score as well as MMGBSA score. PolyR was kept as a control in our study as it has been shown to possess strong binding affinity for VIM-2 and is experimentally-validated.

### Free energy decomposition analysis

In free energy decomposition analysis, we found that the total binding free energy (Total MMGBSA B.F.E.) was the most negative for PolyR, followed by AMP13, AMP18, AMP10 and finally PolyF. Here, PolyF shows the least Total B.F.E., indicative of the fact that F repeats are not alone sufficient to bind efficiently with VIM-2 as opposed to R and K repeats which possess strong binding affinity for the MBL. We further studied the contribution of each of the energy components or type of energy to the Total B.F.E. and found that Electrostatic interaction energy follows the same trend as the Total B.F.E. indicating that stronger interactions of R repeats are driven by electrostatic bonds. The analysis of Van der Waals energy showed that it is most negative for PolyR, followed by AMP13, AMP18, PolyF and finally AMP10. SASA energy also followed the same pattern. Interestingly, PolyF shows the least polar solvation energy (PSE) followed by AMP10, AMP18, AMP13 and PolyR. A low PSE suggests that the complex is more stable in a polar solvent environment, such as water, due to favorable interactions between the polar parts of the protein-peptide complex with the polar solvent molecules. Thus, it became evident that F repeats are playing a role in stabilizing the protein-peptide complex in presence of a solvent. The same was further looked into when analyzing the MD simulation results.

The free energy decomposition analysis results have been visually depicted in **Figure 1**.

**Figure 1.**
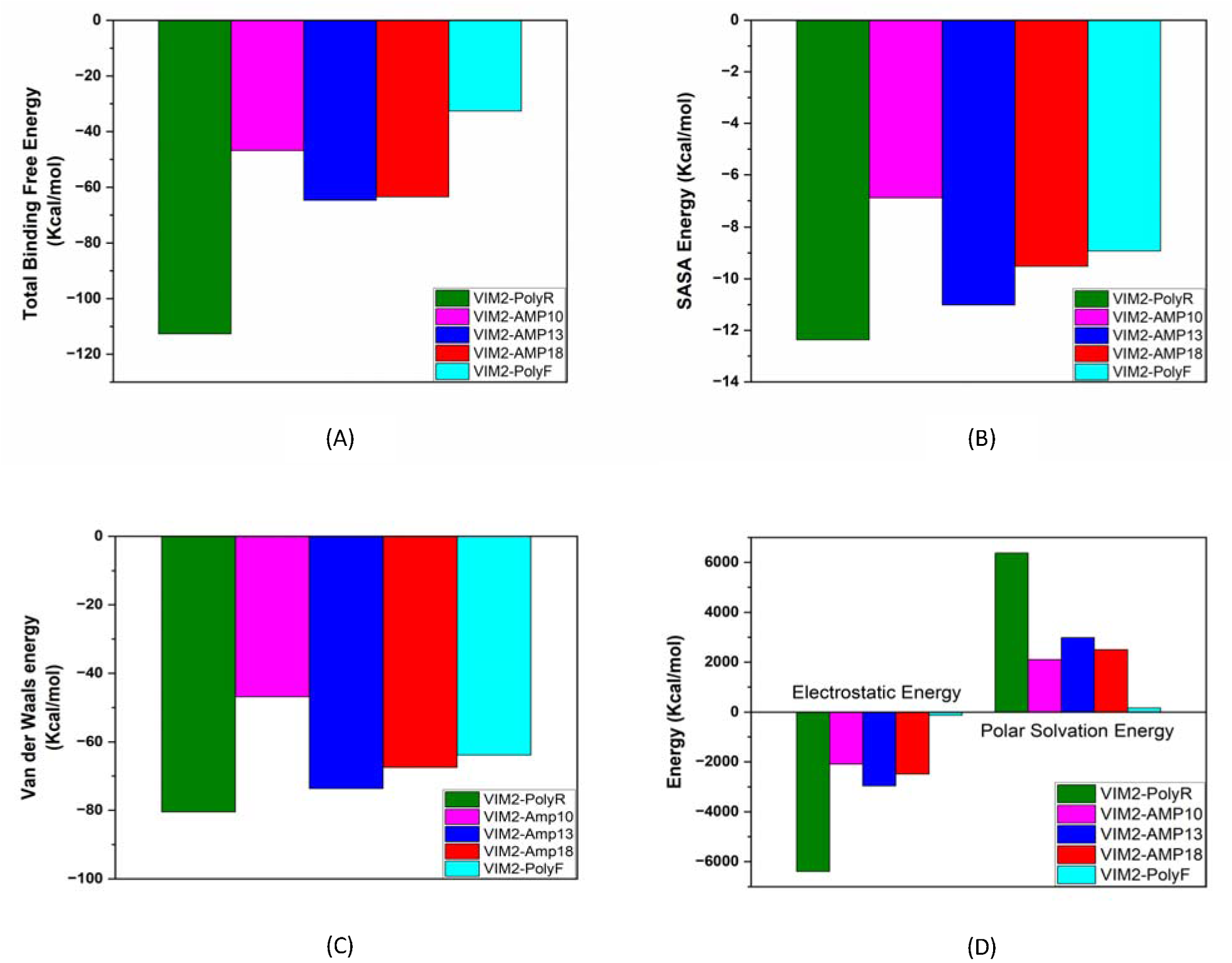
Free energy residue decomposition analysis for VIM-2-peptide complexes.

### Residue decomposition analysis

Further, we calculated the direct contribution of F residues to the Total B.F.E., which can be seen for all the AMPs in **Table 2**. Further, we observed that K repeats show consistent binding with VIM-2 in case of all peptides which have K repeats. These follow the trend that we reported previously in one of our studies **(Anurag Anand et al., 2024)**. The same has been mentioned in **Table 2**.

**Table 2.**
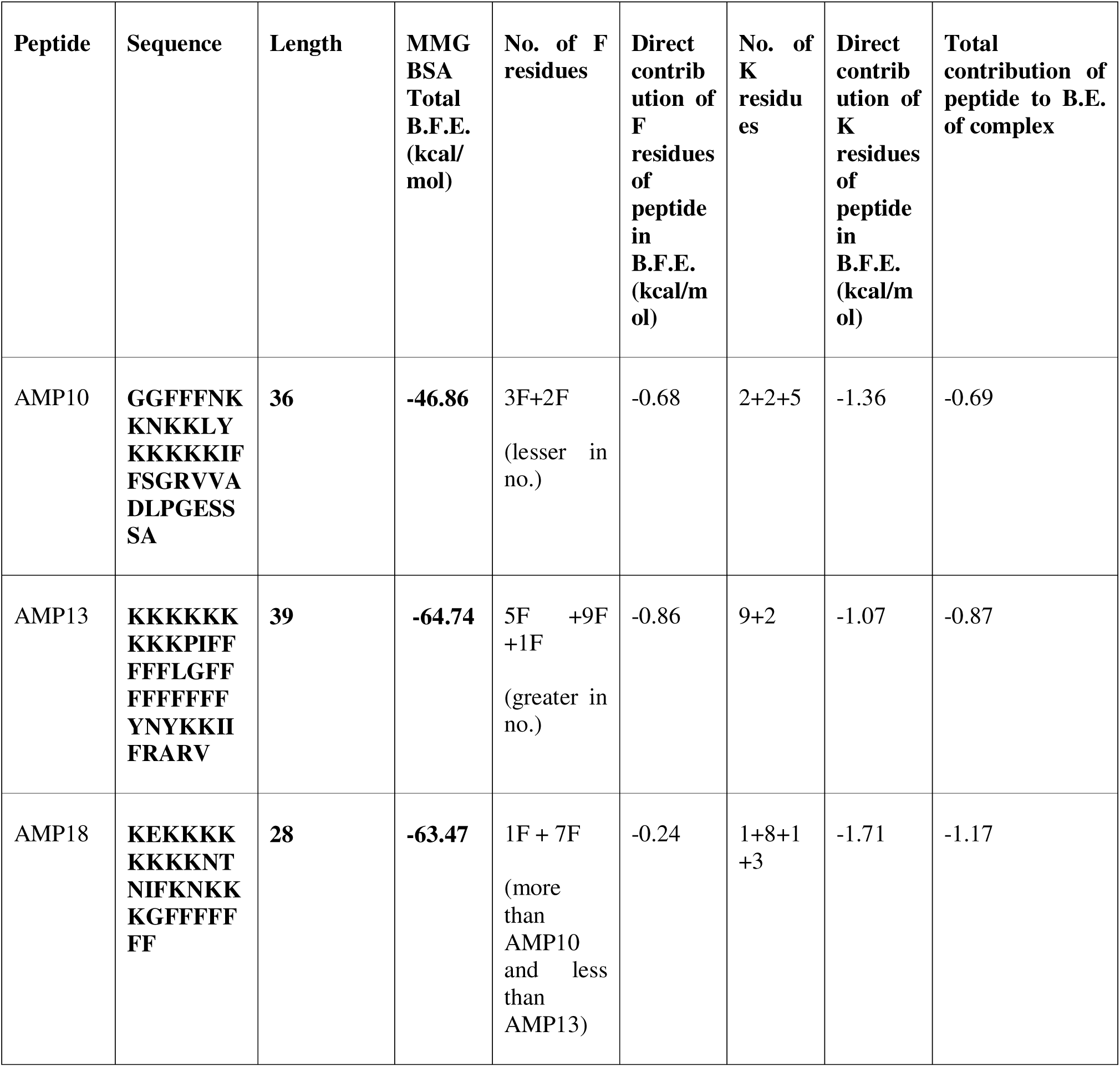
Contribution of F and K residues of peptide to the B.F.E. of the protein-peptide complexes.

Here, we observe that F repeats show overall negative B.E. in case of each peptide. However, F repeats alone are not sufficient for binding with VIM-2, as the peptides here are using other amino acids such as Asn, Ile, Lys and others as well to bind effectively with VIM-2.

**Table 3** shows contribution of all other residues to Total B.F.E. of the peptide-VIM-2 complex.

**Table 3.**
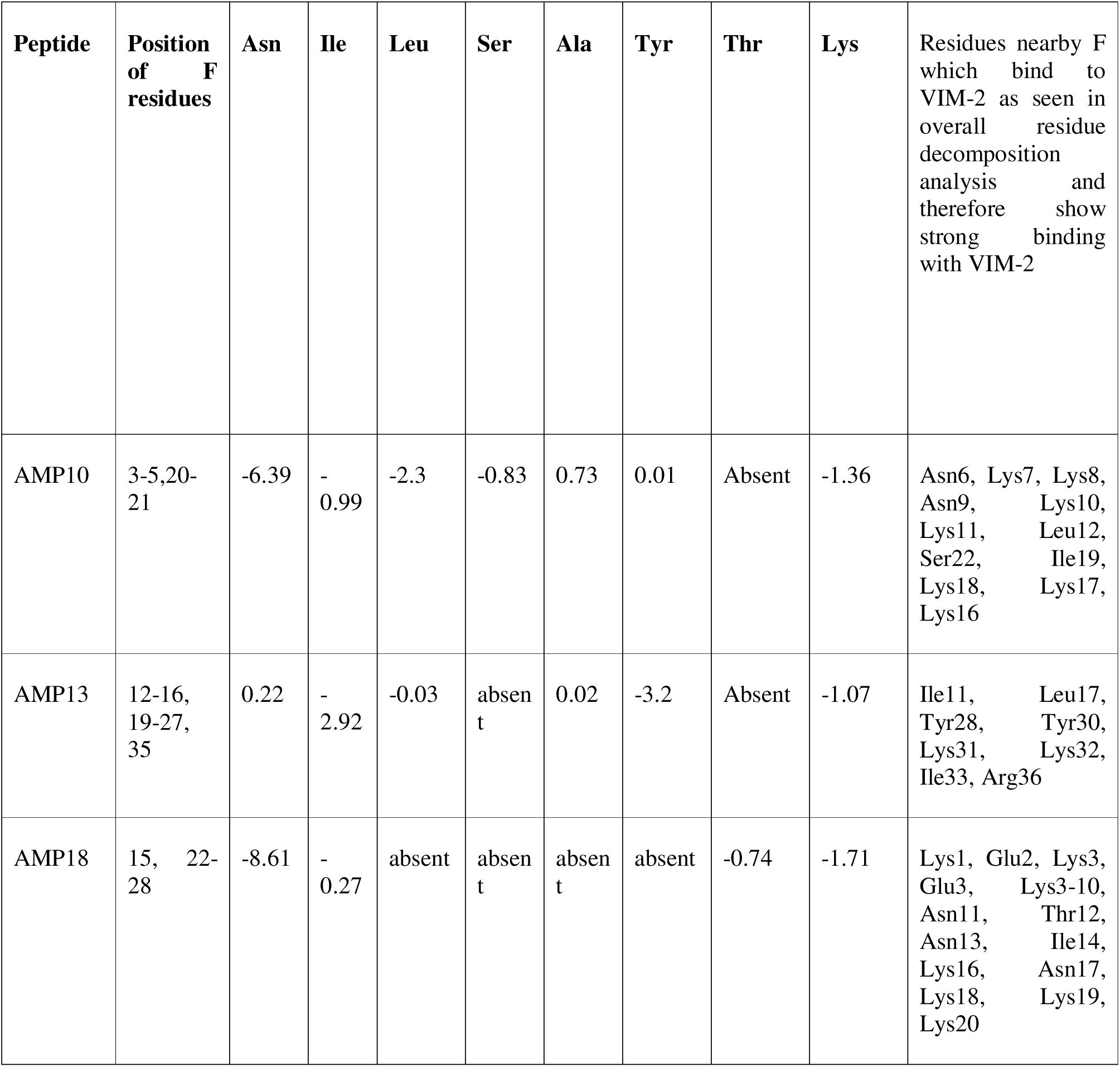
Contribution of residues other than F of peptide that bind to VIM to the B.F.E. of the protein-peptide complexes and their proximity to F.

A closer look at Table 5 shows that residues which lie closer to Phe show negative B.E., such as, in AMP10 we see that since Ala and Tyr are far away from Phe, they show positive B.E. but other residues likeAsn6 and Asn9 which lie adjacent to F residues lead to an overall negative B.E. for VIM-2. The details of all peptides showing dependence of contribution of residues to B.F.E. on their proximity to Phe residues can be clearly studied in **Table 3**.

**Table 4.** Average and standard deviation values of structure order parameters.

| Parameter | VIM2 | VIM2-PolyR | VIM2-AMP10 | VIM2-AMP13 | VIM2-AMP18 | VIM2-PolyF |
| --- | --- | --- | --- | --- | --- | --- |
| Rg | 1.330342679 ± 0.05183193 | 1.380713 ± 0.064147 | 1.455946 ± 0.063845 | 1.585726 ± 0.027804 | 1.472005 ± 0.04877 | 1.541891 ± 0.097729 |
| RM SD | 2.36E-01 ± 2.68E-02 | 2.92E-01 ± 3.00E-02 | 4.16E-01 ± 4.30E-02 | 3.96E-01 ± 4.71E-02 | 4.28E-01 ± 9.35E-02 | 3.18E-01 ± 2.57E-02 |
| RM SF | 0.121036957 ± 0.063378756 | 0.135104783 ± 0.062238381 | 0.158472609 ± 0.07340017 | 0.114891739 ± 0.047338277 | 0.139563043 ± 0.055789133 | 0.120265652 ± 0.063696844 |
| SASA | 106.6026202 ± 1.692827189 | 134.7570885 ± 2.737608195 | 131.2303038 ± 2.824286826 | 124.5507841 ± 3.516660358 | 132.6305102 ± 3.060753517 | 129.308965 ± 2.264631639 |

**Table 5.**
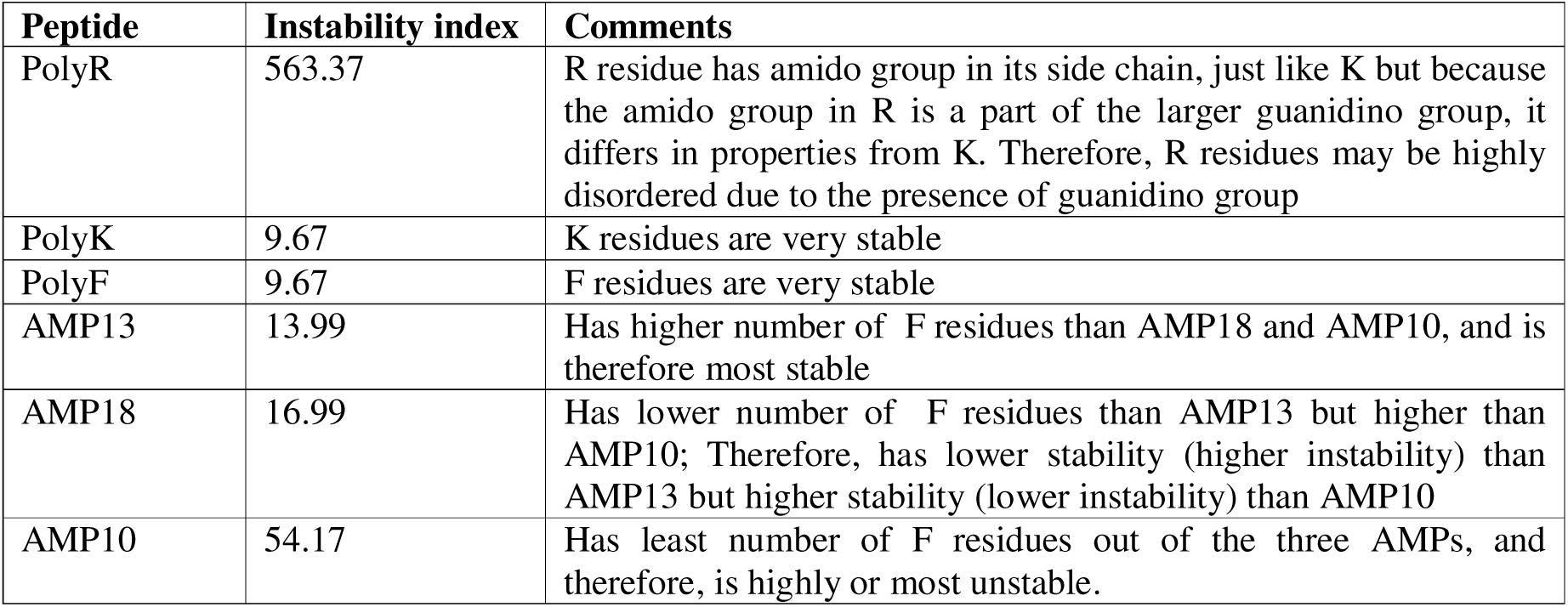
Instability index of unbound peptides.

| Peptide | Instability index | Comments |
| --- | --- | --- |
| PolyR | 563.37 | R residue has amido group in its side chain, just like K but because the amido group in R is a part of the larger guanidino group, it differs in properties from K. Therefore, R residues may be highly disordered due to the presence of guanidino group |
| PolyK | 9.67 | K residues are very stable |
| PolyF | 9.67 | F residues are very stable |
| AMP13 | 13.99 | Has higher number of F residues than AMP18 and AMP10, and is therefore most stable |
| AMP18 | 16.99 | Has lower number of F residues than AMP13 but higher than AMP10; Therefore, has lower stability (higher instability) than AMP13 but higher stability (lower instability) than AMP10 |
| AMP10 | 54.17 | Has least number of F residues out of the three AMPs, and therefore, is highly or most unstable. |

This means that although F residues themselves directly bind with VIM-2 only weakly, they help other residues which lie adjacent or near to them in very tight binding with VIM-2, such as, Lys in case of all 3 peptides (AMP10, 13 and 18), Asn in case of AMP10 and 18, and Tyr in case of AMP13.

Further insights from residue decomposition analysis include the top 10 residues that our peptides bind with. All our AMPs except PolyF bind more with negatively charged residues Asp and Glu (forming electrostatic bonds/bonds between negatively charged residues of the protein and positively charged residues like Lys of the peptides), whereas PolyF seems to mostly bind with residues other than Asp and Glu. **Figure 2** shows the binding energy of the top 10 binding residues for each of the protein-peptide complexes.

**Figure 2.**
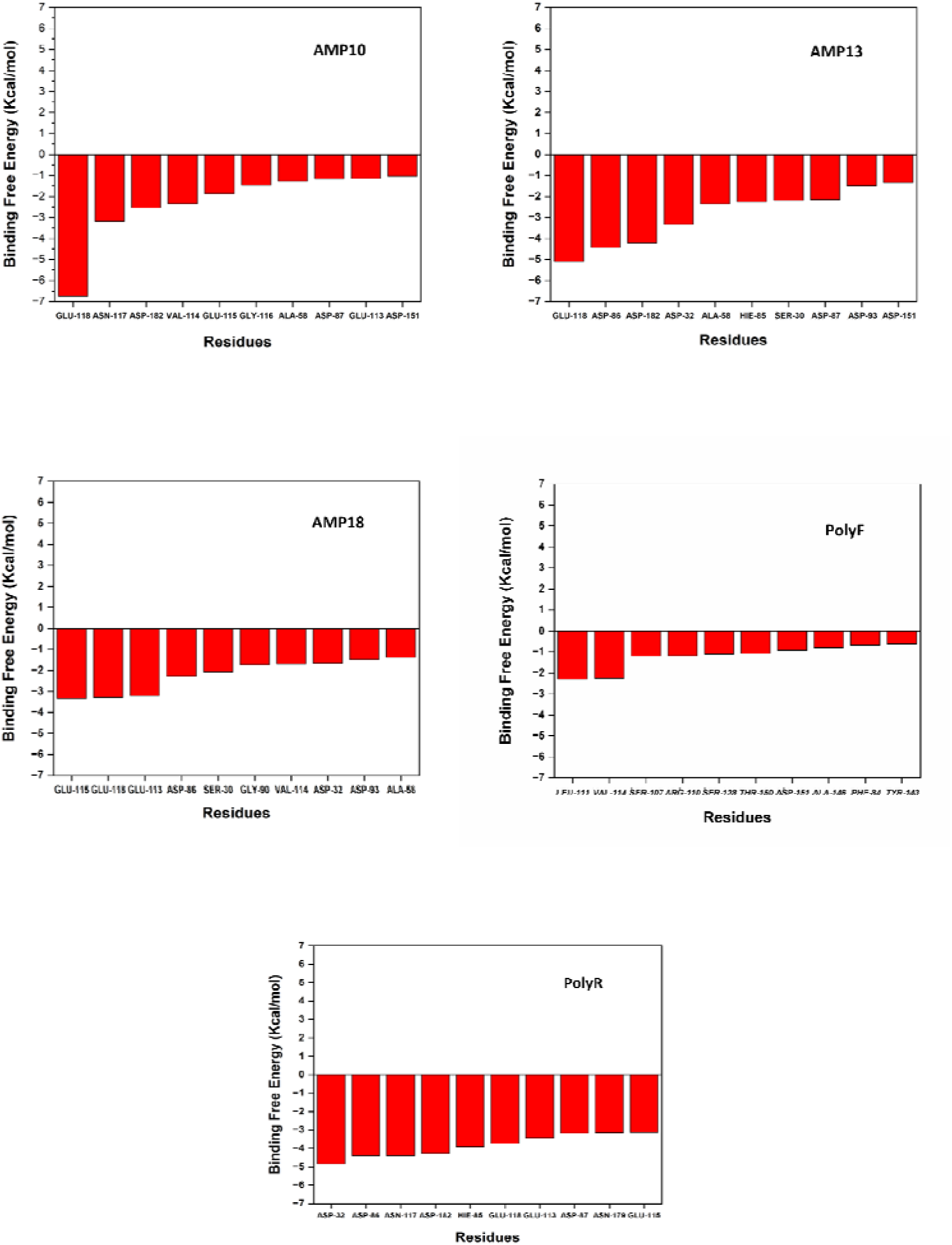
Top 10 residues of protein interacting with the peptides.

A deeper dive into the active site of the enzyme VIM-2 when bound to the peptides show that PolyR, followed by AMP13, AMP18 and AMP10 bind most effectively with the conserved active site residue Asp87 (corresponding to Asp120 of the original VIM-2 1KO3 structure; the structure was renumbered for its sequence while performing docking using HAWKDOCK, by default), whereas PolyF binds minimally with it (Anurag Anand et al., 2023). **Figure 3** shows the binding energy of the active site residues for each of the protein-peptide complexes.

**Figure 3.**
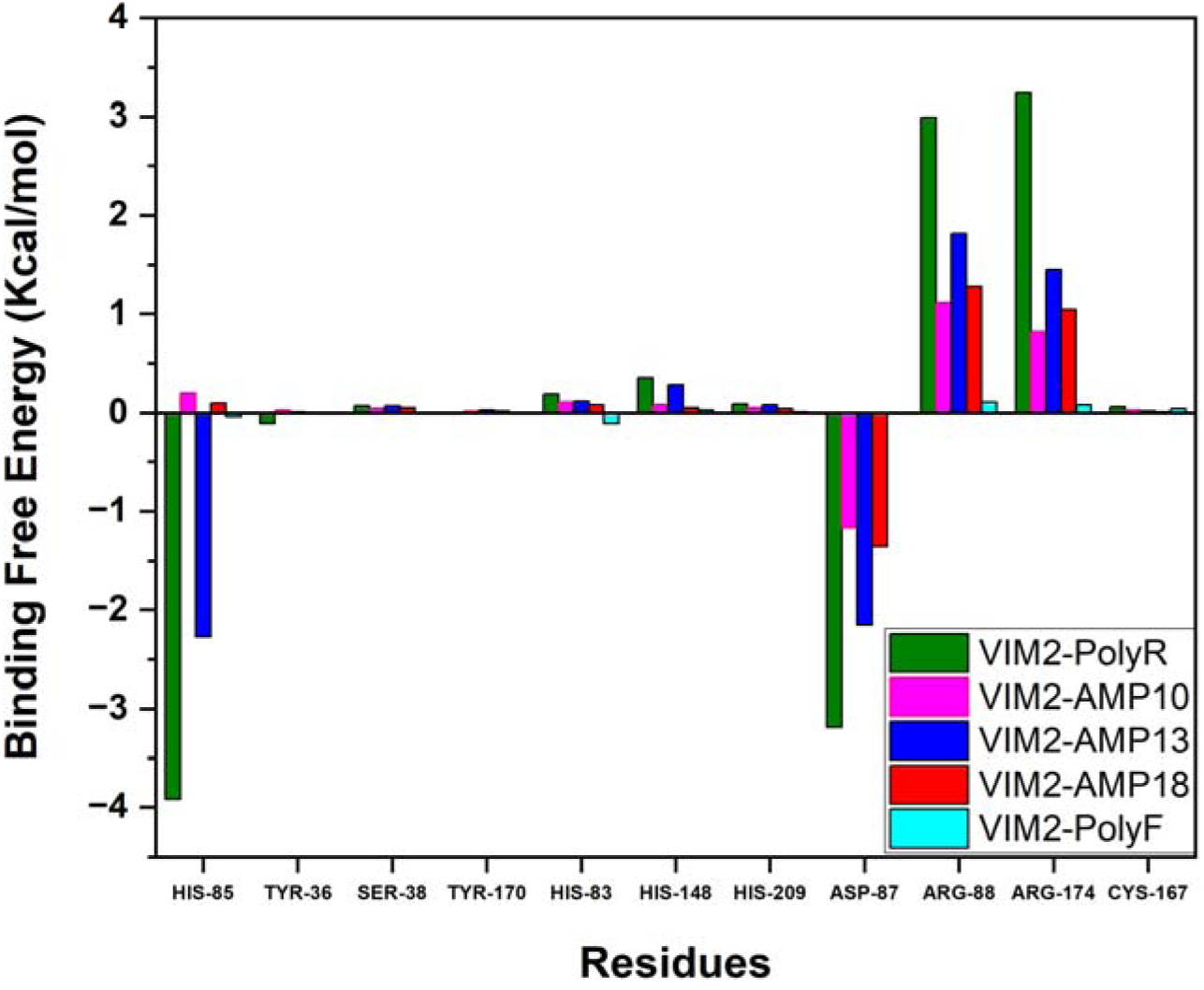
Interaction of active site residues of VIM-2 with the peptides.

### MD Simulation analysis results and interpretation

Figure 4-9 shows the plots from MD simulation analysis.

**Figure 4.**
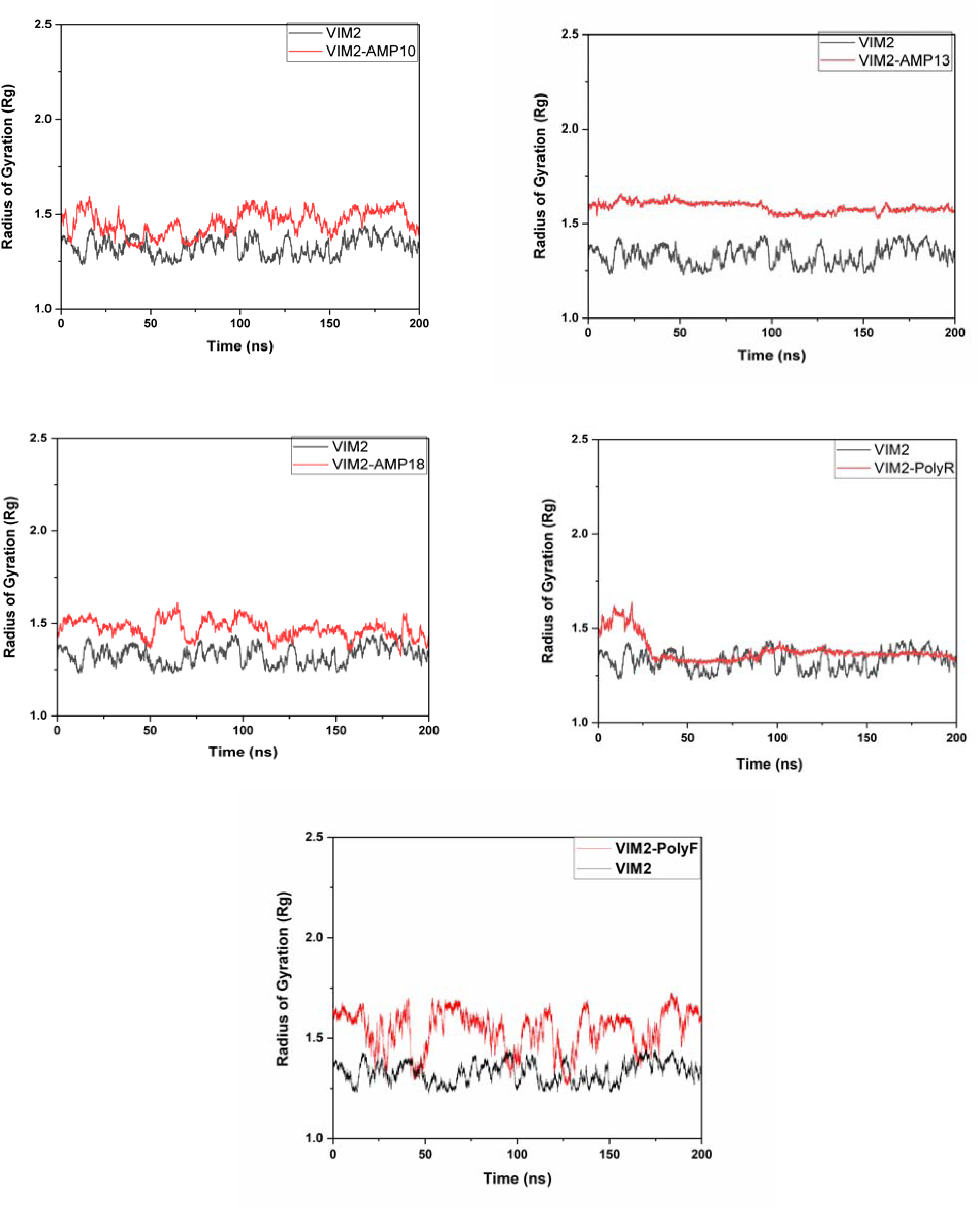
Plots showing change in Rg of VIM-2 upon binding with different peptides.

As expected from MMGBSA findings, we found that the protein’s Rg increases significantly after binding with PolyF, in addition to undergoing sharp fluctuations when the protein-peptide complex was simulated over a period of time, indicating unstable binding (Figure 4).

RMSD analysis revealed that all complexes undergo some fluctuations over the period of time (Figure 5). It is important to note that the protein when bound to AMP13 which has the highest no. of F residues out of our 3 main AMPs, shows constant SASA and Rg, as well as a low and constant RMSD. On the other hand, AMP18 and 10 having low no. of F residues, lead to higher and more fluctuating structure order parameters.

**Figure 5.**
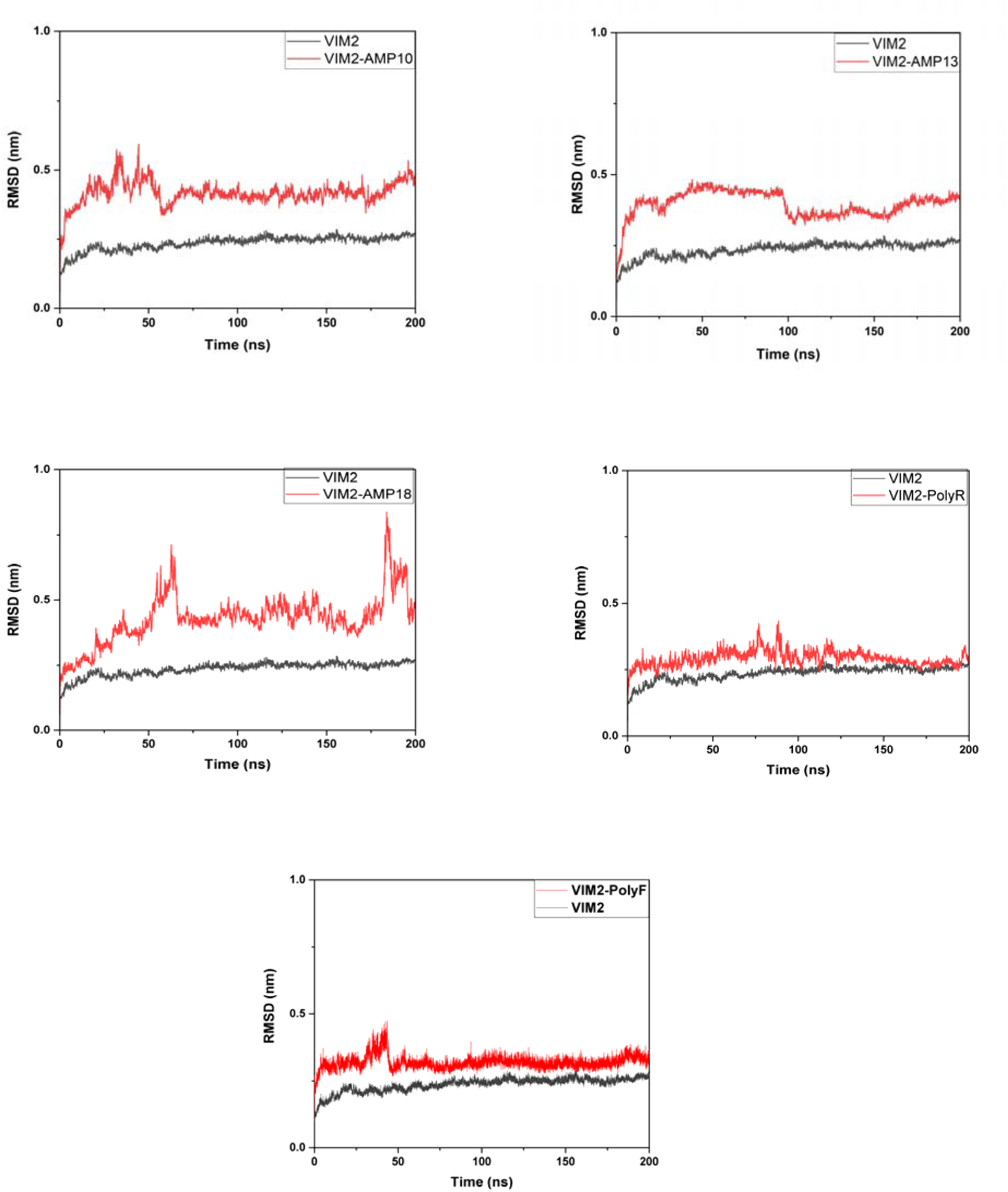
Plots showing change in RMSD of VIM-2 upon binding with different peptides.

RMSF showed very less variations for PolyF as well as for all other complexes. However, PolyF showed least variations in RMSF (Figure 6). SASA was found to be constant over the period of time for all complexes, including PolyF-VIM-2 (Figure 7). The high value of SASA was not our concern because peptides usually lead to an increase in SASA as per their length, therefore we focused only on the constant nature of SASA. Therefore, as expected, SASA was found to be constant over the period of time.

**Figure 6.**
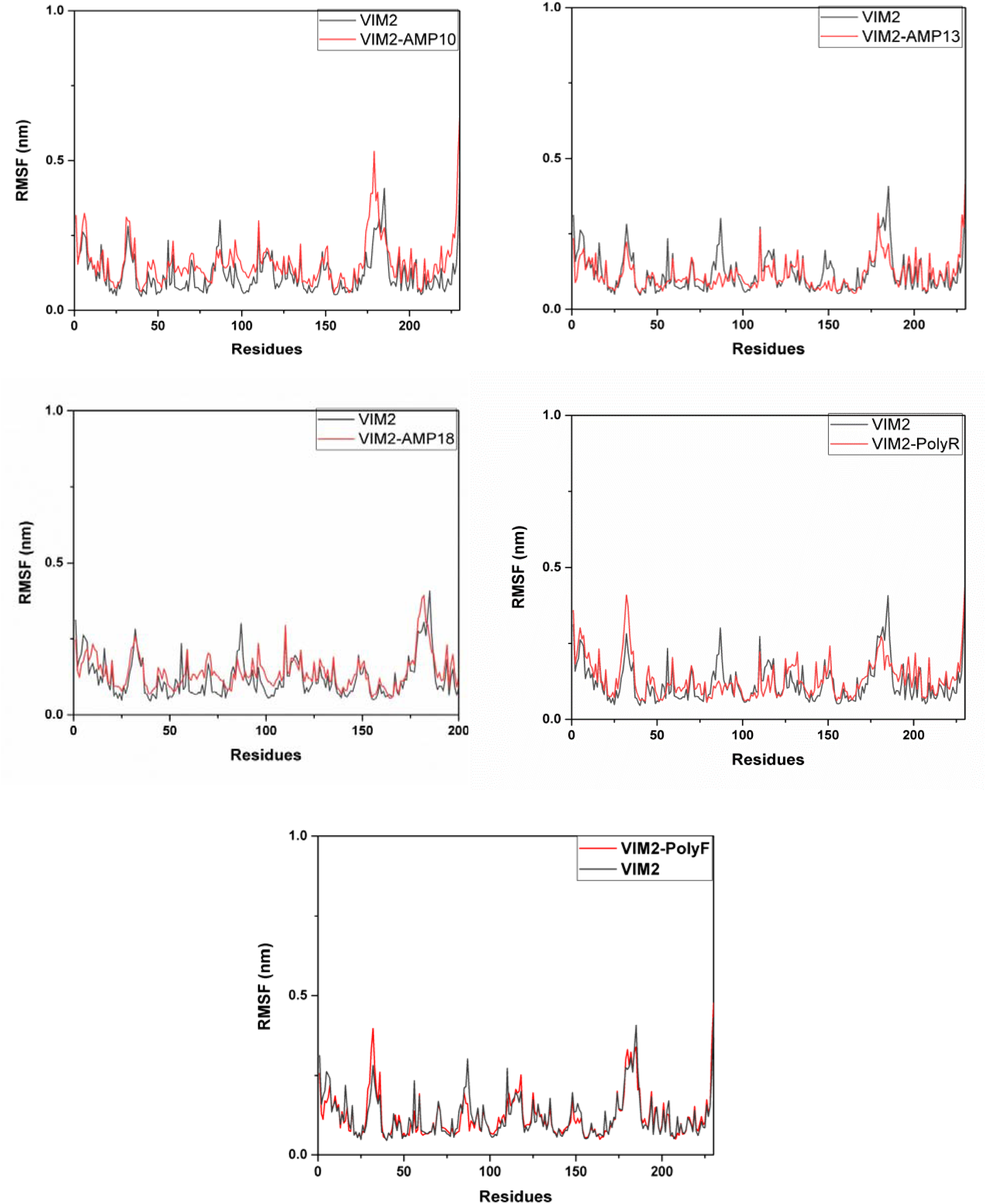
Plots showing change in RMSF of VIM-2 residues upon binding with different peptides.

**Figure 7.**
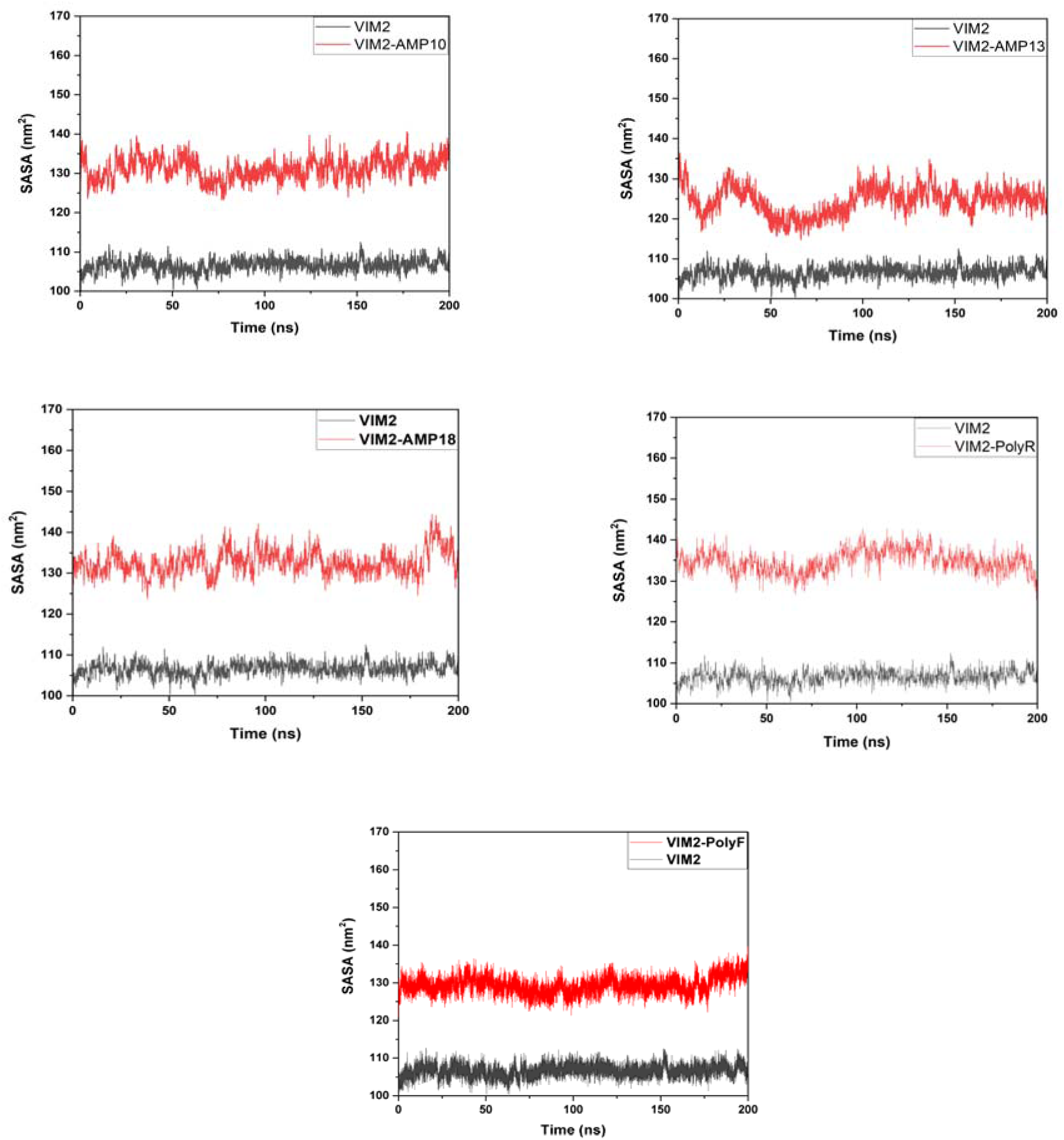
Plots showing change in SASA of VIM-2 residues upon binding with different peptides, depicting the change in surface area exposed to solvent.

Principal component analysis of PolyF-VIM-2 revealed that the complex doesn’t attain a stable conformation, as indicated by large dimensionality of PC1 and PC2 (**Figure 8** showing PC1 and PC2 alongside Gibb’s Free Energy). The PC1 range for PolyF-VIM-2 was found to be 8 to 10 folds higher than other complexes. FEL plots further indicate that, in case of PolyF-VIM-2, maximum number of residues do not leave the low energy state (stay in red region itself) (**Figure 8-9**). On the other hand, AMP13 shows maximum no. of residues in the blue region, thereby entering the lower energy state or leaving the higher energy state.

**Figure 8.**
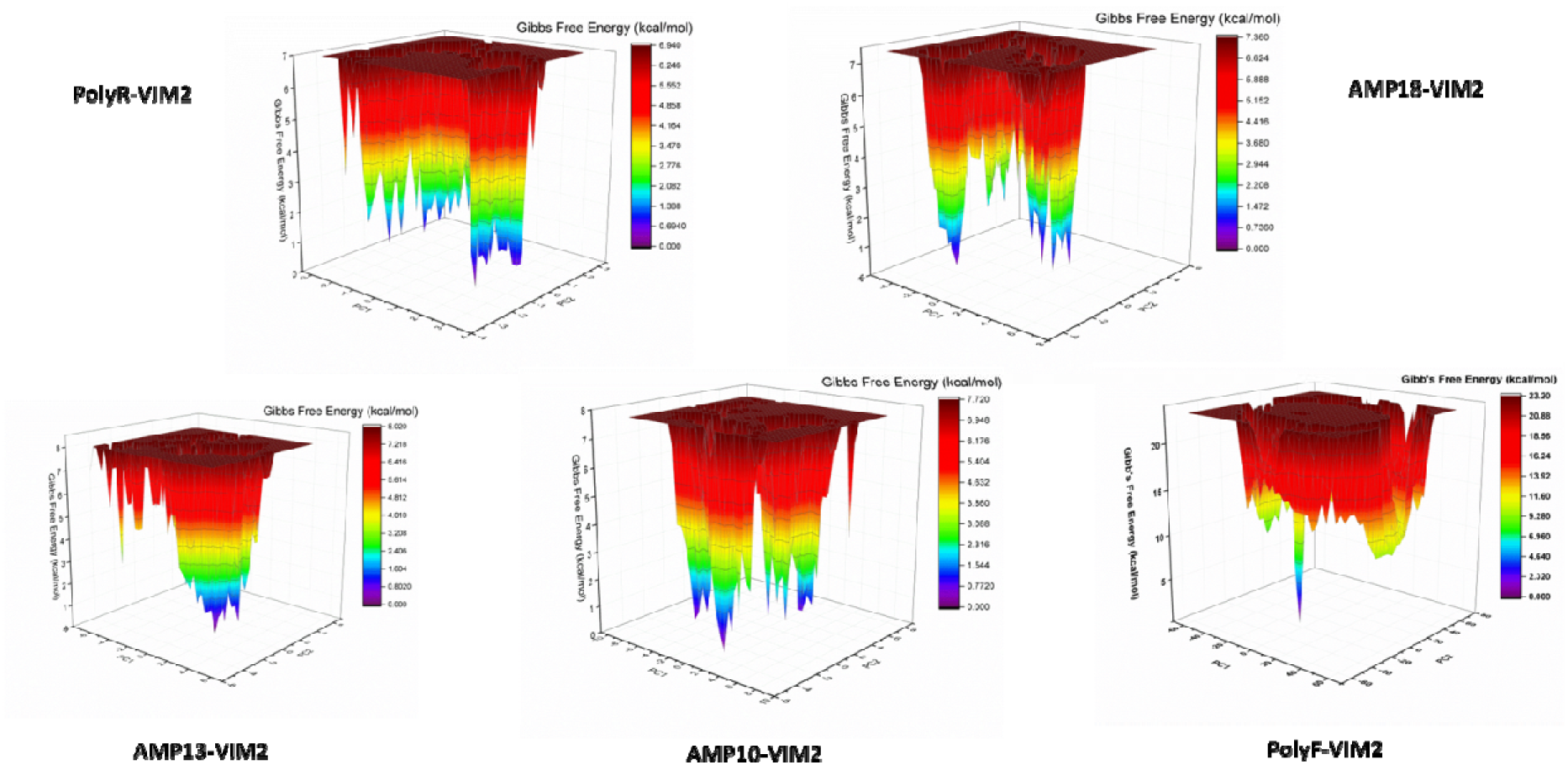
Free Energy Landscape of protein when bound to different peptides. The plots show: (i) change in Gibbs Free Energy of VIM-2 residues upon binding with different peptides, (ii) PC1 and PC2.

**Figure 9.**
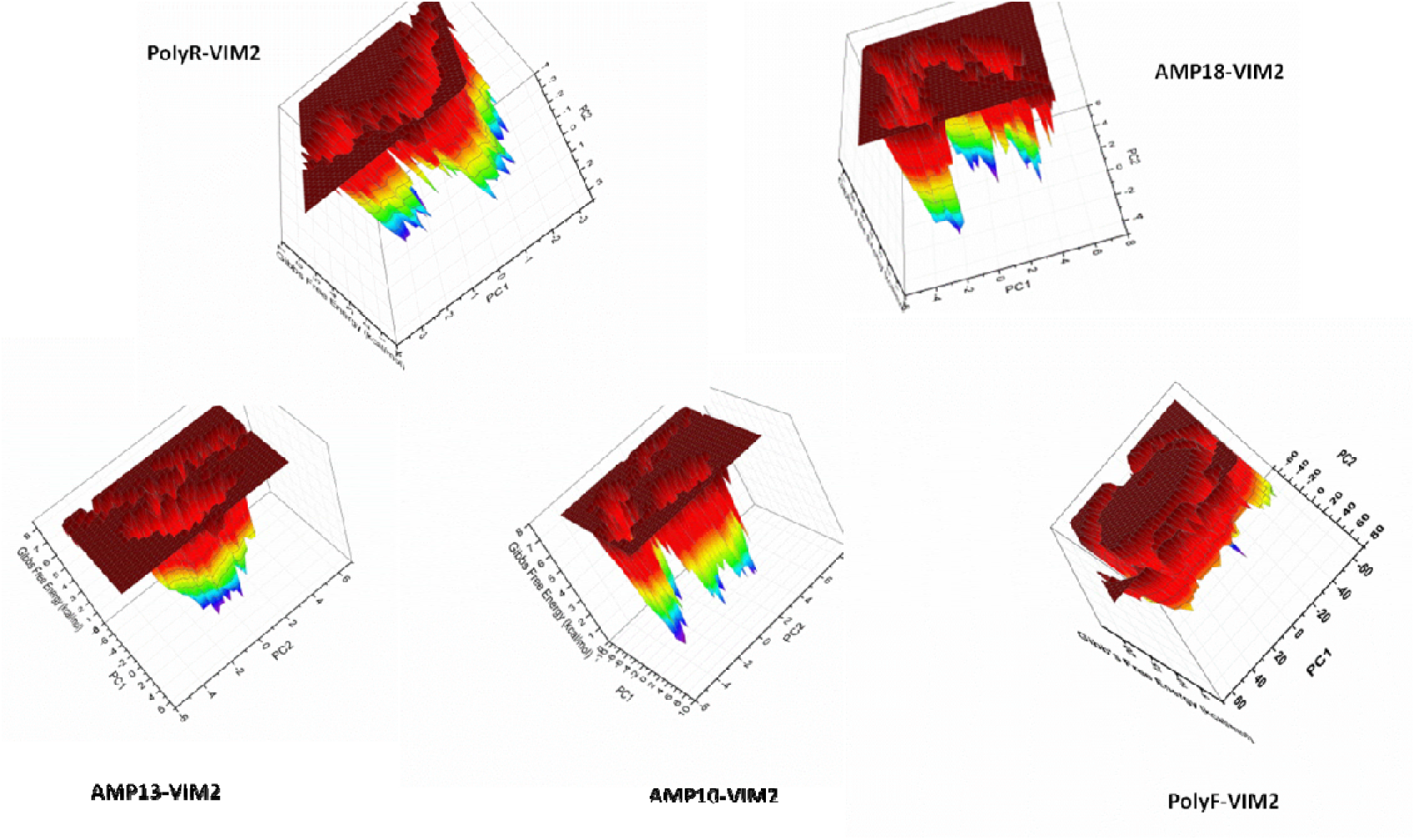
A view of protein-peptide complexes from FEL analysis showing the amount of residues leaving the red region or the higher energy state.

It is possible that PolyF which is not showing tight binding with VIM-2 (as indicated by least number of residues attaining low energy state as well as less negative B.F.E.) is loosely packed with it, but remains stably bound to it. Therefore, the loosely packed conformations do not strongly affect RMSD (which is protein-focused) or RMSF (residue-level fluctuation) which remain constant, but do affect the global shape of the complex, and hence Rg fluctuates.

The average values for RMSD, RMSF, SASA, and Rg alongside their standard deviation have been given in **Table 4**.

### PolyF shows high stability further strengthening our findings

In our study above, we have discussed that F repeats are showing weak binding with VIM-2. However, it plays a role in stabilizing the overall complex in presence of solvent. We further studied the stability of PolyF and other peptides in their unbound form (**Table 5**). We found that the instability index of PolyF is extremely low, just like K repeats. Secondly, AMP13 which has highest number of F residues is the most stable peptide with lowest instability index out of AMP10, 13 and 18. Similarly, AMP10 is most unstable due to least number of F residues out of the three AMPs. Therefore, despite the weak binding observed in molecular docking or initial binding free energy calculations, the solvent-mediated interactions and the intrinsic stability of the polyF chain contribute to maintaining the integrity of the peptide– protein complex during the simulation.

### Protein-peptide interaction analysis and active site proximity

We anticipated that if our F repeats bind near to (if not overlap) the Zn1/Zn2 binding sites, F repeats could influence the coordination of these ions. Therefore, we performed a detailed analysis for studying protein-peptide interactions between VIM-2 and different peptides at different time points of MD Simulation. In residue analysis we found that F residues do not bind to any of the active site residues except His 83 (corresponding to His 116 of original VIM-2 1KO3) and that too with very less negative binding energy. However, we found that F repeats bind to several residues lying adjacent to and near to important active site residues, mainly His (histidine residues) lying in the active site of VIM-2. In Figure 2, we can see that residues Tyr 143, Ala146, Thr150 and Asp151 show negative binding free energy values for F repeats. All of these residues lie near the active site residue H148 (corresponding to H196 of original VIM-2 1KO3), which is an active site residue. Further Phe84 also showed negative binding free energy when VIM is in complex with PolyF, which lies between two other active site residues His83 and His85 (corresponding to His 116 and His118 of original VIM-2 1KO3). Thus, F repeats hold the capacity to influence the coordination of Zn ions, thereby affecting the functioning of VIM-2 MBL. It is important to note that the functioning of Zn1 is dependent on His116, His118 and His196 residues as seen for the 1KO3 structure of VIM-2, which has VIM-2 MBL in its native reduced state. Therefore, the fact that F repeats could interfere with the functioning of VIM-2 in coordination with Zn ions is inevitable.

Additionally, we found F repeats to repeatedly interact with Arg110 (corresponding to Arg144 of original VIM-2 1KO3) and different serine residues at different time points of simulation (Figure 10).

**Figure 10.**
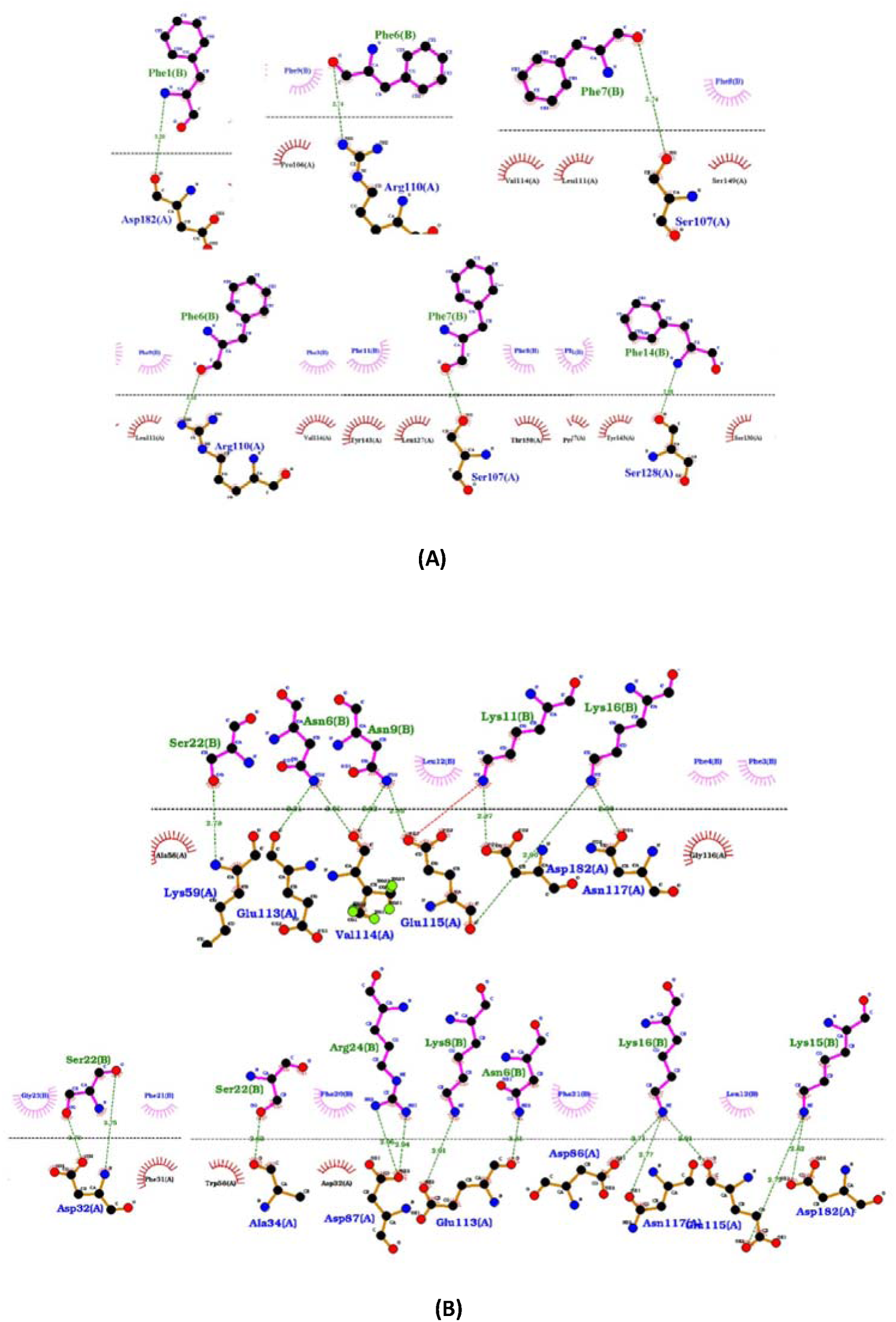
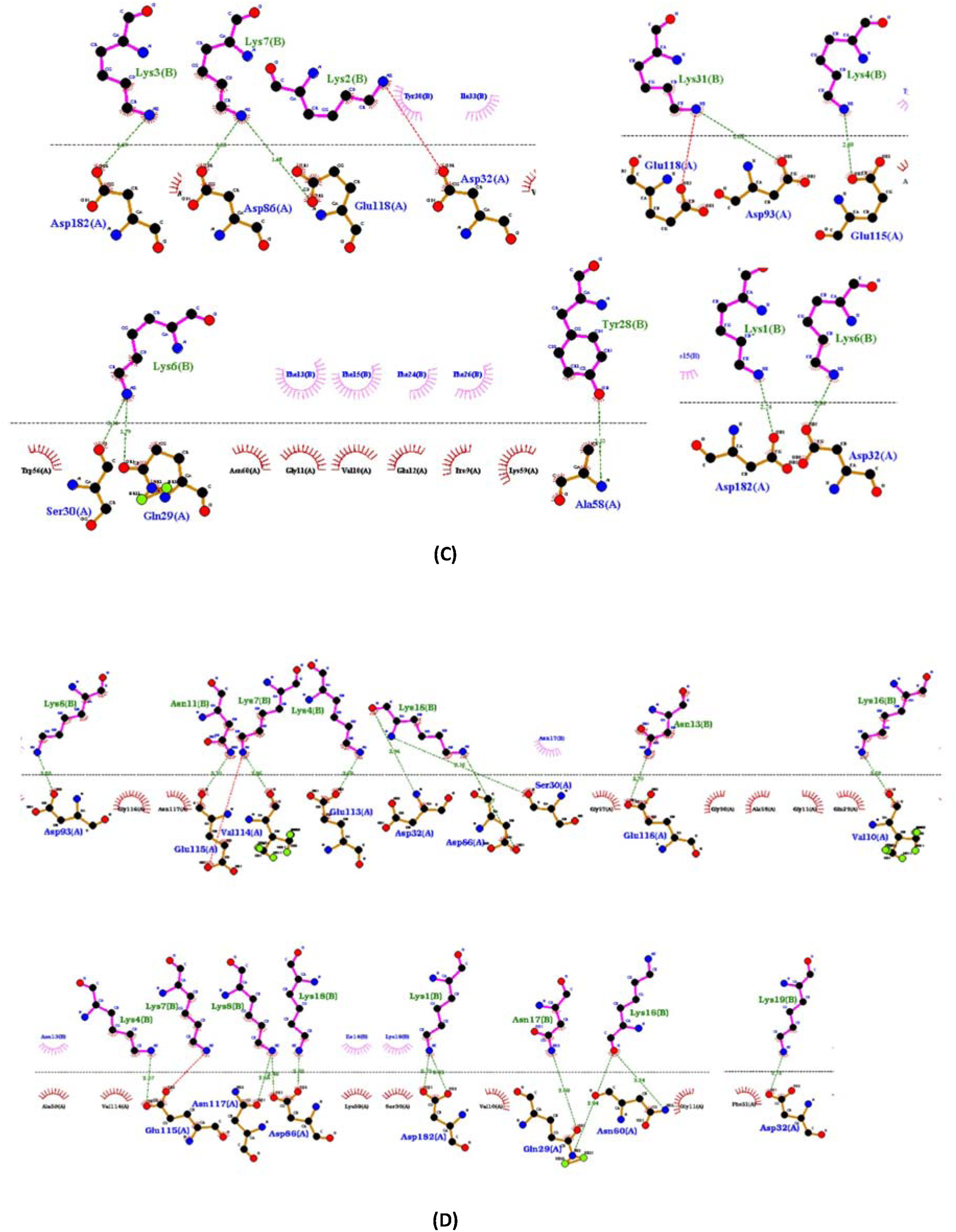
Ligplot analysis of VIM-2 when bound to peptides (A) PolyF or F repeats (B) AMP10 (C) AMP13 (D) AMP18. The figures show detailed protein-peptide interactions, mainly H-bonds and salt bridges, obtained from 0, 25, 50, 75, 100, 150 and 200 ns.

In case of AMP10, 13 and 18, we see that the positively charged Lys residues of the peptide bind with several Asp and Glu residues of the protein, as in both residue decomposition analysis as well as ligplot analysis (Figure 2**, Figure S1-S3)**. Similarly, in case of PolyR, we see that the positively charged Arg residues of the peptide bind with several Asp and Glu residues of the protein ((Figure 2). Additionally, as seen previously, AMP13 and PolyR bind very effectively with some active site residues (Figure 3). Further, in ligplots of AMP13, 18 and 10, we studied the proximity of binding residues of these peptides with the F repeats present in them. We then compared the results with overall decomposition analysis to get a clearer picture. We found that several residues of AMP13, 10 and 18, that lie around Phe, form H-bonds to VIM-2 (**Table 6**). The complete residue decomposition analysis for all the peptides when bound to VIM-2 can be found in **Table S5-S9.**

**Table 6.** Residues lying adjacent to and near Phenylalanine that bind with VIM-2.

| Peptide | Sequence | Length | Position of F residues | Residues nearby F that bind to VIM-2 as seen in ligplot analysis at different time points (0, 25, 50, 75, 100, 150 and 200 ns)<br>(H-bonds) | Residues nearby F that bind to VIM-2 as seen in overall residue decomposition analysis and free energy decomposition analysis<br>(inclusive of all types of bonds) | No. of F residues |
| --- | --- | --- | --- | --- | --- | --- |
| AMP10 | GGFFFNK<br>KNKKLYK<br>KKKKIFFS<br>GRVVADL<br>PGESSSA | 36 | 3-5,20-21 | Ser22, Arg 24 | Asn6, Lys7, Lys8, Asn9, Lys10, Lys11, Leu12, Ser22, Ile19, Lys18, Lys17, Lys16 | 3F+2F<br>(lesser in no.) |
| AMP13 | KKKKKKK<br>KKKPIFFF<br>FFLGFFFF<br>FFFFFYNY<br>KKIIFRAR<br>V | 39 | 12-16, 19-27, 35 | Tyr28 | Ile11, Leu17, Tyr28, Tyr30, Lys31, Lys32, Ile33, Arg36 | 5F +9F<br>+1F<br>(greater in no.) |
| AMP18 | KEKKKKK<br>KKKNTNI<br>FKNKKKG<br>FFFFFFF | 28 | 15, 22-28 | Lys16, Asn17, Lys18, Lys19 | Lys1, Glu2, Lys3, Glu3, Lys3-10, Asn11, Thr12, Asn13, Ile14, Lys16, Asn17, Lys18, Lys19, Lys20 | 1F + 7F<br>(more than AMP10 and less than AMP13) |

### Loop analysis shows conformational changes in loop 2 that affects substrate binding

Interestingly, while the F repeats do not appear to directly occupy the metal-binding sites, our simulations suggest that their presence influences the flexibility of loop 2, which is known to be critical for substrate binding (as discussed in relation to PhenylC3SH in previous sections). This indirect effect could play a role in maintaining the enzyme’s active conformation. While analyzing ligplots, we observed that PolyF forms H-bond to Asp182 of our VIM-2 structure (corresponding to Asp236 of original VIM-2 1KO3) (Figure 10). This residue is a part of loop 2, as mentioned in previous sections, which is crucial for substrate binding. We confirmed that the residues forming loop 2 of VIM-2 undergo a conformational change when bound to PolyF by visualizing the complex from MD Simulation. The same can be visualized in **Figure 11A**. Add it was seen that in AMP10, AMP13 and AMP18, the residues lying adjacent or near to F repeats show tight binding with loop1 and loop 2 of VIM-2 thereby affecting their conformations and even destabilizing them. The same can be visualized in **Figure 11B**. It is to note that in case of AMP10-VIM-2 the bonds formed with loops include: 1) Ser22 forming bonds with Asp32 and Ala34 of loop 1 (corresponding to Asp62 and Ala64 of 1KO3), and 2) Lys15 and Lys16 of the peptide form bond with Asp182 of VIM-2 (Asp236 of 1KO3). Similarly, Lys 18 and Lys19 of AMP10 form bonds with Asp32, whereas Lys1 forms bonds with Asp182. In AMP13, Lys2 and Lys6 form bonds with Asp32 whereas Lys1 and 3 form bond with Asp182. Therefore, Asp32 (or Asp62 originally) of loop1 and Asp182 (or Asp236 originally) of VIM-2 are the residues that interact with peptides containing KF repeats. On the other hand, PolyF only interacts directly with Asp182 (or originally Asp236) of loop1.

**Figure 11.**
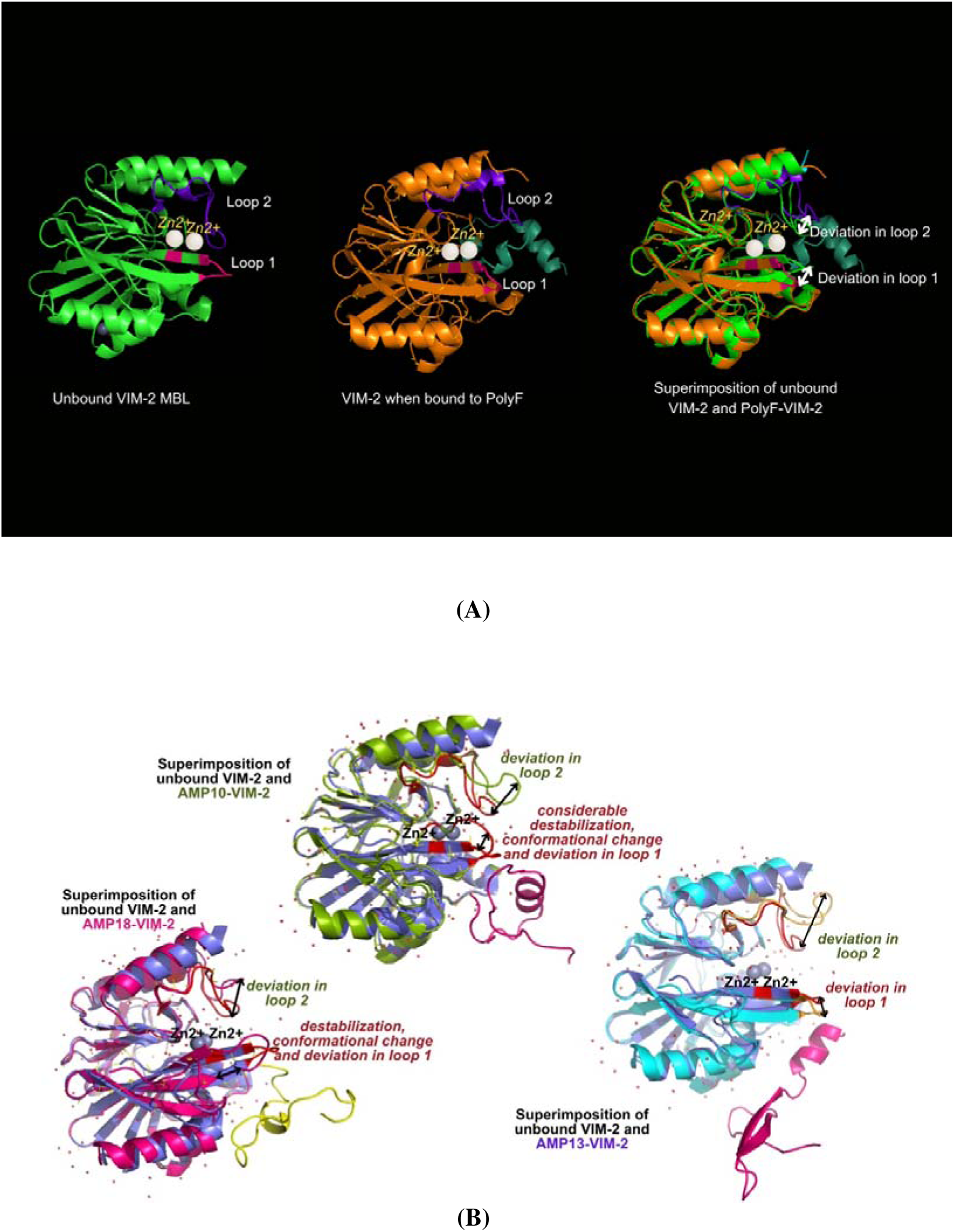
Analysis of conformational changes undergone by loops of VIM-2 (critical for substrate binding) when F repeats containing peptides bind to VIM-2. The study shows that conformational changes and destabilization effect is produced by such AMPs in VIM-2. (A) Changes brought about by PolyF (B) Conformational changes produced due to binding of AMP10, AMP13, and AMP18, and destabilization effect produced by AMP10 and AMP18 by destabilizing the beta-sheet and converting it into coil.

**Figure 12.**
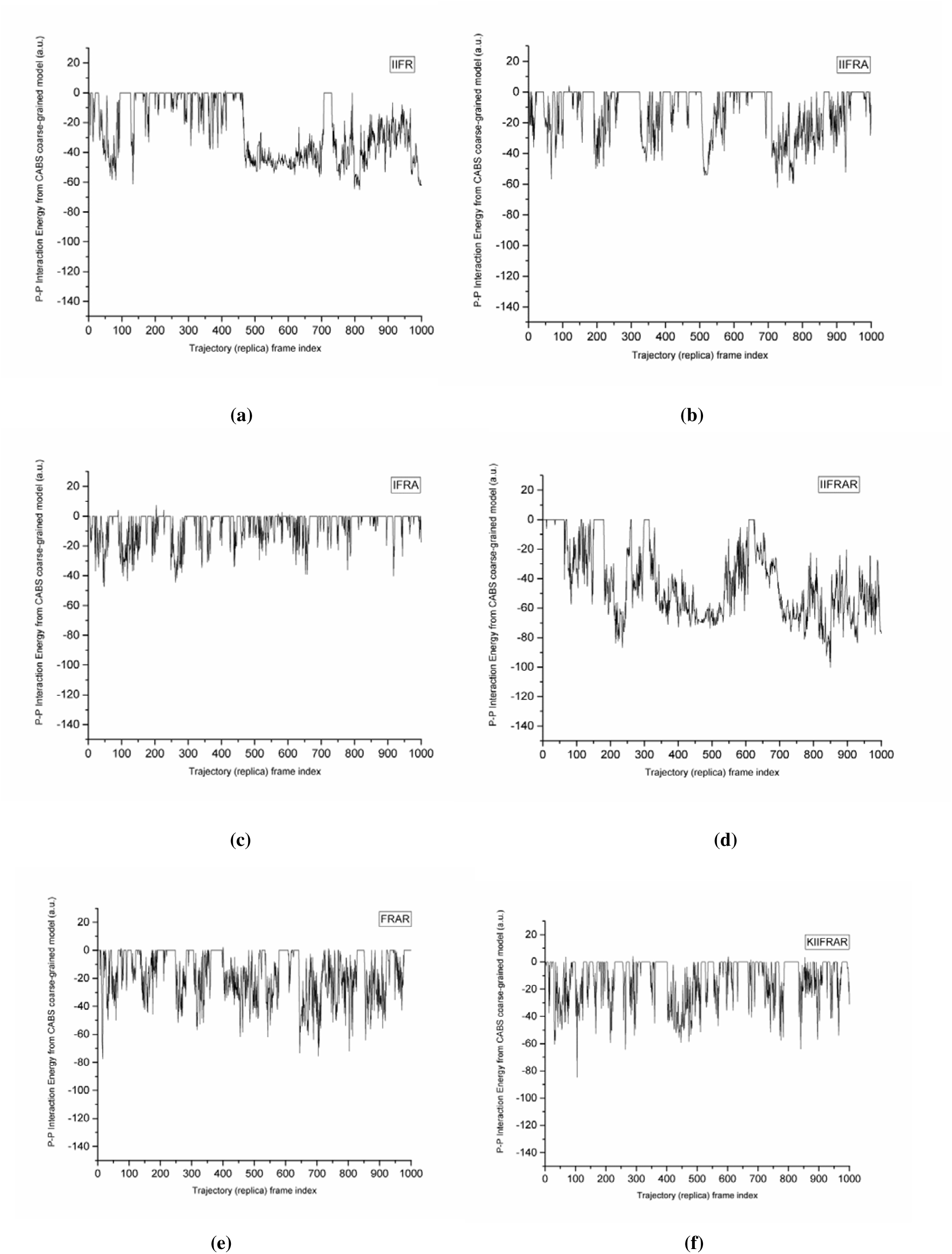
CABSDOCK trajectory of the phenylalanine-containing analogs belonging to set 2. When A was added to IIFR to form IIFRA, the peptide binding decreased noticeably, indicating that A interferes with binding of peptide with VIM-2 when added to C-terminal carrying R. Thus, A is interrupting the binding of R residue with VIM-2. When an Ile residue was removed from the N-terminal of IIFRA to form IFRA, peptide binding further decreased showing the impact of Ile residues on peptide binding with VIM-2. When comparing IIFR with IIFRAR, it is seen that IIFRAR shows highly enhanced binding with VIM-2 because of the addition of the AR moiety, which has a neutral residue towards the N-terminal whereas a charged residue on the C-terminal. When an R residue was added to IIFRA to form IIFRAR, the binding improved significantly, showing the necessity of R residue on C-terminal of peptide in order to bind with VIM-2. Again, when both Ile were removed from the N-terminal from IIFRAR (to form FRAR), the binding decreased noticeably ensuring the need for Ile residues for binding with VIM-2. To check having a positively charged residue like R or K on each of the terminals (both N and C) leads to good binding with VIM-2 or not, we added K to IIFRAR to form KIIFRAR, and found that the binding became insignificant as compared to the former. Thus, having positively charged residues on both terminals of the peptide could decrease binding with VIM-2. Thus, we anticipate that having one terminal as positively charged (moainly C-terminal) while other as neutral or hydrophobic (N-terminal) favours peptide binding with the VIM-2 MBL.

### Evaluating the library of amino acid repeats

To assess the binding efficiency of peptide repeats with the metallo-β-lactamase VIM-2, we employed a combination of flexible and rigid docking strategies, followed by binding free energy estimation. Initial flexible docking using CABSdock revealed that the 12-mer variants generally exhibited stronger interaction energies compared to their 6-mer or smaller counterparts. Among them, F12 showed the most favorable average interaction energy for Model 1 (−92.57 kcal/mol), followed closely by R12 (−59.39 kcal/mol) and K12 (−58.80 kcal/mol), indicating enhanced flexibility-driven binding potential for longer peptides. Rigid-body docking performed using HawkDock demonstrated a consistent trend, where the best docking scores were obtained for R12 (−5950.64), R6 (−4460.01), and K12 (−4216.93), suggesting strong shape complementarity and electrostatic interactions between these peptides and the VIM-2 active site. Further refinement using MM/GBSA free energy calculations identified R12 as the top binder with the most negative ΔG value (−107.81 kcal/mol), followed by K12 (−78.55 kcal/mol) and R6 (−76.29 kcal/mol). The consistently superior performance of arginine- and lysine-rich peptides, particularly in their 12-mer forms, highlights the critical role of positive charge and peptide length in enhancing binding affinity toward the negatively charged VIM-2 surface. In contrast, phenylalanine-rich peptides (F6 and F12) showed comparatively weaker binding in Hawkdock scores for docking and MMGBSA, likely due to the absence of favorable electrostatic interactions, but most negatives scores in CABSDOCK.

In our comparative analysis, CABSdock favored the F12 peptide due to its preference for hydrophobic contacts and conformational flexibility, whereas HawkDock/MMGBSA prioritized peptides rich in positively charged residues, particularly arginine (R) and lysine (K), because the negatively charged surface of VIM-2 promotes strong electrostatic and hydrogen-bond interactions. Given that electrostatic interactions generally dominate over hydrophobic packing in aqueous environments, the HawkDock/MMGBSA ranking is considered more biologically relevant. Integrating both docking and MMGBSA findings, the overall binding efficiency order is R12 > K12 > R6 > F12 > K6 > F6, with significantly weaker binding for shorter peptides. On a per-residue basis, arginine demonstrated the highest binding potential, followed by lysine and then phenylalanine, with the effect being most pronounced in the longer 12-mer peptides.

The results for the same have been summarized in **Table 7**.

**Table 7.**
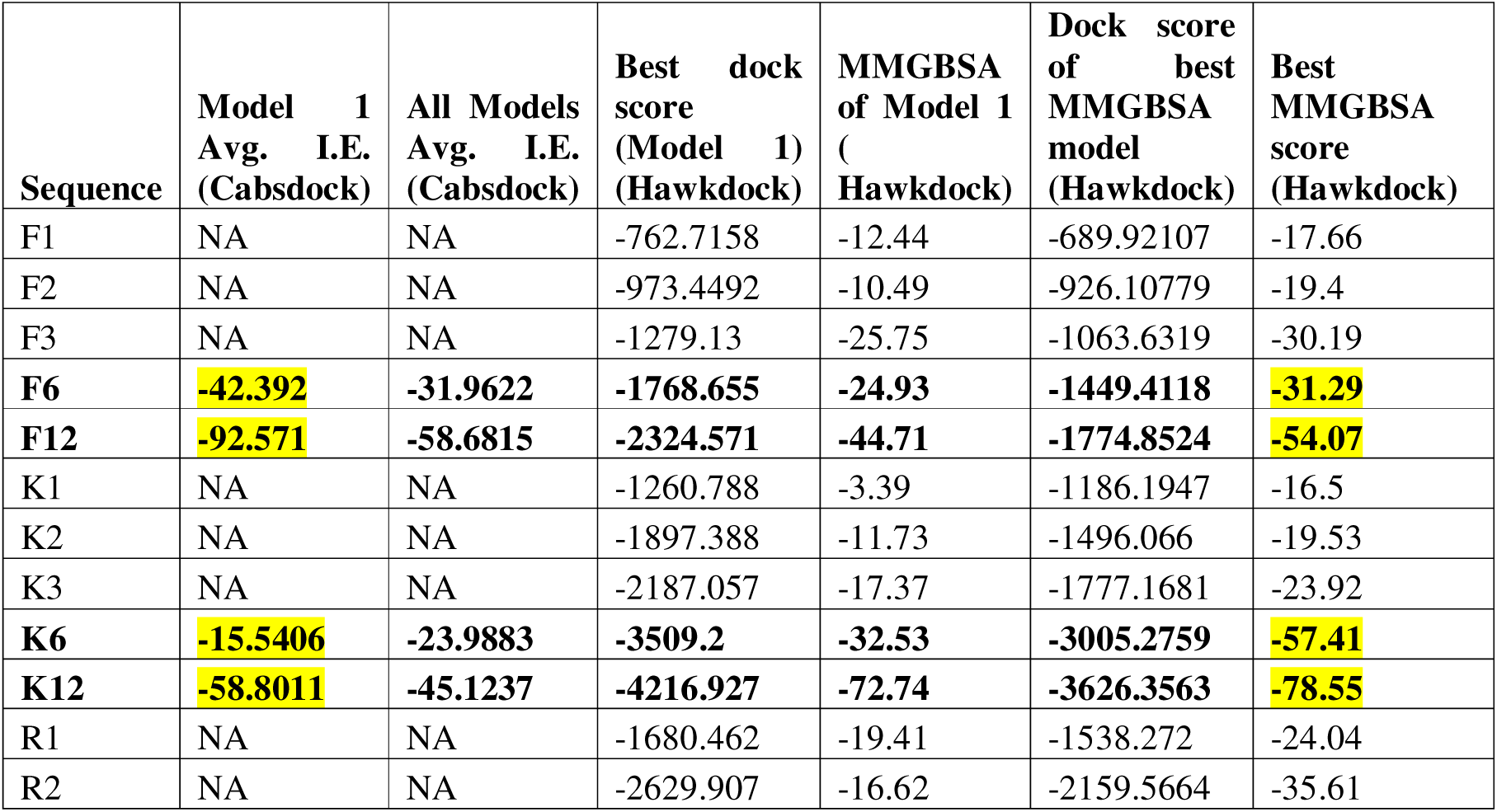
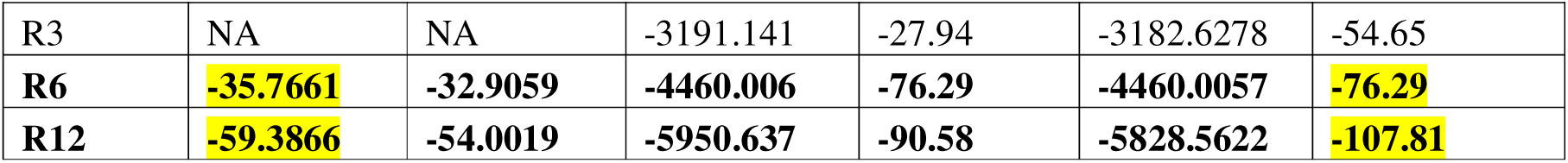
Docking and MMGBSA results for different amino acid repeats.

### Major findings from the gut-derived peptides

AMP13, having the highest no. of F residues, shows a lower and constant protein RMSD and SASA, and the lowest binding free energy (Total B.F.E.) when bound to VIM-2. AMP18, having lower no. of F residues as compared to AMP13, leads to high SASA, high and fluctuating RMSD, and comparatively high B.F.E. as compared to AMP13. AMP10, having the lowest no. of F residues leads to high SASA, high and fluctuating RMSD, alongside highest B.F.E. Finally, PolyF lacks positively charged residues, therefore do not bind effectively with VIM-2, thereby showing low binding affinity as indicated by the high binding free energy. However, F residues contribute to the stability of the protein-peptide complex in aqueous environment as indicated by low polar solvation energy. Our study also suggests a potentially distinct mechanism of stabilization, where the F repeats contribute to the overall structural integrity, by directly (by binding with His 116 of active site) as well as indirectly (by binding with several residues lying near and adjacent to several active site residues) optimizing the active site for interactions with other residues of a peptide like K such as in AMP13, AMP10 and AMP18. The low polar solvation energy of the F-repeats highlights the role of F repeats in stabilizing the protein-peptide complex in presence of a solvent. F repeats also show direct binding to loop 2 of VIM-2 but most importantly helps other residues of the peptide in binding with both the functional loops (loop 1 and 2, which help in functioning of VIM-2 at the active site), leading to their destabilization.

The major findings have been summarized in **Table 8** below.

**Table 8.** Overall major findings from docking, MMGBSA, free energy decomposition analysis, residue analysis and MD simulation.

| Parameter | AMP10 | AMP13 | AMP18 | PolyF |
| --- | --- | --- | --- | --- |
| RMSD | High but constant | Low and constant | High but constant | Low and constant |
| Rg | Fluctuating | Most constant | Relatively less constant | Highly fluctuating |
| RMSF | Slight fluctuation | Slight fluctuation | Slight fluctuation | Slightest/ least fluctuation; maximum overlapping with unbound protein |
| PC1 and PC2 variation | Less compact dimensions | More compact dimensions | More compact dimensions | Very high PC variation |

| Total Binding Free Energy (kcal/mol) | Least negative out of AMP13, 10 and 18 | Most negative | Less negative than AMP13 but more negative than AMP10 | Least negative |
| --- | --- | --- | --- | --- |
| F repeats and their contribution | Lesser in number; less compact structure | Very high in no.; most compact structure | Lesser in number than AMP13 but more than AMP10; compactness follows the trend | Stabilizing the peptide-VIM-2 complex in presence of solvent; direct binding to active site (weak); direct binding to loop 2 which affects substrate binding; helping nearby residues to bind tightly with the protein and affect active site binding as well as functioning of both loop 1 and loop 2. |
| K repeats and their contribution | Less in number; less no. of electrostatic interactions | Very high in no.; very high amount of electrostatic interactions | Less than AMP13 but more than AMP10; electrostatic interactions follow the pattern | None; Very few/insignificant number of electrostatic interactions |

### Derivation and distribution of Library I, II and III analogs

We found 22 high binding segments (HBS) from the parent AMPs, which formed library I of our study. In Library I, out of the 22 analogs derived from the parents, only 5 consisted of the F tripeptide. On the other hand, the number of sequences having F dipeptide and monopeptide were 8 and 9, respectively. The library I sequences can be understood in detail in **Table S10.** Further, 17 analogs were derived from the above analogs to create library II. Out of these, 10 were found to have F monopeptide whereas 4 were found to have F dipeptide. Only 3 were found to possess F monopeptide. The library II sequences can be understood in detail in **Table S11.**

This distribution indicates a divergence in the role of F-containing motifs between the parent AMPs and the high binding segments (HBS) enriched for interaction with VIM-2 MBL. While the parent AMPs contain long stretches of phenylalanine—such as FFFFFFFFFFFF— these regions are not retained in the majority of the VIM-2-binding HBS. Instead, the HBS predominantly feature shorter F motifs (mono- and dipeptides), suggesting that extended F-rich sequences may not be optimal for specific binding to the VIM-2 active site or surface.

The long F stretches in the parent peptides may serve other structural or functional roles, such as facilitating stabilization of the complex in solvent (as described above) or contributing to overall hydrophobicity. However, for targeted binding to VIM-2, the selection process appears to favor more moderate hydrophobic contributions from phenylalanine, likely to avoid steric clashes or nonspecific hydrophobic interactions that could interfere with precise binding. This shift highlights a functional distinction: while long F-rich domains may enhance general antimicrobial properties, fine-tuned interaction with VIM-2 requires more selective and spatially controlled incorporation of F residues.

Further, new analogs were designed to study the impact of F residues further, mainly in the presence of K and N this time because our binding sequences were found to be dominated by the presence of these residues. Additionally, few sequences were also designed to study the transition from one analog to another. These intermediate sequences alongwith the K, N and F containing sequences were included in our library III, which can also be called as the custom library of this study. The custom library was further extended based on our insights from the overall analysis of the existing analogs at any given point of time. The final custom library can be studied in **Table S12.**

### Formation of similarity based sets and overall evaluation of analogs

Once the analogs were created they were collated and distributed into 10 new sets based on the similarity in the sequences, known as similarity based sets (SBS). The list of composition of each of the SBS can be seen in **Table S13.**

**Table 9** summarizes findings from all sets of SBS analogs. These findings can be understood in detail below.

**Table 9.**
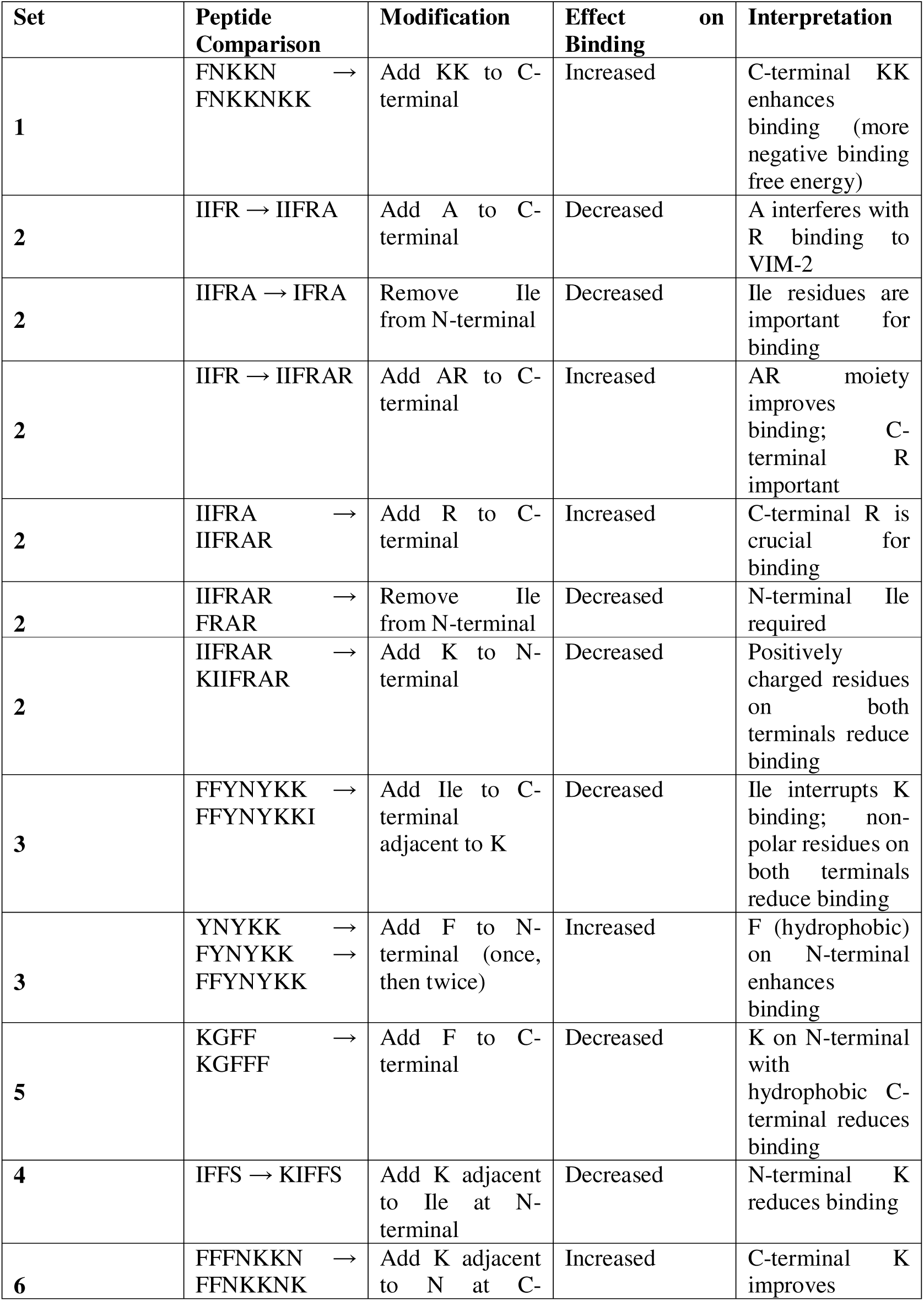
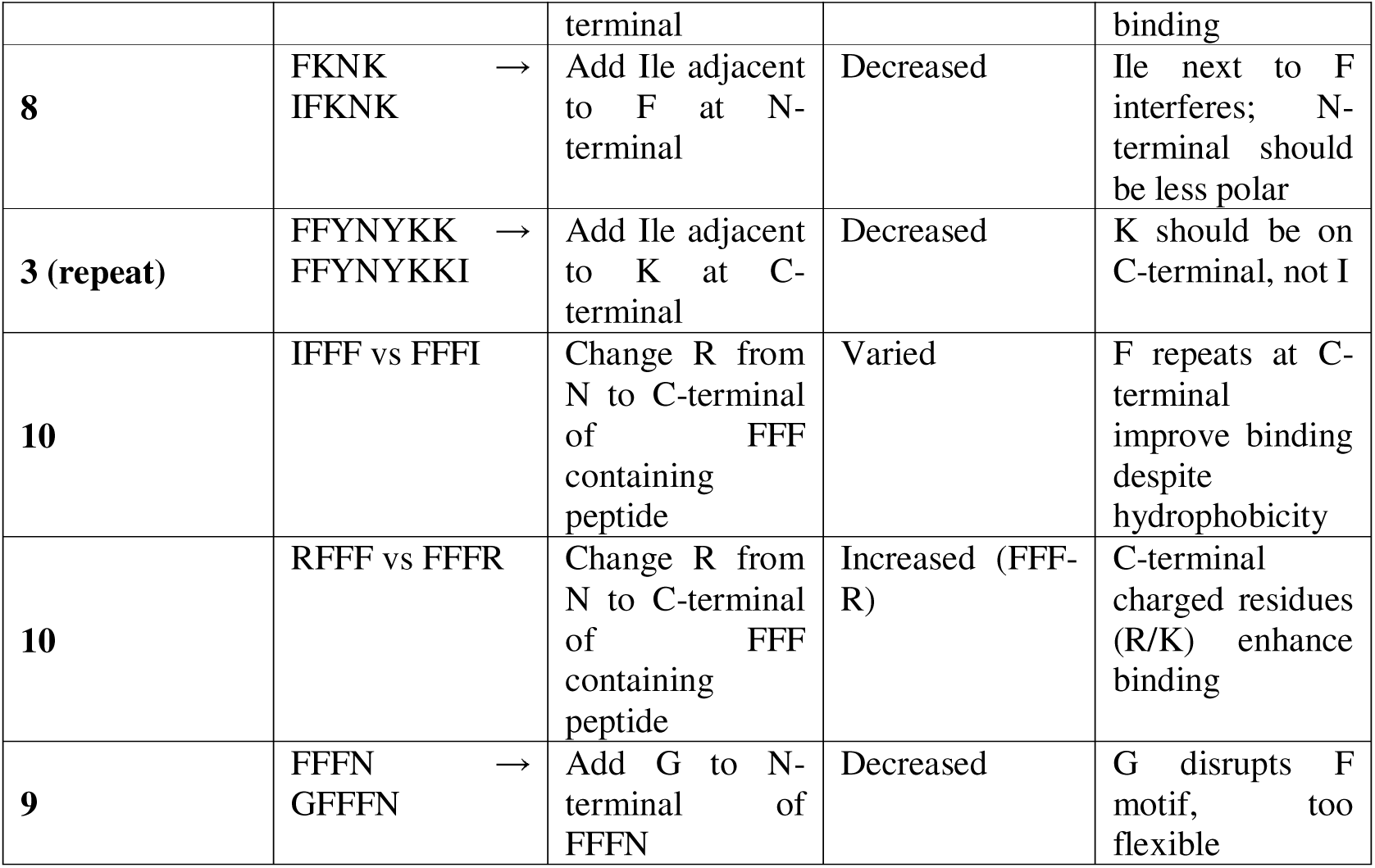
Peptide Binding Modification Summary. The optimal design features a hydrophobic N-terminal and a positively charged C-terminal.

In set 1, adding KK to FNKKN leads to formation of FNKKNKK. Comparison of FNKKN with FNKKNKK revealed that adding KK on C-terminal led to a significant increase in binding affinity (indicated by the more negative binding free energy of the sequence carrying KK on the C-terminal).

In set 2, when A was added to IIFR to form IIFRA, the peptide binding decreased noticeably, indicating that A interferes with binding of peptide with VIM-2 when added to C-terminal carrying R. Thus, A is interrupting the binding of R residue with VIM-2 (Figure 13).

**Figure 13.**
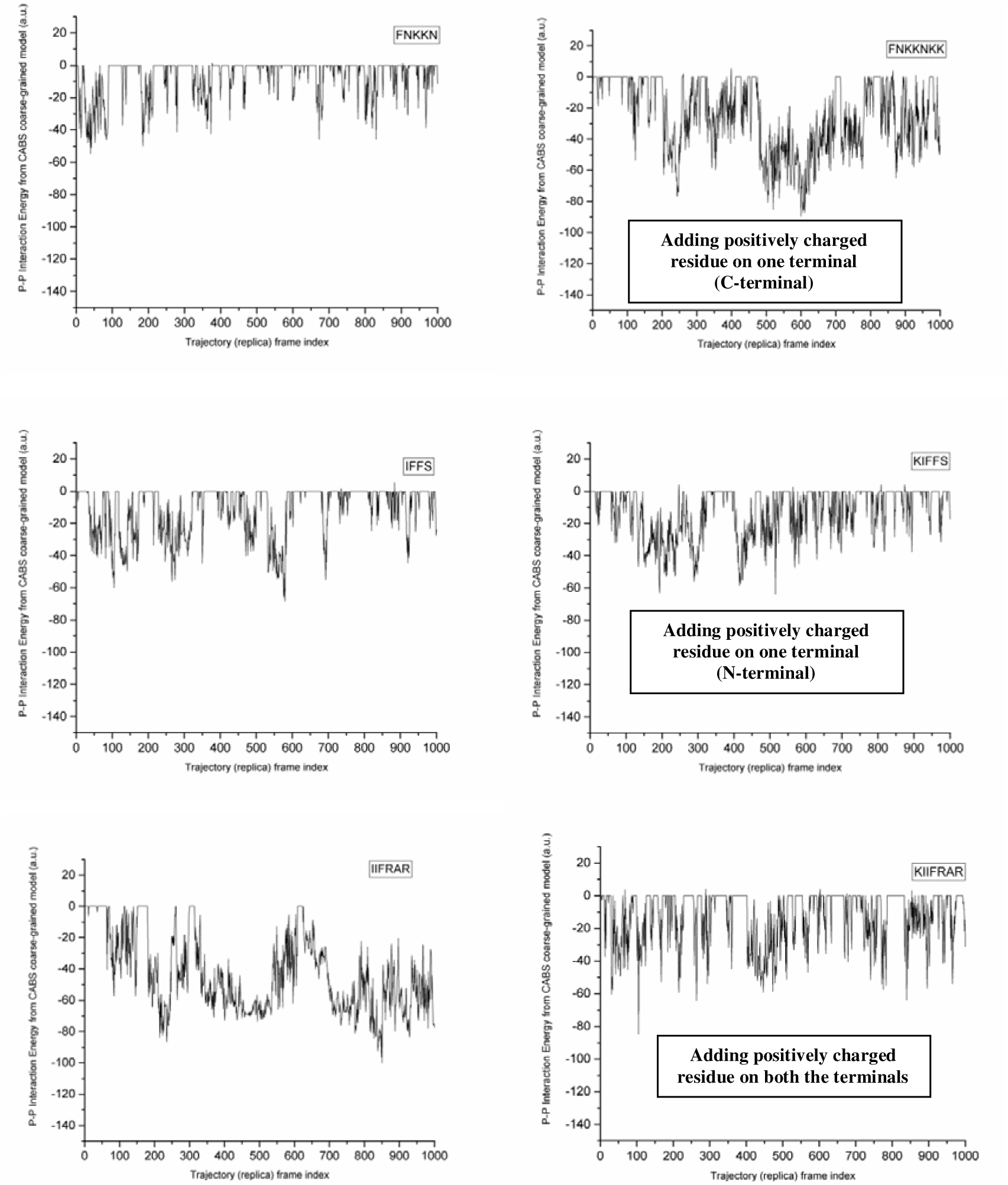
Suitable arrangement of amino acid types on the N and C terminals of the peptide sequences targetting MBLs like VIM-2. The figure clearly shows the need for one terminal to be hydrophobic and other to be positively charged with hydrophobic being on the N-terminal whereas the positively charged residue being on the C-terminal.

When an Ile residue was removed from the N-terminal of IIFRA to form IFRA, peptide binding further decreased showing the impact of Ile residues on peptide binding with VIM-2 (Figure 13).

When comparing IIFR with IIFRAR, it is seen that IIFRAR shows highly enhanced binding with VIM-2 because of the addition of the AR moiety, which has a neutral residue towards the N-terminal whereas a charged residue on the C-terminal.

When an R residue was added to IIFRA to form IIFRAR, the binding improved significantly, showing the necessity of R residue on C-terminal of peptide in order to bind with VIM-2. Again, when both Ile were removed from the N-terminal from IIFRAR (to form FRAR), the binding decreased noticeably ensuring the need for Ile residues for binding with VIM-2. To check having a positively charged residue like R or K on each of the terminals (both N and C) leads to good binding with VIM-2 or not, we added K to IIFRAR to form KIIFRAR, and found that the binding became insignificant as compared to the former. Thus, having positively charged residues on both terminals of the peptide could decrease binding with VIM-2. Thus, we anticipate that having one terminal as positively charged (mostly C-terminal, can be seen in discussion below) while other as neutral (N-terminal) favours peptide binding with the VIM-2 MBL (Figure 13).

In set 3, When Ile was added on C-terminal of FFYNYKK to form FFYNYKKI, it was found that binding is decreased largely, meaning that Ile on C-terminal adversely affects or interrupts the binding of K residues with the MBL. This also indicates that having two non-polar residues on both the terminals, does not help with peptide binding with MBL. In set 3 itself, when F was added to YNYKK on the N-terminal to form FYNYKK, the binding affinity increased, indicating that F is more advantageous than Y in peptide binding with the MBL, may be because F is more hydrophobic and therefore needed on N-terminal. Furthermore, adding another F to form FFYNYKK, increased the peptide binding with MBL further.

In set 3, we saw that adding F on the N-terminal when K is on C-terminal, increases binding affinity. However, in set 5, we see that when K is on N-terminal, then adding F on C-terminal of KGFF to form KGFFF further decreased binding, meaning that when less polar or more hydrophobic residue is on the N-terminal whereas the more polar or charged residue is on the C-terminal, the peptide binds exceptionally well with VIM-2 as compared to when the N and C terminals are vice-versa.

In set 4 also, when the positively-charged residue K was added on the N-terminal of IFFS to form KIFFS, the binding decreased. However, in set 6, we found that adding K on C-terminal of FFFNKKN to form FFNKKNK increased the binding with MBL significantly.

In set 8, when Ile was added to N-terminal adjacent to F in FKNK (to form IFKNK), it decreased binding, meaning that Ile adversely affects the binding of F residues with the MBL when lying adjacent to it.

We further studied the same in the presence of F repeats. In set 10, when Ile was added to FFF on either terminals to form IFFF and FFFI and the two were compared, it was observed that the F repeats are strengthening the binding of peptide with the MBL when present of the C-terminal even when F is more hydrophobic than Ile. This was in contrast to our findings for the previous peptides. To further test our observation, we performed the same study for Leu, Asn and Tyr, and observed similar results, even though F is more hydrophobic than Leu, Asn and Tyr. The study was then extended to Lys, Arg (which are extremely polar as compared to Phe), to check the impact of F repeats in presence of these residue on either terminal. When RFFF and FFFR were compared, it was found that FFFR showed better binding due to the positively charged residue on the C-terminal, similar to the finding described previously when F was present as a monopeptide or dipeptide in sequences on the N-terminal. Similarly, just like R when K was studied in combination with FFF, it was found that that K on the C-terminal showed excellent binding that surpass the sequence which has K on the N-terminal. This means that F repeats (F as tripeptide or more than 3 F) are able to overcome the effect of the adjacent residues, when the adjacent N-terminal residue is either non-polar or uncharged polar. Probably, this is why the naturally-derived gut AMPs have F repeats (FFFFF…) ahead of the non-polar residue glycine or isoleucine both of which are uncharged residues.

In set 9, we found that when G was added to FFFN to form GFFFN, the binding of peptide decreased significantly, indicating that G adversely impacts the binding of F with the VIM-2 MBL. However, a closer look at our study shows that G followed by F repeats (GFFFF) are found in the naturally derived AMPs above (parent AMPs-AMP10, AP13 and AMP18), especially in AMP18 where GFFFFFFF occurs at the C-terminal or end of the long peptide sequence which has the N-terminal as K. This indicates two things: (i) GFFFF might have an important role to play when binding with VIM-2, (ii) GFFF alone is not sufficient for binding with VIM-2, and that presence of electrovalent residues like K is extremely important. To describe, when G is added to FFFN (forming GFFFN), binding decreases significantly. This suggests that Gly may disrupt an optimal F-repeat binding motif (e.g., by introducing too much flexibility or altering secondary structure) as Gly is a highly flexible amino acid. The N-terminal position of G might interfere with Phe-based interactions critical for VIM-2 binding when no other stabilizing residues (like K) are present. In contrast, GFFFF appears in natural AMPs (e.g., AMP18: K…GFFFFFFF), where binding is retained. This suggests that Gly may act as a flexible linker that allows the F-repeat region to adopt a favorable conformation when other residues (like lysine, K) are present. This also suggests that the C-terminal position of G (after K or other charged residues) might not disrupt binding as severely as an N-terminal G.

The plots for all the analogs of set 2 have been depicted in Figure 12.

Plots for the analogs belonging to set 1 and set 3-10 can be seen in **Figure S1-S9 of the supplementary file.**

Overall, the study of our analogs shows that peptides with hydrophobic residues on N-terminal and positively charged residues on C-terminal are more likely to interact more strongly with the VIM-2 MBL. Probably, this is why, in our previous research we found that peptides with K on N-terminal while R on C-terminal bind more effectively with VIM-2 to form a highly stable complex as compared to when R is on N terminal and K on C-terminal, because R is more positively charged than K and therefore supported on the C-terminal with respect to K **(Anurag Anand et al., 2024)**.

**Figure 13** shows the suitable arrangement of amino acid types on the N and C terminals, in the peptide sequences meant to target MBLs like VIM-2.

We, hereby, propose a model mechanism by which Phe residues and their repeats can help in binding of peptide inhibitors with VIM-2 metallo-beta-lactamase and destabilizing the enzyme, which has been detailed in Figure 14.

**Figure 14.**
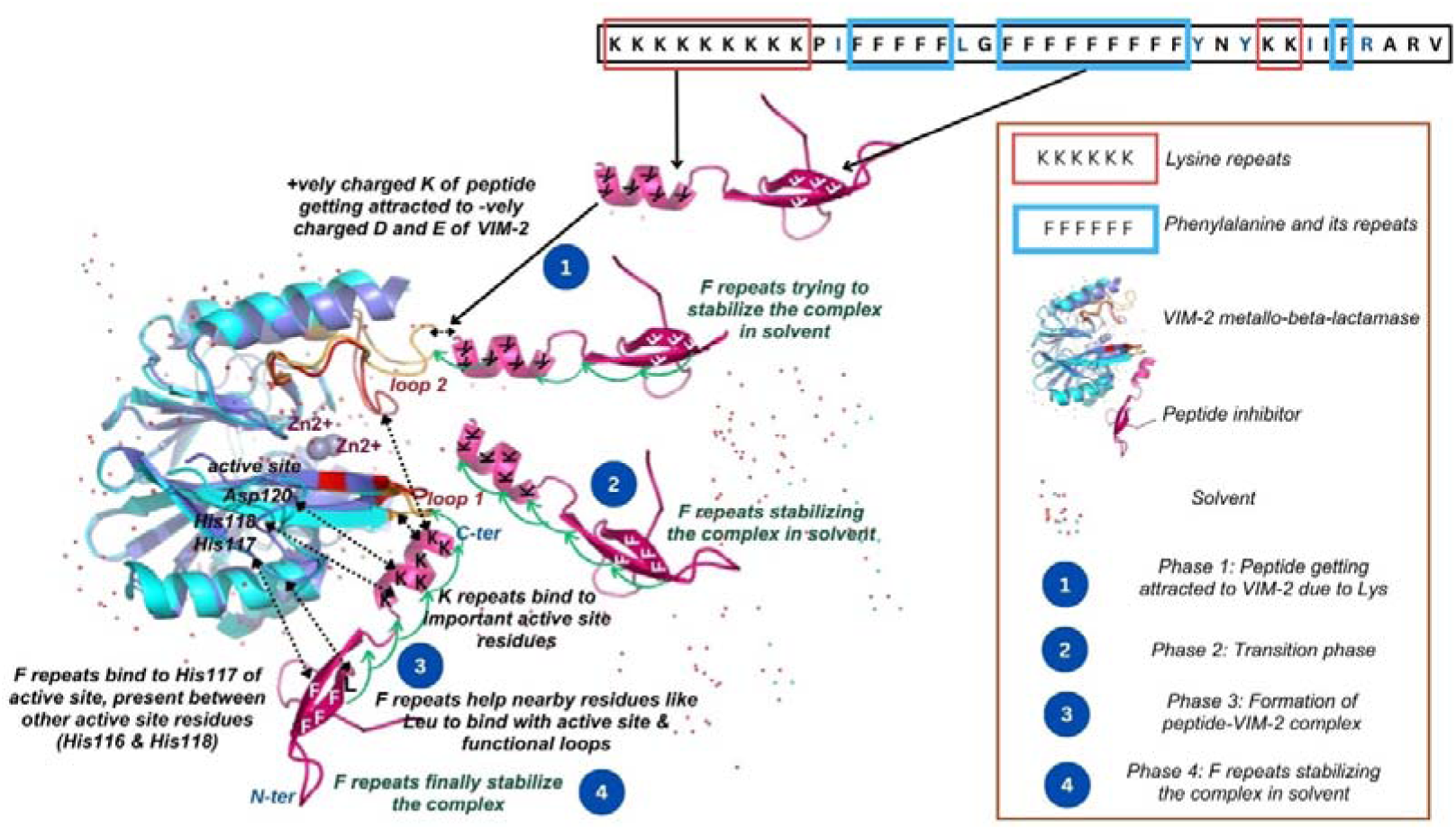
Model mechanism by which F-repeats help in binding of peptide inhibitors with VIM-2. F repeats have been shown to help peptide inhibitors in: i) binding with active site of VIM-2, ii) binding with functional loops of VIM-2, and iii) stabilizing the complex, especially in the presence of solvent. The image shows movement of F-repeats-containing-peptide (here AMP13) towards VIM-2, in 3 phases. In phase 1, peptide starts getting attracted to VIM-2’s Glu and Asp due to Lys present in peptide sequence. Additionally, F repeats try to provide stabilization to the peptide as well as the complex by keeping water molecules away. In phase 2, F repeats try to further stabilize the complex. Finally, in phase 3, we see that the final peptide-VIM-2 complex is formed where F repeats were finally able to stabilize it and several important interactions between the peptide inhibitor and VIM-2 remain intact. Finally, it is important to remember that the presence of hydrophobic residues like F is supported on N-terminal whereas that of positively charged residues like K is supported on C-terminal, when identifying MBL targeting peptides such as those targeting VIM-2 MBL.

### Conclusion and Future perspectives

F repeats exhibit very low polar solvation energy, indicative of the fact that F repeats play a role in stabilizing a VIM-2-peptide complex in presence of a solvent. Loop1 and loop2 analysis of VIM-2 revealed that F repeats bind with loop 2, thereby interfering in functioning of substrate binding at the active site. Additionally, F repeats have the ability to bind to several residues lying adjacent to or near to active site residues, which points towards the fact that F helps other residues like Asn and Lys to bind with the protein (even to the active site residues) of VIM-2. Further, F repeats help other peptide residues bind with loops of importance in VIM-2 which affect the functioning of active site. F repeats also bind to some active site residues and help nearby residues to interact with several active site residues. In our previous findings, we have seen that K repeats help in electrostatic interactions with VIM-2. Here as well, we see that K repeats in AMP10, 13 and 18 show similar interactions. Therefore, while K repeats are important for high binding affinity of peptide inhibitors towards VIM-2 and for direct binding with VIM-2 active site, F repeats are crucial for stabilizing the VIM-2-inhibitor complex, especially in presence of a solvent, followed by destabilization of loops, and inhibiting active site functioning by binding itself as well as facilitating binding of nearby residues to VIM-2. The 12-mer arginine-rich peptide (R12) exhibited the strongest binding to VIM-2 across all docking platforms (CABSdock, HawkDock, and MM/GBSA), followed by K12 and R6, highlighting the role of positively charged residues, especially arginine, in mediating stable, high-affinity interactions with VIM-2. In contrast, phenylalanine-rich peptides (F6 and F12) showed relatively weaker binding energies, although they appear to play a crucial role in stabilizing the complex by interacting with loop regions and facilitating nearby residue engagement. Further, our findings suggest that a balance of aromatic (F) and positively charged (K or R) residues could enhance peptide binding with VIM-2 through synergistic hydrophobic and electrostatic interactions. Most interestingly, the study of our analogs shows that peptides with hydrophobic residues on N-terminal and positively charged residues on C-terminal are more likely to interact more strongly with the VIM-2 MBL. Probably, this is why, in our previous research we found that peptides with K on N-terminal while R on C-terminal bind more effectively with VIM-2 to form a highly stable complex as compared to when R is on N terminal and K on C-terminal, because R is more positively charged than K and therefore supported on the C-terminal with respect to K **(Anurag Anand et al., 2024)**.

This work informs the rational design of next-generation peptide-based inhibitors targeting MBLs like VIM-2. Finally, this model can further be tested for other MBLs as well, to address the critical gaps in the invention of AMR therapeutics. Finally, we must also remember that nature supports amino acids that introduce stability so as to reduce off-target effects, such as, phenylalanine (as seen in this study), and lysine (as studied in our previous research).

## CRediT authorship contribution statement

Ananya Anurag Anand: Conceptualization, Methodology, Writing - original draft, review and editing. Sarfraz Anwar: Methodology. Rajat Kumar Mondal: Methodology. Sintu Kumar Samanta: Supervision and Conceptualization, Investigation, Resources, Writing – original draft, review and editing.

## Acknowledgements

Ananya Anurag Anand and Rajat Kumar Mondal would like to thank the Ministry of Education, Government of India, for their fellowship. Sarfraz Anwar would like to thank Council of Scientific and Industrial Research, University Grants Commission for his fellowship. All authors are extremely thankful to the Indian Institute of Information Technology Allahabad for providing us the research facility, especially the CCF.

## Data availability

Data will be made available on request.

## References

1. Aitha, M., Nix, J. C., Crowder, M. W., & Page, R. C. (2014). Structure of Zn(II)-bound metallo-beta-lactamse VIM-2 from Pseudomonas aeruginosa. Worldwide Protein Data Bank. 10.2210/pdb4nq2/pdb

2. Anand, A. A., Yadav, V., Anwar, S., Kshitiz, K., Sreedevi, P. C., & Samanta, S. K. (2025). The importance of phylogenetic studies in the evolution of beta-lactamases: Tracing antimicrobial resistance in priority pathogens. Gene & Protein in Disease, 0(0), 8278. 10.36922/gpd.8278

3. Anurag Anand, A., Amod, A., Anwar, S., Sahoo, A. K., Sethi, G., & Samanta, S. K. (2024). A comprehensive guide on screening and selection of a suitable AMP against biofilm-forming bacteria. Critical Reviews in Microbiology, 50(5), 859–878. 10.1080/1040841X.2023.2293019

4. Anurag Anand, A., Anwar, S., Sahoo, A. K., & Samanta, S. K. (2025). In silico identification of Corylifol C as a potential natural inhibitor of BfrB-Bfd interaction in Pseudomonas aeruginosa. Journal of Biomolecular Structure & Dynamics, 1–15. 10.1080/07391102.2025.2472171

5. Anurag Anand, A., Sahoo, A. K., & Samanta, S. K. (2024). Exploring the potential of designed peptides containing lysine and arginine repeats against VIM-2 metallo-beta-lactamases. International Journal of Peptide Research and Therapeutics, 30(4). 10.1007/s10989-024-10619-5

6. Bhat, B. A., Mir, R. A., Qadri, H., Dhiman, R., Almilaibary, A., Alkhanani, M., & Mir, M. A. (2023). Integrons in the development of antimicrobial resistance: critical review and perspectives. Frontiers in Microbiology, 14, 1231938. 10.3389/fmicb.2023.1231938

7. Christopeit, T., Yang, K. W., Yang, S. K., & Leiros, H. K. S. (2016). The structure of the metallo-β-lactamase VIM-2 in complex with a triazolylthioacetamide inhibitor. *Acta Crystallographica. Section F*, Structural Biology Communications, 72(Pt 11), 813–819. 10.1107/S2053230X16016113

8. Dalal, V., Dhankhar, P., Singh, V., Singh, V., Rakhaminov, G., Golemi-Kotra, D., & Kumar, P. (2021). Structure-based identification of potential drugs against FmtA of staphylococcus aureus: Virtual screening, molecular dynamics, MM-GBSA, and QM/MM. The Protein Journal, 40(2), 148–165. 10.1007/s10930-020-09953-6

9. Essmann, U., Perera, L., Berkowitz, M. L., Darden, T., Lee, H., & Pedersen, L. G. (1995). A smooth particle mesh Ewald method. The Journal of Chemical Physics, 103(19), 8577–8593. 10.1063/1.470117

10. Evans, D. J., & Holian, B. L. (1985). The Nose–Hoover thermostat. The Journal of Chemical Physics, 83(8), 4069–4074. 10.1063/1.449071

11. Garcia-Saez, I., Docquier, J.-D., Rossolini, G. M., & Dideberg, O. (2008). The three-dimensional structure of VIM-2, a Zn-β-lactamase from Pseudomonas aeruginosa in its reduced and oxidised form. Journal of Molecular Biology, 375(3), 604–611. doi:10.1016/j.jmb.2007.11.012

12. Garcia-Saez, I., Docquier, J.-D., Rossolini, G. M., & Dideberg, O. (2003). VIM-2, a Zn-beta-lactamase from Pseudomonas aeruginosa with Cys221 reduced. Worldwide Protein Data Bank. 10.2210/pdb1ko3/pdb

13. Kumari, R., & Dalal, V. (2022). Identification of potential inhibitors for LLM of Staphylococcus aureus: structure-based pharmacophore modeling, molecular dynamics, and binding free energy studies. Journal of Biomolecular Structure & Dynamics, 40(20), 9833–9847. 10.1080/07391102.2021.1936179

14. Livermore, D. M. (2009). Has the era of untreatable infections arrived? The Journal of Antimicrobial Chemotherapy, 64 *Suppl 1* (Supplement 1), i29-36. 10.1093/jac/dkp255

15. Lucic, A., Hinchliffe, P., & Schofield, C. (2023). Metallo beta-lactamase VIM2 with compound AK110. Worldwide Protein Data Bank. 10.2210/pdb8pjm/pdb

16. Lucic, A., & Schofield, C. J. (2021). Structure of VIM-2 metallo-beta-lactamase with hydrolysed Faropenem imine product. Worldwide Protein Data Bank. 10.2210/pdb7a5z/pdb

17. Mojica, M. F., Rossi, M.-A., Vila, A. J., & Bonomo, R. A. (2022). The urgent need for metallo-β-lactamase inhibitors: an unattended global threat. The Lancet Infectious Diseases, 22(1), e28–e34. 10.1016/S1473-3099(20)30868-9

18. Mondal, R. K., Sen, D., Arya, A., & Samanta, S. K. (2023). Developing anti-microbial peptide database version 1 to provide comprehensive and exhaustive resource of manually curated AMPs. Scientific Reports, 13(1), 17843. 10.1038/s41598-023-45016-3

19. Page, M. I., & Badarau, A. (2008). The mechanisms of catalysis by metallo beta-lactamases. Bioinorganic Chemistry and Applications, 2008(1), 576297. 10.1155/2008/576297

20. Parrinello, M., & Rahman, A. (1981). Polymorphic transitions in single crystals: A new molecular dynamics method. Journal of Applied Physics, 52(12), 7182–7190. 10.1063/1.328693

21. Pingali, M. S., Singh, A., Anurag Anand, A., Gupta, S. K., Sahoo, A. K., Varadwaj, P. K., & Samanta, S. K. (2024). Identification of naturally occurring compounds as alternatives to radiation therapy for treatment of small cell lung cancer. Journal of Biomolecular Structure & Dynamics, 42(21), 11942–11953. 10.1080/07391102.2023.2265505

22. Poirel, L., Naas, T., Nicolas, D., Collet, L., Bellais, S., Cavallo, J. D., & Nordmann, P. (2000). Characterization of VIM-2, a carbapenem-hydrolyzing metallo-beta-lactamase and its plasmid- and integron-borne gene from a Pseudomonas aeruginosa clinical isolate in France. Antimicrobial Agents and Chemotherapy, 44(4), 891–897. 10.1128/AAC.44.4.891-897.2000

23. Prakash, A., Kumar, V., Meena, N. K., & Lynn, A. M. (2018). Elucidation of the structural stability and dynamics of heterogeneous intermediate ensembles in unfolding pathway of the N-terminal domain of TDP-43. RSC Advances, 8(35), 19835–19845. 10.1039/c8ra03368d

24. Queenan, A. M., & Bush, K. (2007). Carbapenemases: the versatile beta-lactamases. Clinical Microbiology Reviews, 20(3), 440–458, table of contents. 10.1128/CMR.00001-07

25. Rossino, G., Marchese, E., Galli, G., Verde, F., Finizio, M., Serra, M., Linciano, P., & Collina, S. (2023). Peptides as therapeutic agents: Challenges and opportunities in the Green transition era. *Molecules (Basel*, Switzerland), 28(20). 10.3390/molecules28207165

26. Rotondo, C. M., Marrone, L., Goodfellow, V. J., Ghavami, A., Labbé, G., Spencer, J., Dmitrienko, G. I., & Siemann, S. (2015). Arginine-containing peptides as potent inhibitors of VIM-2 metallo-β-lactamase. Biochimica et Biophysica Acta, 1850(11), 2228–2238. 10.1016/j.bbagen.2015.07.012

27. Singh, A., Amod, A., Mulpuru, V., Mishra, N., Sahoo, A. K., & Samanta, S. K. (2023). Finding novel AMPs secreted from the human microbiome as potent antibacterial and antibiofilm agents and studying their synergistic activity with Ag NCs. ACS Applied Bio Materials, 6(9), 3674–3682. 10.1021/acsabm.3c00302

28. Singh, A., Amod, A., Pandey, P., Bose, P., Pingali, M. S., Shivalkar, S., Varadwaj, P. K., Sahoo, A. K., & Samanta, S. K. (2022). Bacterial biofilm infections, their resistance to antibiotics therapy and current treatment strategies. *Biomedical Materials (Bristol*, England), 17(2), 022003. 10.1088/1748-605X/ac50f6

29. Toleman, M. A., Vinodh, H., Sekar, U., Kamat, V., & Walsh, T. R. (2007). blaVIM-2-harboring integrons isolated in India, Russia, and the United States arise from an ancestral class 1 integron predating the formation of the 3’ conserved sequence. Antimicrobial Agents and Chemotherapy, 51(7), 2636–2638. 10.1128/AAC.01043-06

30. Wachino, J. (2024). Crystal structure of VIM-2 metallo-beta-lactamase in complex with 10-HHIA: 8I52 [Data set]. In Worldwide Protein Data Bank. Worldwide Protein Data Bank.

31. Weng, G., Wang, E., Wang, Z., Liu, H., Zhu, F., Li, D., & Hou, T. (2019). HawkDock: a web server to predict and analyze the protein-protein complex based on computational docking and MM/GBSA. Nucleic Acids Research, 47(W1), W322– W330. 10.1093/nar/gkz397

32. Xiao, J., Fang, M., Shi, Y., Chen, H., Shen, B., Chen, J., Lao, X., Xu, H., & Zheng, H. (2015). Identification and validation novel of VIM-2 metallo-β-lactamase tripeptide inhibitors. Molecular Informatics, 34(8), 559–567. 10.1002/minf.201400178

33. Yan, Y.-H., Ding, H.-S., Zhu, K.-R., Mu, B.-S., Zheng, Y., Huang, M.-Y., Zhou, C., Li, W.-F., Wang, Z., Wu, Y., & Li, G.-B. (2023). Metal binding pharmacophore click-derived discovery of new broad-spectrum metallo-β-lactamase inhibitors. European Journal of Medicinal Chemistry, 257(115473), 115473. 10.1016/j.ejmech.2023.115473

34. Yan, Y.-H., Li, W., Chen, W., Li, C., Zhu, K.-R., Deng, J., Dai, Q.-Q., Yang, L.-L., Wang, Z., & Li, G.-B. (2022). Structure-guided optimization of 1H-imidazole-2-carboxylic acid derivatives affording potent VIM-Type metallo-β-lactamase inhibitors. European Journal of Medicinal Chemistry, 228(113965), 113965. 10.1016/j.ejmech.2021.113965

