## Supplementary Data for "Influence of F Repeats and Terminal Residue Polarity on Peptide–VIM-2 Metallo-β-Lactamase Interactions"

Dr. Sintu Kumar Samanta

Assistant Professor

Department of Applied Sciences,

Indian Institute of Information Technology Allahabad,

Allahabad-211012, India,

**Table S1. Comparison of different PDB structures of VIM-2 in *P. aeruginosa***

·

| **S. No.** | **Entry PDB**  **ID** | **Structure Title** | **Refinement**  **Resolution (Å)** | **Parameters** | **Sequence**  **Length** |
| --- | --- | --- | --- | --- | --- |
| 1 | 1KO3 **(Garcia-Saez et al., 2003)** | VIM-2, a Zn-beta-lactamase from *Pseudomonas aeruginosa* with Cys221 reduced | 1.91 Å | R free \| 0.203; Clashscore \| 6; Ramachandranoutliers \| 0; Sidechain outliers \| 3.3%; RSRZ outliers \| 7.8% | 230 |
| 2 | 1KO2 **(Garcia-Saez et al., 2003)** | VIM-2, a Zn-beta-lactamase from *Pseudomonas aeruginosa* with an oxidized Cys (cysteinesulfonic) | 2.20 Å | R free \| 0.251; Clashscore \| 7; Ramachandran outliers \| 0.4%; Sidechain outliers \| 2.2%; RSRZ outliers \| 2.2% | 230 |
| 3 | 4NQ2 (**Aitha et al., 2014)** | Structure of Zn(II)-bound metallo-beta-lactamse VIM-2 from *Pseudomonas aeruginosa* | 1.55 Å | R free \| 0.208; Clashscore \| 3; Ramachandran outliers \| 0; Sidechain outliers \| 0.6%; RSRZ outliers \| 2.2% | 261 |
| 4 | 8PJM (**Lucic et al., 2023)** | Metallo beta-lactamase VIM2 with compound AK110 | 1.44 Å | R free \| 0.155; Clashscore \| 4; Ramachandran outliers \| 0.4%; Sidechain outliers \| 1.1%; RSRZ outliers \| 1.7% | 242 |
| 5 | 7DUY **(Yan et al., 2022)** | Crystal structure of VIM-2 MBL in complex with 1-(2-(1H-1,2,3-triazol-1-yl)ethyl)-1H-imidazole-2-carboxylic acid | 2.00 Å | R free \| 0.216; Clashscore \| 2; Ramachandran outliers \| 0.4%; Sidechain outliers \| 1.1%; RSRZ outliers \| 2.6% | 231 |
| 6 | 8I52 (**Wachino, 2024)** | Crystal structure of VIM-2 metallo-beta-lactamase in complex with 10-HHIA | 1.58 Å | R free \| 0.192; Clashscore \| 3; Ramachandran outliers \| 0.4%; Sidechain outliers \| 1.1%; RSRZ outliers \| 0.6% | 246 |
| 7 | 7A5Z (**Lucic et al., 2021)** | Structure of VIM-2 metallo-beta-lactamase with hydrolysed Faropenem imine product | 1.29 Å | R free \| 0.142; Clashscore \| 4; Ramachandran outliers \| 0.4%; Sidechain outliers \| 1.1%; RSRZ outliers \| 0.4% | 242 |
| 8 | 7YHD (Yan et al., 2023)) | Crystal structure of VIM-2 MBL in complex with 3-(4-(4-(2-aminoethoxy)phenyl)-1H-1,2,3-triazol-1-yl)phthalic acid | 1.70 Å | R free \| 0.342; Clashscore \| 13; Ramachandran outliers \| 3.1%; Sidechain outliers \| 3.3%; RSRZ outliers \| 20.3% | 231 |
| 9 | 8HYD (Yan et al., 2023)) | Crystal structure of B1 VIM-2 MBL in complex with 2-amino-5-(2-(thiophen-2-yl)ethyl)thiazole-4-carboxylic acid | 1.71 Å | R free \| 0.210; Clashscore \| 4; Ramachandran outliers \| 0.4%; Sidechain outliers \| 0.5%; RSRZ outliers \| 7.4% | 231 |
| 10 | 7YHB (Yan et al., 2023)) | Crystal structure of VIM-2 MBL in complex with (2-(4-phenyl-1H-1,2,3-triazol-1-yl)benzyl)phosphonic acid | 1.43 Å | R free \| 0.311; Clashscore \| 5; Ramachandran outliers \| 0.7%; Sidechain outliers \| 0.8%; RSRZ outliers \| 15.8% | 231 |
| 11 | 7DZ1(Yan et al., 2023)) | Crystal structure of VIM-2 MBL in complex with 1-benzyl-5-methyl-1H-imidazole-2-carboxylic acid | 2.71 Å | R free \| 0.268; Clashscore \| 7; Ramachandran outliers \| 0.9%; Sidechain outliers \| 6.0%; RSRZ outliers \| 0.4% | 231 |
| 12 | 7DZ0 (Yan et al., 2023)) | Crystal structure of VIM-2 MBL in complex with 1-(but-3-en-1-yl)-5-methyl-1H-imidazole-2-carboxylic acid | 3.23 Å | R free \| 0.293; Clashscore \| 13; Ramachandran outliers \| 2.4%; Sidechain outliers \| 1.9%; RSRZ outliers \| 1.1% | 231 |

**Table S2. Fraction of modeled and unmodeled regions of different VIM-2 structures**

| **PDB ID** | **Structure Title** | **%Modeled Segment** | **%Unmodeled Segment** | **Ligands attached** |
| --- | --- | --- | --- | --- |
| 1KO3 | VIM-2, a Zn-beta-lactamase from *Pseudomonas aeruginosa* with Cys221 reduced | 86% | 0% | 3 Zn, 2 ACT, 1 Cl, 1 OH |
| 1KO2 | VIM-2, a Zn-beta-lactamase from *Pseudomonas aeruginosa* with an oxidized Cys (cysteinesulfonic) | 84% | 0% | 2 Zn, 2 ACT |
| 4NQ2 | Structure of Zn(II)-bound metallo-beta-lactamse VIM-2 from *Pseudomonas aeruginosa* | 83% | 12% | 3 Zn, 2 ACT |
| 8PJM | Metallo beta-lactamase VIM2 with compound AK110 | 94% | 4% | 1 A1H7J,  3 Zn, 3 Cl, 1 Na |
| 7DUY | Crystal structure of VIM-2 MBL in complex with 1-(2-(1H-1,2,3-triazol-1-yl)ethyl)-1H-imidazole-2-carboxylic acid | 93% | 0% | 1 HLL, 2 Zn |
| 8I52 | Crystal structure of VIM-2 metallo-beta-lactamase in complex with 10-HHIA | Chain A: 88%  Chain D: 88% | Chain A: 6%  Chain D: 6% | 5 Zn, 4 FMT, 1 OR6 |
| 7A5Z | Structure of VIM-2 metallo-beta-lactamase with hydrolysed Faropenem imine product | 88% | 4% | 1 QZH, 3 Zn, 2 Cl, 1 MG, 1 Na |
| 7YHD | Crystal structure of VIM-2 MBL in complex with 3-(4-(4-(2-aminoethoxy)phenyl)-1H-1,2,3-triazol-1-yl)phthalic acid | 71% | 0% | 1 IU7, 2 Zn |
| 8HYD | Crystal structure of B1 VIM-2 MBL in complex with 2-amino-5-(2-(thiophen-2-yl)ethyl)thiazole-4-carboxylic acid | 93% | 7% | 1 60 J, 2 Zn |
| 7YHB | Crystal structure of VIM-2 MBL in complex with (2-(4-phenyl-1H-1,2,3-triazol-1-yl)benzyl)phosphonic acid | Chain A: 89%  Chain B: 88% | Chain A:11%  Chain B:11% | 2 ITV, 4 Zn |
| 7DZ1 | Crystal structure of VIM-2 MBL in complex with 1-benzyl-5-methyl-1H-imidazole-2-carboxylic acid | Chain A: 82%  Chain B: 77% | Chain A: 0%  Chain B: 0% | 2 HQ3, 4 Zn |
| 7DZ0 | Crystal structure of VIM-2 MBL in complex with 1-(but-3-en-1-yl)-5-methyl-1H-imidazole-2-carboxylic acid | Chain A: 70%  Chain B:68% | Chain A: 0%  Chain B: 0% | 2 HQ0, 4 Zn |

**Table S3.** Details of active site residues in VIM-2

| **Active Site Residue In VIM-2 Structure Considered For Docking** | **Corresponding Residue In Reference Structure (PDB ID: 1KO3)** |
| --- | --- |
| Y 36 | Y 67 |
| Y 170 | Y 224 |
| S 38 | S 69 |
| H 83 | H 116 |
| H 85 | H 118 |
| H 148 | H 196 |
| H 209 | H 263 |
| D 87 | D 120 |
| R 88 | R 121 |
| R 174 | R 228 |
| Cys 167 | Cys 221 |

**Table S4.** Details of residues in VIM-2 structure involved in docking vs residues in the original structure (PDB ID: 1KO3)

| Amino acid residues in VIM-2 structure involved in docking | Amino acid residues in original VIM-2 structure PDB ID 1KO3 |
| --- | --- |
| GLU 1  TYR 2  PRO 3  THR 4  VAL 5  SER 6  GLU 7  ILE 8  PRO 9  VAL 10  GLY 11  GLU 12  VAL 13  ARG 14  LEU 15  TYR 16  GLN 17  ILE 18  ALA 19  ASP 20  GLY 21  VAL 22  TRP 23  SER 24  HIS 25  ILE 26  ALA 27  THR 28  GLN 29  SER 30  PHE 31  ASP 32  GLY 33  ALA 34  VAL 35  TYR 36  PRO 37  SER 38  ASN 39  GLY 40  LEU 41  ILE 42  VAL 43  ARG 44  ASP 45  GLY 46  ASP 47  GLU 48  LEU 49  LEU 50  LEU 51  ILE 52  ASP 53  THR 54  ALA 55  TRP 56  GLY 57  ALA 58  LYS 59  ASN 60  THR 61  ALA 62  ALA 63  LEU 64  LEU 65  ALA 66  GLU 67  ILE 68  GLU 69  LYS 70  GLN 71  ILE 72  GLY 73  LEU 74  PRO 75  VAL 76  THR 77  ARG 78  ALA 79  VAL 80  SER 81  THR 82  HIS 83  PHE 84  HIS 85  ASP 86  ASP 87  ARG 88  VAL 89  GLY 90  GLY 91  VAL 92  ASP 93  VAL 94  LEU 95  ARG 96  ALA 97  ALA 98  GLY 99  VAL 100  ALA 101  THR 102  TYR 103  ALA 104  SER 105  PRO 106  SER 107  THR 108  ARG 109  ARG 110  LEU 111  ALA 112  GLU 113  VAL 114  GLU 115  GLY 116  ASN 117  GLU 118  ILE 119  PRO 120  THR 121  HIS 122  SER 123  LEU 124  GLU 125  GLY 126  LEU 127  SER 128  SER 129  SER 130  GLY 131  ASP 132  ALA 133  VAL 134  ARG 135  PHE 136  GLY 137  PRO 138  VAL 139  GLU 140  LEU 141  PHE 142  TYR 143  PRO 144  GLY 145  ALA 146  ALA 147  HIS 148  SER 149  THR 150  ASP 151  ASN 152  LEU 153  VAL 154  VAL 155  TYR 156  VAL 157  PRO 158  SER 159  ALA 160  SER 161  VAL 162  LEU 163  TYR 164  GLY 165  GLY 166  CYS 167  ALA 168  ILE 169  TYR 170  GLU 171  LEU 172  SER 173  ARG 174  THR 175  SER 176  ALA 177  GLY 178  ASN 179  VAL 180  ALA 181  ASP 182  ALA 183  ASP 184  LEU 185  ALA 186  GLU 187  TRP 188  PRO 189  THR 190  SER 191  ILE 192  GLU 193  ARG 194  ILE 195  GLN 196  GLN 197  HIS 198  TYR 199  PRO 200  GLU 201  ALA 202  GLN 203  PHE 204  VAL 205  ILE 206  PRO 207  GLY 208  HIS 209  GLY 210  LEU 211  PRO 212  GLY 213  GLY 214  LEU 215  ASP 216  LEU 217  LEU 218  LYS 219  HIS 220  THR 221  THR 222  ASN 223  VAL 224  VAL 225  LYS 226  ALA 227  HIS 228  THR 229  ASN 230 | GLU 30  TYR 31  PRO 32  THR 33  VAL 34  SER 35  GLU 36  ILE 37  PRO 38  VAL 39  GLY 40  GLU 41  VAL 42  ARG 43  LEU 44  TYR 45  GLN 47  ILE 48  ALA 49  ASP 50  GLY 51  VAL 52  TRP 53  SER 54  HIS 55  ILE 56  ALA 57  THR 58  GLN 59  SER 60  PHE 61  ASP 62  GLY 63  ALA 64  VAL 66  TYR 67  PRO 68  SER 69  ASN 70  GLY 71  LEU 72  ILE 73  VAL 74  ARG 75  ASP 76  GLY 77  ASP 78  GLU 79  LEU 80  LEU 81  LEU 82  ILE 83  ASP 84  THR 85  ALA 86  TRP 87  GLY 88  ALA 89  LYS 90  ASN 91  THR 92  ALA 93  ALA 94  LEU 95  LEU 96  ALA 97  GLU 98  ILE 99  GLU 100  LYS 102  GLN 103  ILE 104  GLY 105  LEU 106  PRO 107  VAL 109  THR 110  ARG 111  ALA 112  VAL 113  SER 114  THR 115  HIS 116  PHE 117  HIS 118  ASP 119  ASP 120  ARG 121  VAL 122  GLY 123  GLY 124  VAL 125  ASP 126  VAL 127  LEU 128  ARG 129  ALA 130  ALA 131  GLY 133  VAL 134  ALA 135  THR 136  TYR 137  ALA 138  SER 139  PRO 140  SER 141  THR 142  ARG 143  ARG 144  LEU 145  ALA 146  GLU 147  VAL 148  GLU 149  GLY 150  ASN 165  GLU 166  ILE 167  PRO 168  THR 169  HIS 170  SER 171  LEU 172  GLU 173  GLY 174  LEU 175  SER 176  SER 177  SER 178  GLY 179  ASP 180  ALA 181  VAL 182  ARG 183  PHE 184  GLY 185  PRO 186  VAL 187  GLU 188  LEU 189  PHE 190  TYR 191  PRO 192  GLY 193  ALA 194  ALA 195  HIS 196  SER 197  THR 198  ASP 199  ASN 200  LEU 201  VAL 202  VAL 203  TYR 204  VAL 205  PRO 209  SER 210  ALA 211  SER 215  VAL 216  LEU 217  TYR 218  GLY 219  GLY 220  CYS 221  ALA 222  ILE 223  TYR 224  GLU 225  LEU 226  SER 227  ARG 228  THR 229  SER 230  ALA 231  GLY 232  ASN 233  VAL 234  ALA 235  ASP 236  ALA 237  ASP 238  LEU 239  ALA 240  GLU 241  TRP 242  PRO 243  THR 244  SER 245  ILE 246  GLU 247  ARG 248  ILE 249  GLN 250  GLN 251  HIS 252  TYR 253  PRO 254  GLU 255  ALA 256  GLN 257  PHE 258  VAL 259  ILE 260  PRO 261  GLY 262  HIS 263  GLY 264  LEU 265  PRO 277  GLY 278  GLY 279  LEU 280  ASP 281  LEU 282  LEU 283  LYS 284  HIS 285  THR 286  THR 287  ASN 288  VAL 289  VAL 290  LYS 291  ALA 292  HIS 293  THR 294  ASN 295 |

**Table S5. Residue decomposition analysis for AMP13 when bound to VIM-2**

| RESIDUE_ID | VDW | ELE | GB | SA | TOTAL |
| --- | --- | --- | --- | --- | --- |
| L-B-LYS-1 | -0.14 | -213.7 | 212.78 | 0 | -1.06 |
| L-B-LYS-2 | -0.86 | -135.32 | 132.98 | -0.48 | -3.68 |
| L-B-LYS-3 | -2.7 | -163.77 | 166.31 | -1.02 | -1.19 |
| L-B-LYS-4 | -1.28 | -138.77 | 137.73 | -0.28 | -2.6 |
| L-B-LYS-5 | -0.15 | -94.32 | 93.55 | 0 | -0.92 |
| L-B-LYS-6 | -2.51 | -135.06 | 144.01 | -0.88 | 5.55 |
| L-B-LYS-7 | -2.68 | -173.48 | 172.29 | -0.83 | -4.7 |
| L-B-LYS-8 | -0.17 | -95.57 | 94.7 | 0 | -1.04 |
| L-B-LYS-9 | -0.24 | -95.4 | 95.35 | 0 | -0.29 |
| L-B-PRO-10 | -0.1 | 2.28 | -2.14 | 0 | 0.04 |
| L-B-ILE-11 | -1.22 | -1.39 | 1.6 | -0.16 | -1.17 |
| L-B-PHE-12 | -0.36 | 2.54 | -1.87 | -0.02 | 0.29 |
| L-B-PHE-13 | -5.32 | -2.74 | 3.47 | -0.6 | -5.19 |
| L-B-PHE-14 | -0.76 | 1.86 | -1.01 | -0.14 | -0.04 |
| L-B-PHE-15 | -3.23 | -1.38 | 1.73 | -0.57 | -3.46 |
| L-B-PHE-16 | -0.28 | 1.41 | -1.11 | -0.05 | -0.03 |
| L-B-LEU-17 | -0.1 | -0.02 | 0.09 | 0 | -0.03 |
| L-B-GLY-18 | -0.01 | -0.41 | 0.43 | 0 | 0.02 |
| L-B-PHE-19 | -0.01 | -0.9 | 0.91 | 0 | 0 |
| L-B-PHE-20 | -0.02 | 0.6 | -0.54 | 0 | 0.04 |
| L-B-PHE-21 | -0.03 | -0.34 | 0.39 | 0 | 0.02 |
| L-B-PHE-22 | -0.08 | 0.45 | -0.34 | 0 | 0.03 |
| L-B-PHE-23 | -0.11 | -0.46 | 0.73 | 0 | 0.16 |
| L-B-PHE-24 | -1.08 | 1.33 | -0.51 | -0.13 | -0.38 |
| L-B-PHE-25 | -0.23 | -0.95 | 1.13 | 0 | -0.05 |
| L-B-PHE-26 | -2.79 | 0.11 | 0.02 | -0.38 | -3.04 |
| L-B-PHE-27 | -0.1 | 0.08 | 0.07 | 0 | 0.05 |
| L-B-TYR-28 | -1.79 | -4.11 | 4.01 | -0.22 | -2.12 |
| L-B-ASN-29 | -0.2 | 1.29 | -0.87 | 0 | 0.22 |
| L-B-TYR-30 | -3.86 | -0.03 | 0.76 | -0.53 | -3.67 |
| L-B-LYS-31 | -0.46 | -87.2 | 86.63 | -0.05 | -1.08 |
| L-B-LYS-32 | -0.25 | -77.74 | 77.19 | 0 | -0.8 |
| L-B-ILE-33 | -2.83 | -2.1 | 2.44 | -0.53 | -3.02 |
| L-B-ILE-34 | -0.17 | 1.19 | -0.91 | 0 | 0.1 |
| L-B-PHE-35 | -0.51 | 0.01 | 0.12 | -0.01 | -0.38 |
| L-B-ARG-36 | -0.08 | -67.8 | 67.13 | 0 | -0.75 |
| L-B-ALA-37 | -0.04 | -0.05 | 0.12 | 0 | 0.02 |
| L-B-ARG-38 | -0.04 | -63.47 | 62.83 | 0 | -0.68 |
| L-B-VAL-39 | -0.02 | 59.97 | -59.15 | 0 | 0.79 |

**Table S6. Residue decomposition analysis for AMP10 when bound to VIM-2**

| RESIDUE_ID | VDW | ELE | GB | SA | TOTAL |
| --- | --- | --- | --- | --- | --- |
| L-B-GLY-1 | -0.23 | -102.7 | 103.25 | -0.08 | 0.25 |
| L-B-GLY-2 | -0.27 | 3.28 | -2.6 | -0.04 | 0.36 |
| L-B-PHE-3 | -2.33 | -4.26 | 5.07 | -0.32 | -1.84 |
| L-B-PHE-4 | -1.77 | 1.88 | -0.3 | -0.31 | -0.5 |
| L-B-PHE-5 | -0.93 | -1.88 | 2.01 | -0.01 | -0.81 |
| L-B-ASN-6 | -1.38 | -6.55 | 6.5 | -0.2 | -1.64 |
| L-B-LYS-7 | -0.06 | -80.86 | 79.96 | 0 | -0.95 |
| L-B-LYS-8 | -0.15 | -93.02 | 92.41 | 0 | -0.76 |
| L-B-ASN-9 | -1.27 | -7.8 | 4.61 | -0.3 | -4.75 |
| L-B-LYS-10 | -0.11 | -86.38 | 85.47 | 0 | -1.02 |
| L-B-LYS-11 | -1.06 | -173 | 168.82 | -0.48 | -5.72 |
| L-B-LEU-12 | -2.73 | 0.44 | 0.2 | -0.21 | -2.29 |
| L-B-TYR-13 | -0.13 | 1.44 | -1.3 | 0 | 0.01 |
| L-B-LYS-14 | -0.11 | -101.53 | 100.64 | 0 | -1 |
| L-B-LYS-15 | -1.38 | -145.1 | 148.22 | -0.6 | 1.14 |
| L-B-LYS-16 | -1.17 | -167.08 | 166.41 | -0.28 | -2.12 |
| L-B-LYS-17 | -0.06 | -85.59 | 84.69 | 0 | -0.95 |
| L-B-LYS-18 | -0.17 | -95.82 | 95.09 | 0 | -0.9 |
| L-B-ILE-19 | -0.99 | -4.47 | 4.64 | -0.17 | -0.99 |
| L-B-PHE-20 | -0.4 | 3.57 | -2.69 | 0 | 0.47 |
| L-B-PHE-21 | -1.3 | -0.7 | 1.58 | -0.28 | -0.7 |
| L-B-SER-22 | -1.32 | -1.53 | 2.45 | -0.4 | -0.8 |
| L-B-GLY-23 | -0.38 | 0.28 | 0.15 | -0.02 | 0.02 |
| L-B-ARG-24 | -3.44 | -129.71 | 131.36 | -0.78 | -2.57 |
| L-B-VAL-25 | -0.08 | 2.03 | -1.83 | 0 | 0.12 |
| L-B-VAL-26 | -0.09 | -2.34 | 2.44 | 0 | 0.01 |
| L-B-ALA-27 | -0.03 | -2.91 | 2.91 | 0 | -0.03 |
| L-B-ASP-28 | -0.03 | 88.06 | -86.79 | 0 | 1.23 |
| L-B-LEU-29 | -0.03 | -0.72 | 0.73 | 0 | -0.01 |
| L-B-PRO-30 | -0.01 | 1.93 | -1.9 | 0 | 0.02 |
| L-B-GLY-31 | 0 | 1.05 | -1.02 | 0 | 0.02 |
| L-B-GLU-32 | -0.02 | 84.9 | -83.73 | 0 | 1.15 |
| L-B-SER-33 | 0 | -0.52 | 0.52 | 0 | 0 |
| L-B-SER-34 | 0 | -0.83 | 0.82 | 0 | -0.01 |
| L-B-SER-35 | 0 | -1.62 | 1.6 | 0 | -0.02 |
| L-B-ALA-36 | 0 | 60.85 | -60.08 | 0 | 0.76 |

**Table S7. Residue decomposition analysis for AMP18 when bound to VIM-2**

| RESIDUE_ID | VDW | ELE | GB | SA | TOTAL |
| --- | --- | --- | --- | --- | --- |
| L-B-LYS-1 | -0.05 | -158.23 | 156.77 | 0 | -1.51 |
| L-B-GLU-2 | -0.04 | 82.66 | -81.54 | 0 | 1.08 |
| L-B-LYS-3 | -0.16 | -95.15 | 94.43 | 0 | -0.88 |
| L-B-LYS-4 | -0.85 | -121.43 | 118.92 | -0.44 | -3.8 |
| L-B-LYS-5 | -0.15 | -78.59 | 77.83 | 0 | -0.91 |
| L-B-LYS-6 | -0.11 | -78.89 | 78.1 | 0 | -0.9 |
| L-B-LYS-7 | -3.27 | -155.81 | 156.22 | -0.49 | -3.34 |
| L-B-LYS-8 | -2.47 | -121.06 | 122.17 | -0.56 | -1.93 |
| L-B-LYS-9 | -0.21 | -83.57 | 82.91 | 0 | -0.87 |
| L-B-LYS-10 | -0.24 | -99.23 | 98.5 | 0 | -0.97 |
| L-B-ASN-11 | -2.42 | -10.63 | 9.66 | -0.41 | -3.81 |
| L-B-THR-12 | -1.72 | 1.37 | -0.19 | -0.2 | -0.74 |
| L-B-ASN-13 | -3.95 | -6.06 | 7.7 | -0.89 | -3.2 |
| L-B-ILE-14 | -0.36 | -2.03 | 2.12 | 0 | -0.27 |
| L-B-PHE-15 | -0.86 | 2.08 | -0.56 | -0.09 | 0.58 |
| L-B-LYS-16 | -3.8 | -81.34 | 85.4 | -0.8 | -0.55 |
| L-B-ASN-17 | -2.6 | -0.77 | 2.06 | -0.3 | -1.6 |
| L-B-LYS-18 | -2.56 | -145.33 | 144.81 | -0.75 | -3.84 |
| L-B-LYS-19 | -2.19 | -97.28 | 97.78 | -0.4 | -2.09 |
| L-B-LYS-20 | -0.37 | -80.01 | 79.76 | -0.02 | -0.64 |
| L-B-GLY-21 | -0.05 | -1.56 | 1.69 | 0 | 0.08 |
| L-B-PHE-22 | -0.24 | 1.42 | -1.1 | 0 | 0.08 |
| L-B-PHE-23 | -0.07 | -0.56 | 0.75 | 0 | 0.11 |
| L-B-PHE-24 | -0.63 | 0.66 | -0.4 | -0.13 | -0.49 |
| L-B-PHE-25 | -0.05 | -0.28 | 0.38 | 0 | 0.05 |
| L-B-PHE-26 | -0.31 | 0.19 | 0.22 | 0 | 0.1 |
| L-B-PHE-27 | -3.86 | 0.97 | -0.18 | -0.79 | -3.86 |
| L-B-PHE-28 | -0.15 | 84.7 | -83.07 | 0 | 1.48 |

**Table S8. Residue decomposition analysis for PolyR when bound to VIM-2**

| RESIDUE_ID | VDW | ELE | GB | SA | TOTAL |
| --- | --- | --- | --- | --- | --- |
| L-B-ARG-1 | -0.15 | -172.69 | 171.37 | -0.03 | -1.49 |
| L-B-ARG-2 | -0.16 | -93.47 | 92.8 | 0 | -0.83 |
| L-B-ARG-3 | -0.04 | -71.52 | 70.71 | 0 | -0.84 |
| L-B-ARG-4 | -0.12 | -85.11 | 84.4 | 0 | -0.84 |
| L-B-ARG-5 | -3.29 | -131.06 | 131.59 | -0.74 | -3.5 |
| L-B-ARG-6 | -0.28 | -91.52 | 90.74 | 0 | -1.06 |
| L-B-ARG-7 | -0.1 | -80.37 | 79.54 | 0 | -0.93 |
| L-B-ARG-8 | -2.54 | -119.18 | 119.44 | -0.41 | -2.69 |
| L-B-ARG-9 | -4.02 | -162.06 | 159.74 | -0.9 | -7.24 |
| L-B-ARG-10 | -0.19 | -89.19 | 88.44 | 0 | -0.94 |
| L-B-ARG-11 | -0.14 | -93.46 | 92.53 | 0 | -1.06 |
| L-B-ARG-12 | -5.13 | -177.2 | 175.9 | -1 | -7.44 |
| L-B-ARG-13 | -1.27 | -125.71 | 126.46 | -0.26 | -0.78 |
| L-B-ARG-14 | -0.11 | -89.3 | 88.44 | 0 | -0.97 |
| L-B-ARG-15 | -0.24 | -111.01 | 110.23 | 0 | -1.03 |
| L-B-ARG-16 | -5.74 | -165.03 | 167.59 | -1.14 | -4.32 |
| L-B-ARG-17 | -0.51 | -123.24 | 119.28 | -0.43 | -4.9 |
| L-B-ARG-18 | -0.11 | -88.91 | 88.08 | 0 | -0.95 |
| L-B-ARG-19 | -2.15 | -130.65 | 130.99 | -0.26 | -2.07 |
| L-B-ARG-20 | -4.55 | -114.73 | 118.47 | -0.96 | -1.77 |
| L-B-ARG-21 | -0.19 | -93.68 | 92.77 | 0 | -1.1 |
| L-B-ARG-22 | -0.09 | -89.75 | 88.95 | 0 | -0.9 |
| L-B-ARG-23 | -3.22 | -124.75 | 127.32 | -0.72 | -1.36 |
| L-B-ARG-24 | -0.49 | -104.19 | 103.78 | -0.1 | -1.01 |
| L-B-ARG-25 | -0.06 | -78.8 | 77.95 | 0 | -0.91 |
| L-B-ARG-26 | -0.13 | -95.16 | 94.38 | 0 | -0.92 |
| L-B-ARG-27 | -3.86 | -114.84 | 116.93 | -0.85 | -2.63 |
| L-B-ARG-28 | -0.08 | -80.96 | 80.13 | 0 | -0.91 |
| L-B-ARG-29 | -0.03 | -73.79 | 72.95 | 0 | -0.87 |
| L-B-ARG-30 | -1.23 | -25.18 | 25.76 | -0.28 | -0.93 |

**Table S9. Residue decomposition analysis for PolyF when bound to VIM-2**

| RESIDUE_ID | VDW | ELE | GB | SA | TOTAL |
| --- | --- | --- | --- | --- | --- |
| L-B-PHE-1 | -0.45 | -100.92 | 100.77 | -0.08 | -0.68 |
| L-B-PHE-2 | -0.06 | -0.23 | 0.32 | 0 | 0.03 |
| L-B-PHE-3 | -0.27 | -1.65 | 1.83 | 0 | -0.09 |
| L-B-PHE-4 | -1.7 | 0.65 | 0.12 | -0.47 | -1.41 |
| L-B-PHE-5 | -0.18 | 0.36 | -0.21 | 0 | -0.03 |
| L-B-PHE-6 | -0.63 | -1.54 | 2.18 | -0.06 | -0.05 |
| L-B-PHE-7 | -3.67 | -3.43 | 5.74 | -0.68 | -2.04 |
| L-B-PHE-8 | -5.39 | -0.88 | 0.97 | -0.82 | -6.12 |
| L-B-PHE-9 | -2.62 | -2.69 | 3.01 | -0.37 | -2.66 |
| L-B-PHE-10 | -2.25 | -0.9 | 2.15 | -0.35 | -1.36 |
| L-B-PHE-11 | -5.81 | -1.41 | 2.34 | -0.95 | -5.83 |
| L-B-PHE-12 | -0.71 | 1.83 | -1.44 | -0.02 | -0.35 |
| L-B-PHE-13 | -0.18 | 0.22 | -0.01 | 0 | 0.03 |
| L-B-PHE-14 | -1.39 | -2.24 | 2.8 | -0.21 | -1.05 |
| L-B-PHE-15 | -4.29 | -0.49 | 1.7 | -0.82 | -3.91 |
| L-B-PHE-16 | -1.56 | -0.11 | 0.68 | -0.41 | -1.4 |
| L-B-PHE-17 | -0.06 | -0.95 | 1.03 | 0 | 0.02 |
| L-B-PHE-18 | -0.04 | -2.03 | 2.12 | 0 | 0.05 |
| L-B-PHE-19 | -0.06 | -2.48 | 2.58 | 0 | 0.04 |
| L-B-PHE-20 | -0.37 | -1.49 | 1.76 | -0.05 | -0.16 |
| L-B-PHE-21 | -0.03 | -0.23 | 0.26 | 0 | 0 |
| L-B-PHE-22 | -0.02 | -1.27 | 1.28 | 0 | -0.01 |
| L-B-PHE-23 | -0.05 | -2.01 | 2.05 | 0 | -0.01 |
| L-B-PHE-24 | -0.05 | -0.59 | 0.63 | 0 | -0.01 |
| L-B-PHE-25 | -0.01 | -0.4 | 0.41 | 0 | -0.01 |
| L-B-PHE-26 | -0.01 | -1.52 | 1.52 | 0 | -0.01 |
| L-B-PHE-27 | -0.03 | -1.56 | 1.58 | 0 | -0.01 |
| L-B-PHE-28 | -0.02 | -0.64 | 0.66 | 0 | 0 |
| L-B-PHE-29 | -0.01 | -0.92 | 0.92 | 0 | -0.01 |
| L-B-PHE-30 | -0.01 | 65.82 | -64.96 | 0 | 0.85 |

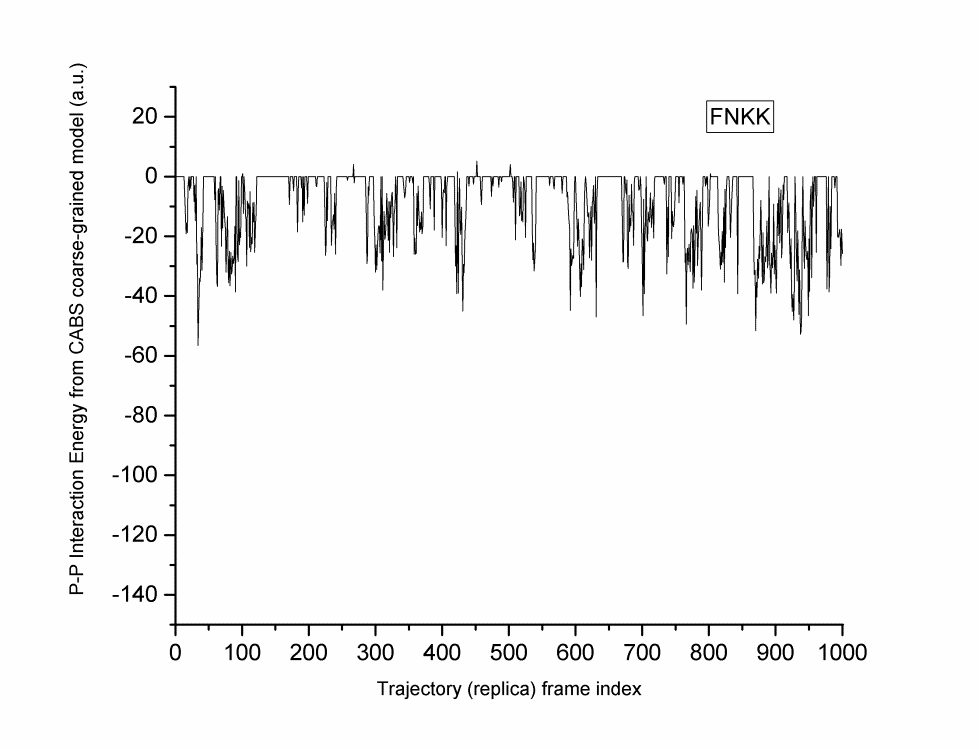

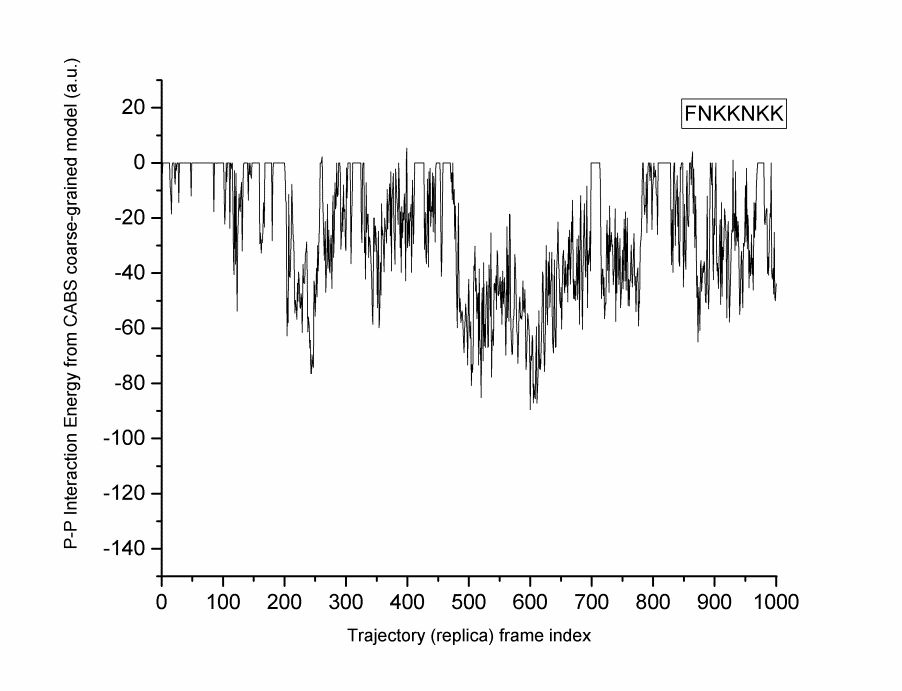

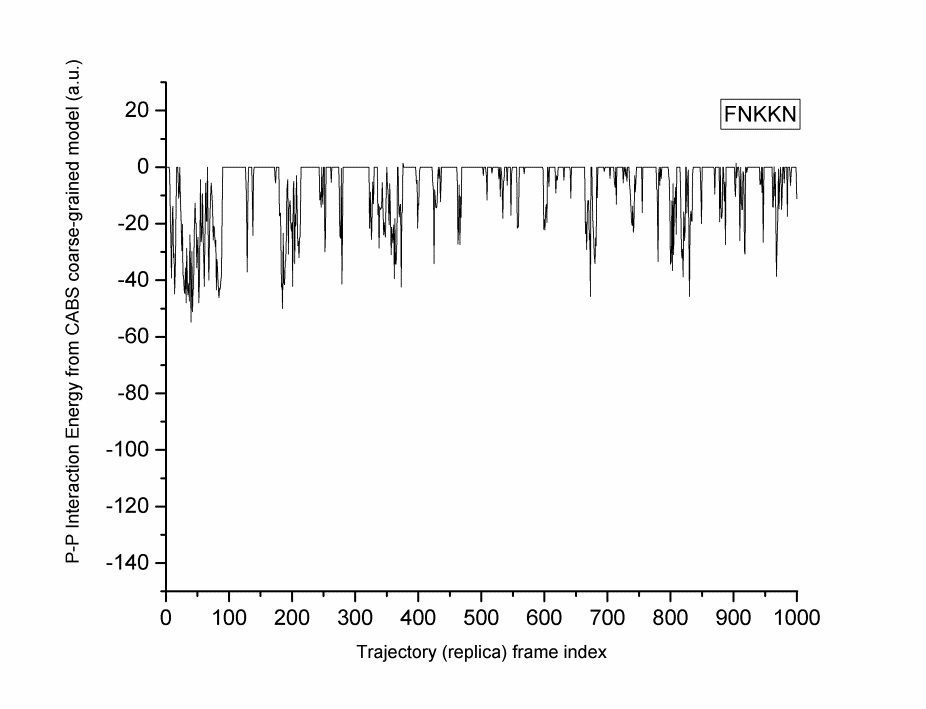

**Figure S1. CABSDOCK trajectory of the phenylalanine-containing analogs belonging to set 1.**

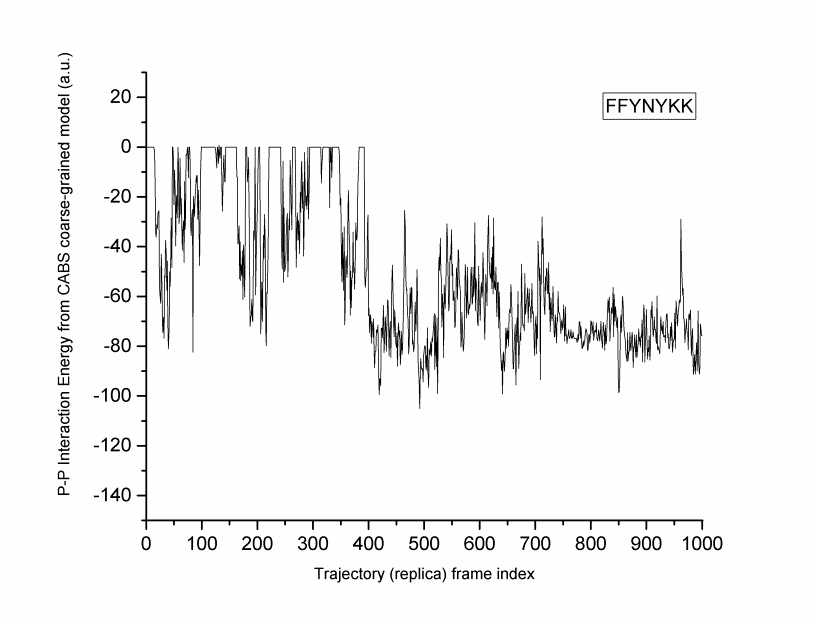

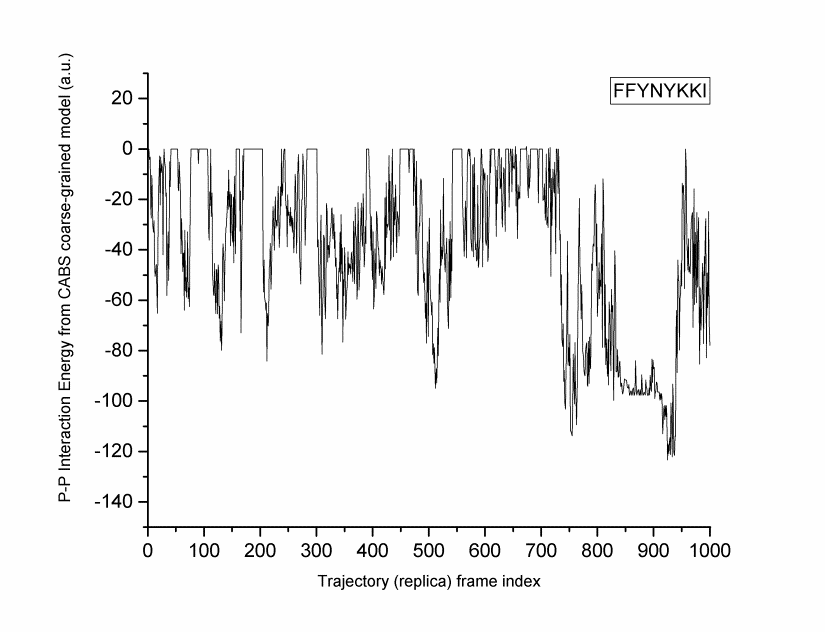

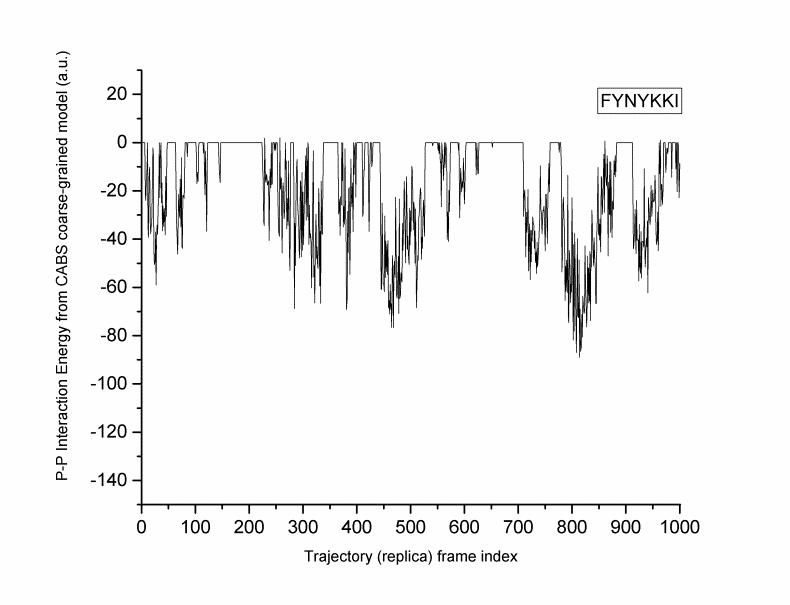

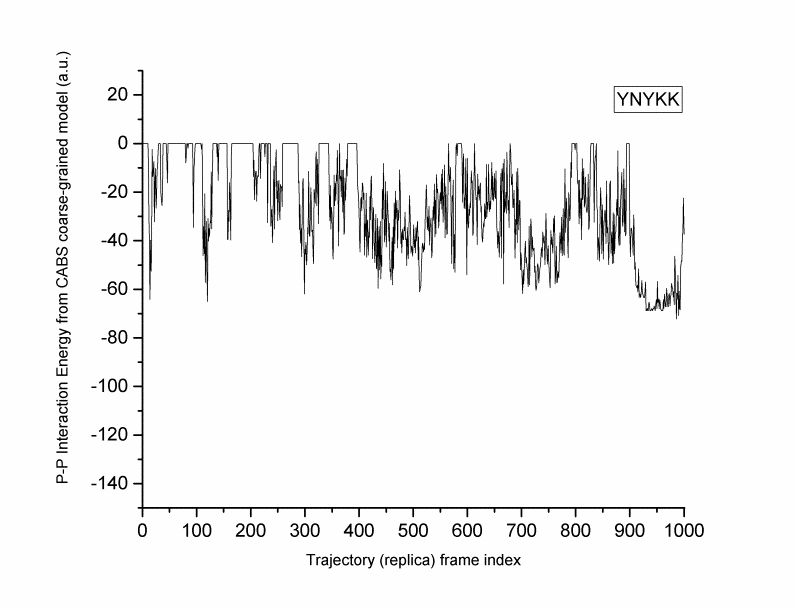

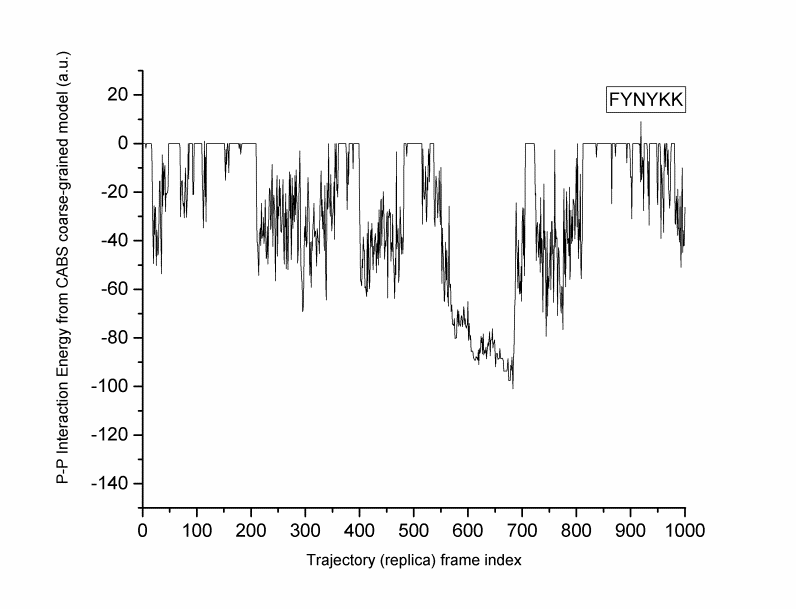

**Figure S2. CABSDOCK trajectory of the phenylalanine-containing analogs belonging to set 3.**

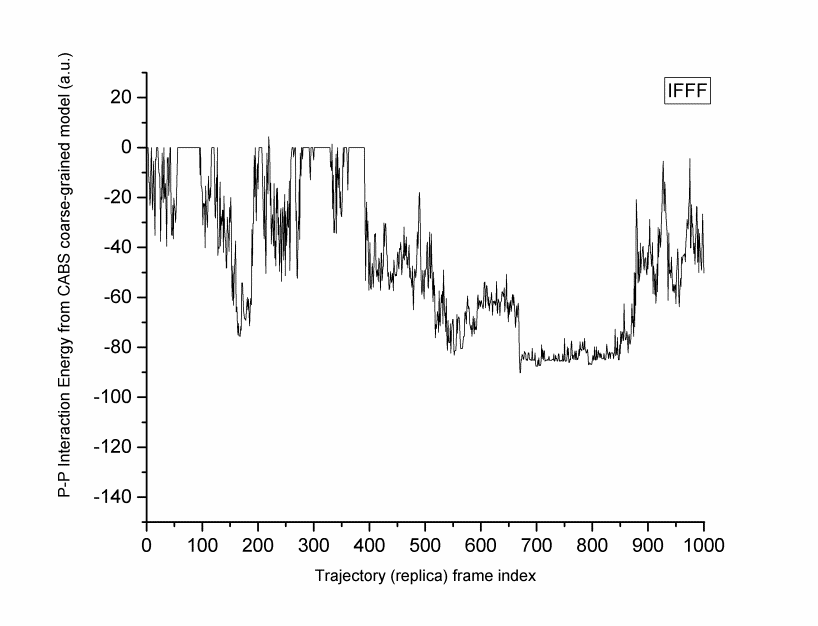

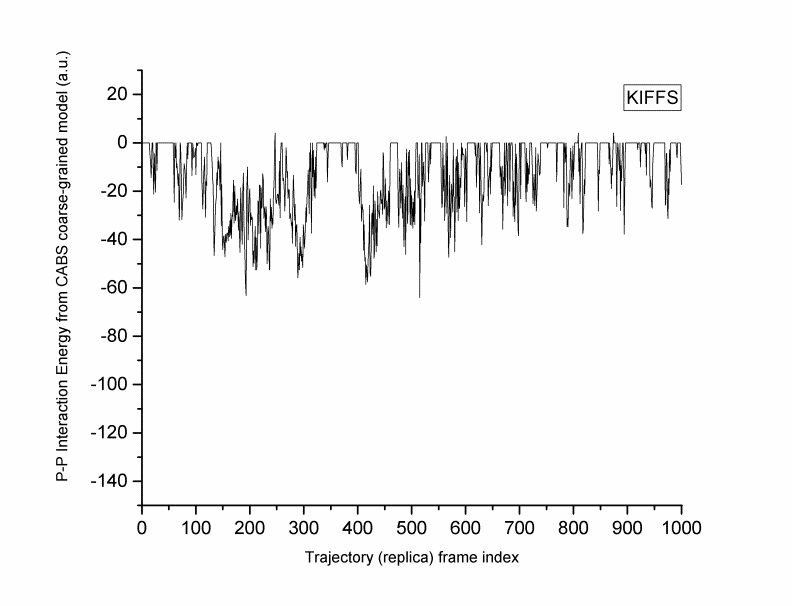

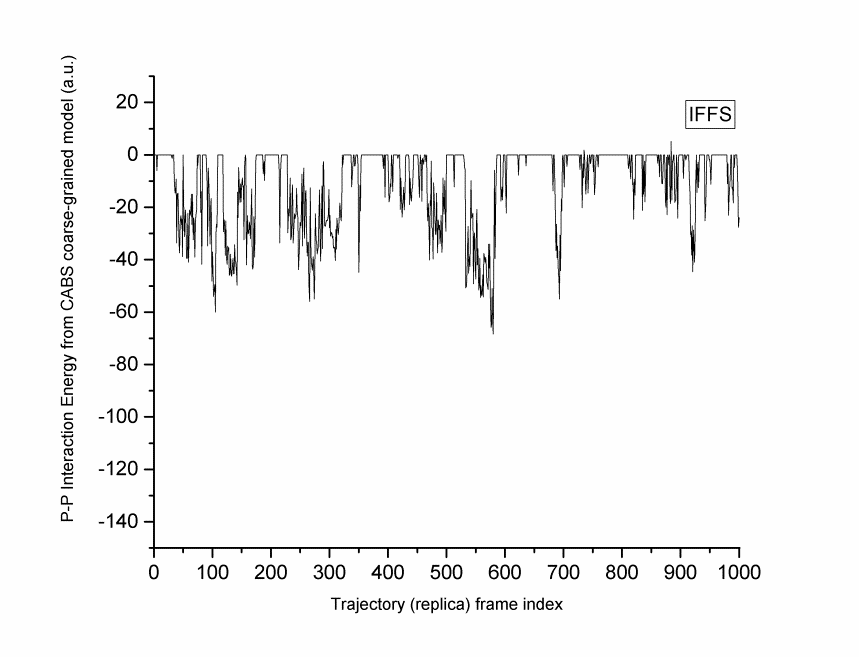

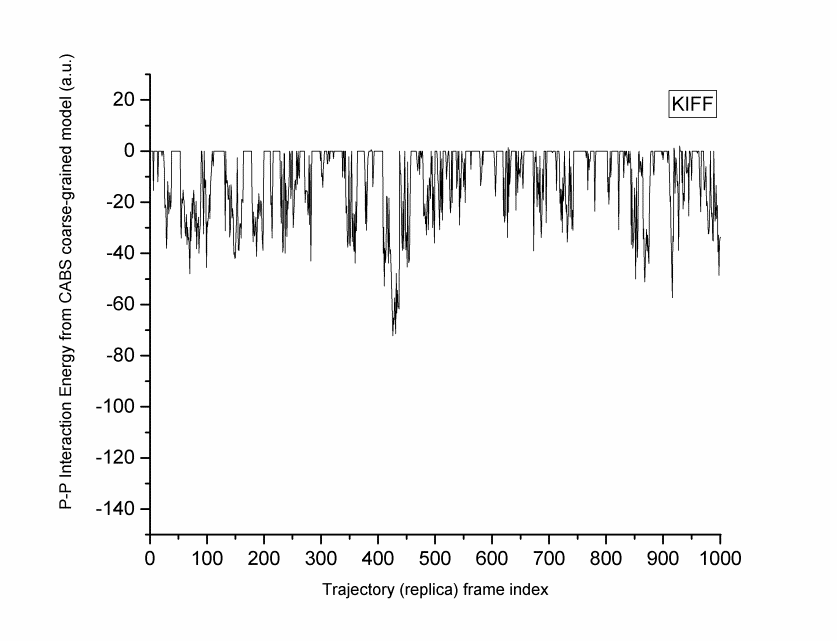

**Figure S3. CABSDOCK trajectory of the phenylalanine-containing analogs belonging to set 4.**

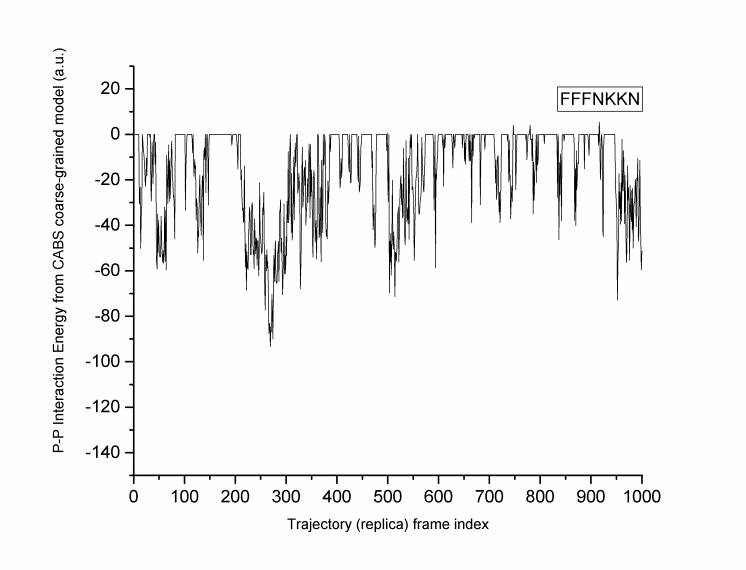

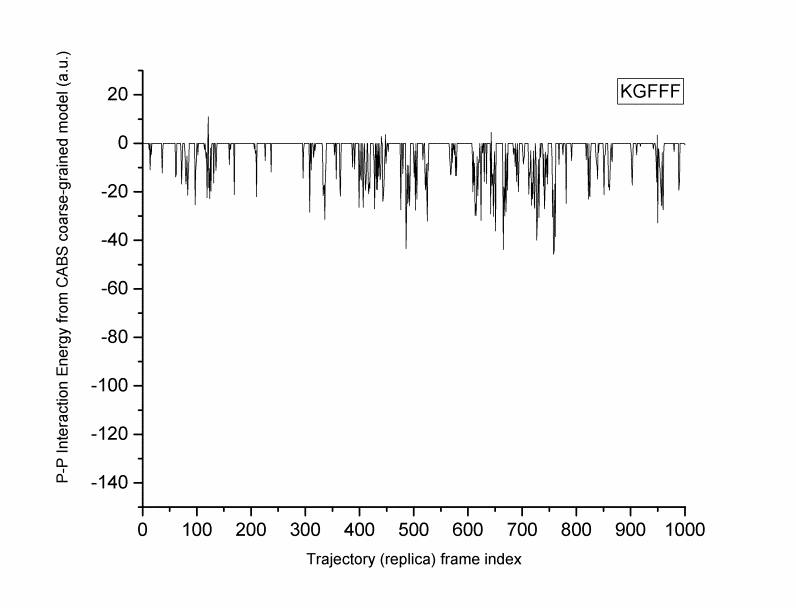

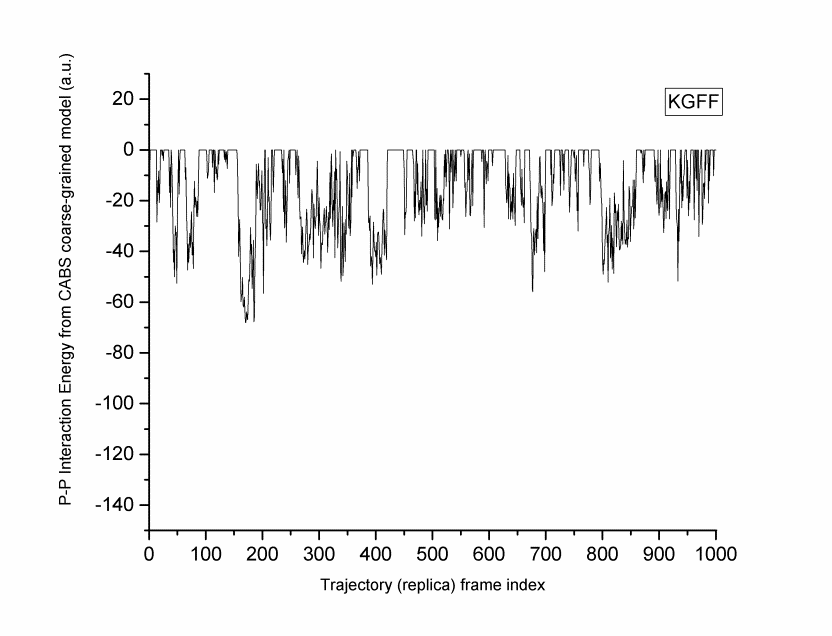
**Figure S4. CABSDOCK trajectory of the phenylalanine-containing analogs belonging to set 5.**

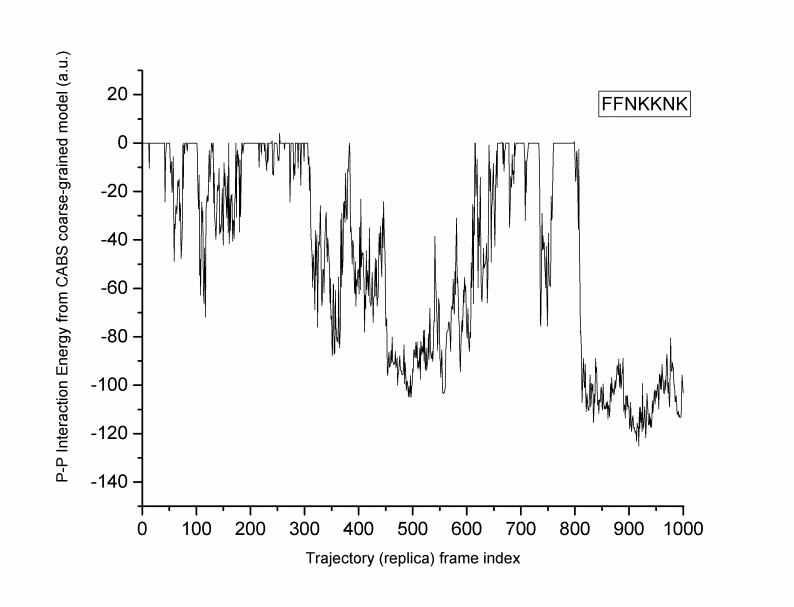

**Figure S5. CABSDOCK trajectory of the phenylalanine-containing analogs belonging to set 6.**

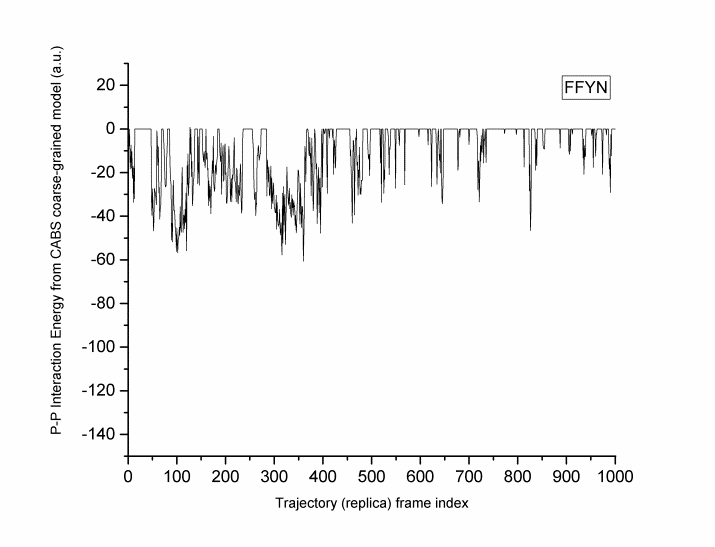

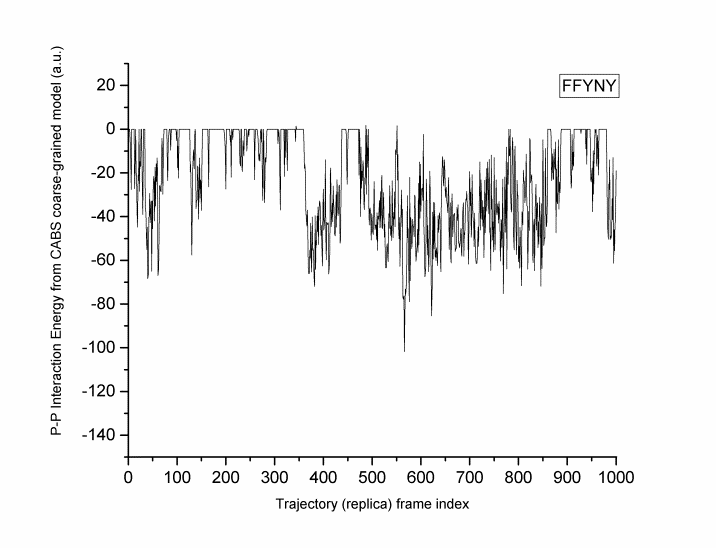

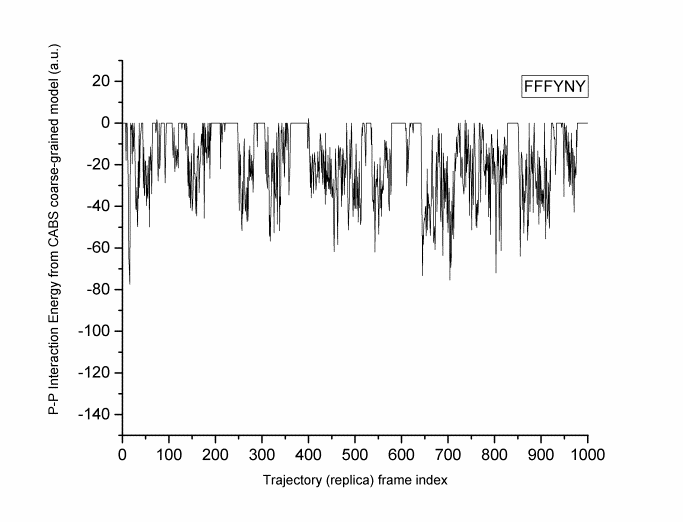

**Figure S6. CABSDOCK trajectory of the phenylalanine-containing analogs belonging to set 7.**

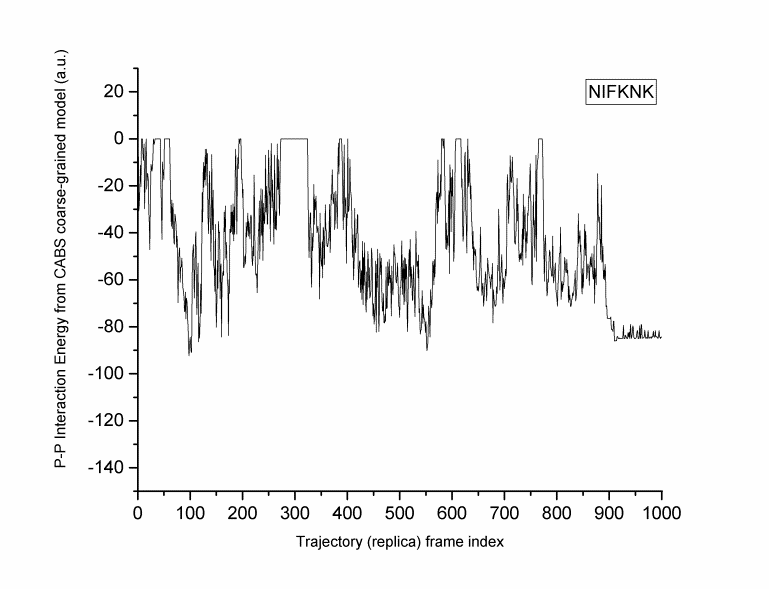

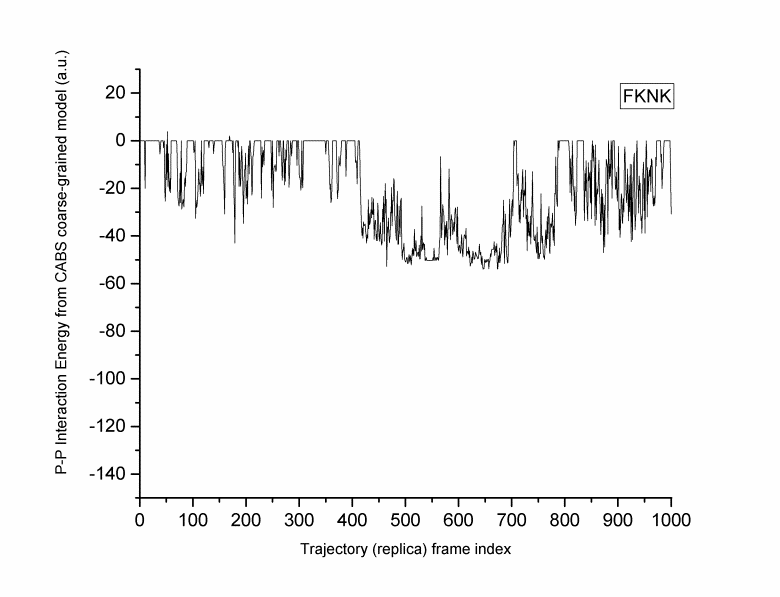

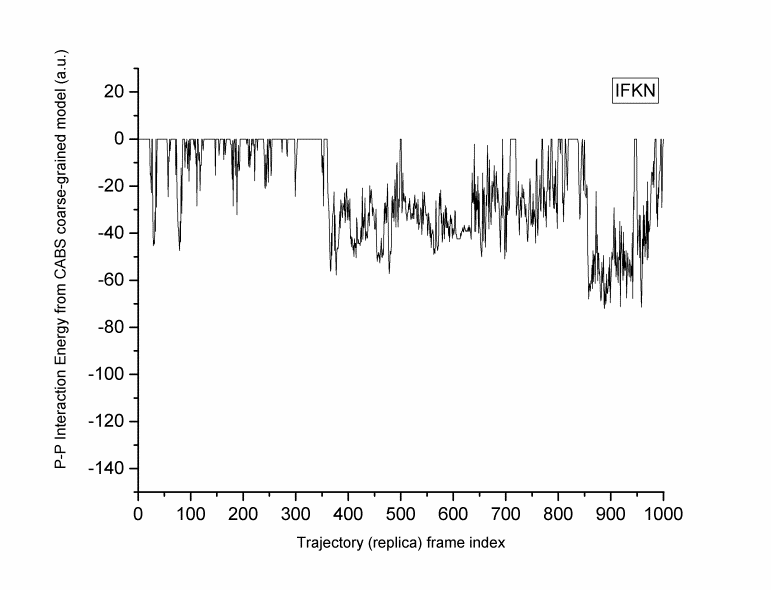

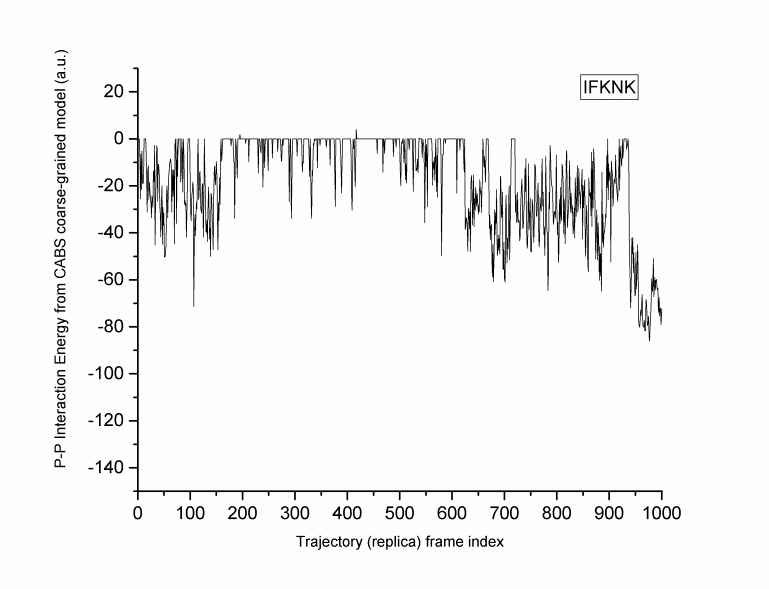

**Figure S7. CABSDOCK trajectory of the phenylalanine-containing analogs belonging to set 8.**

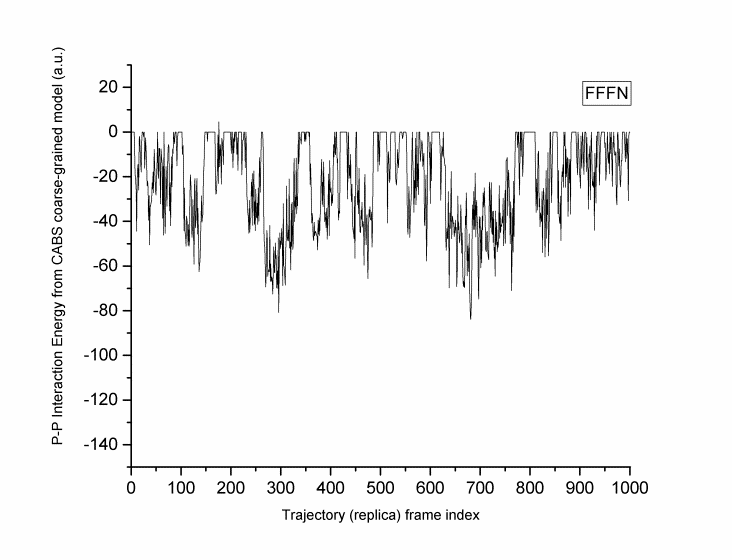

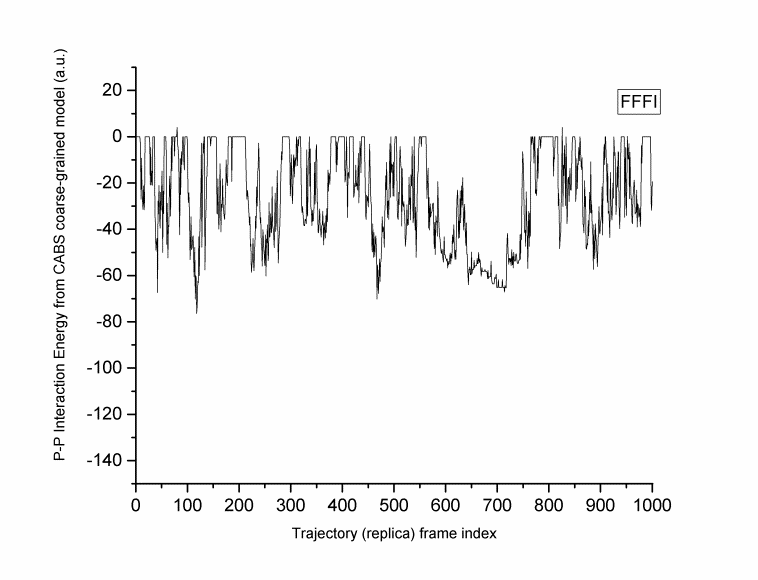

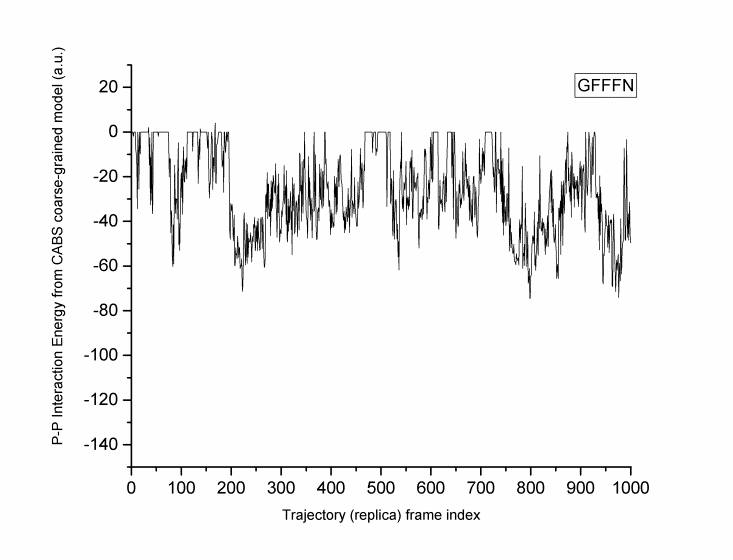

**Figure S8. CABSDOCK trajectory of the phenylalanine-containing analogs belonging to set 9.**

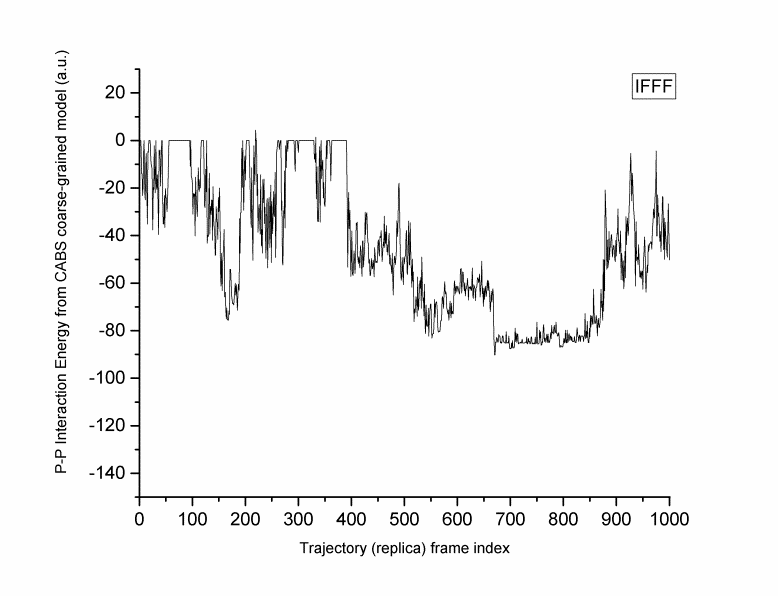

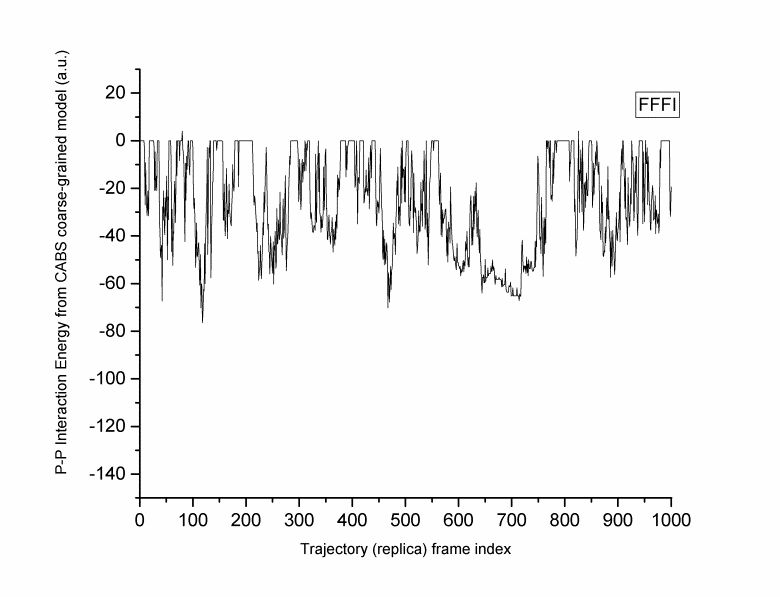

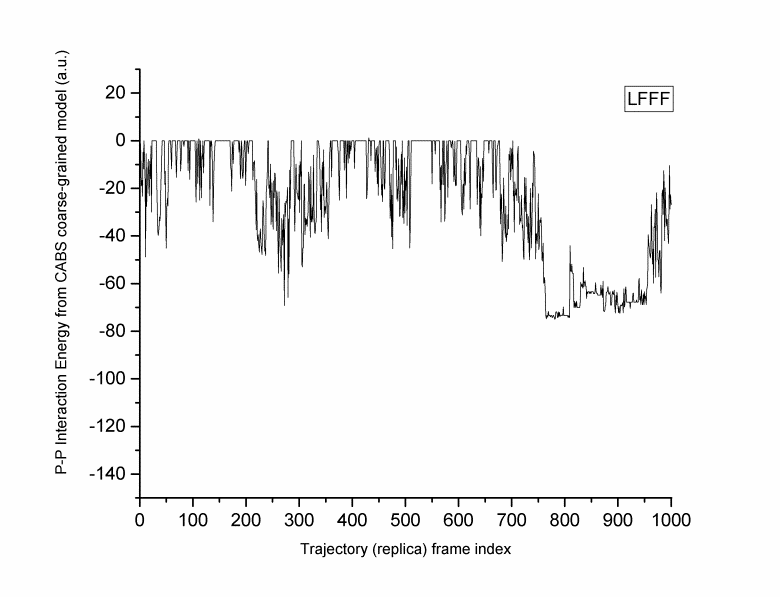

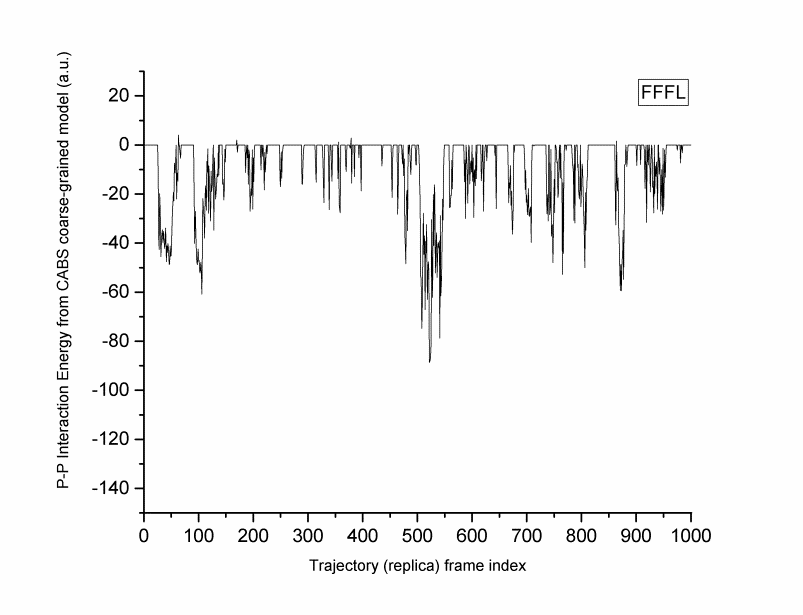

(a)

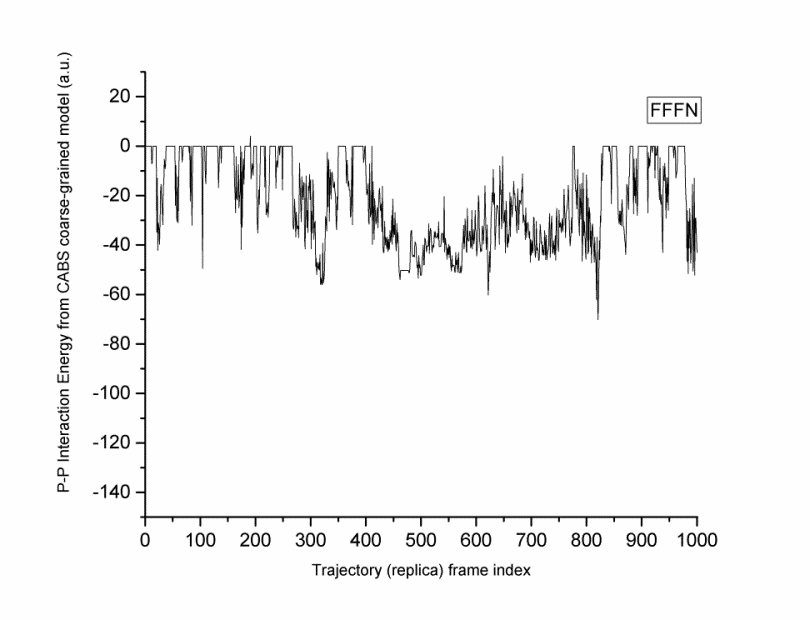

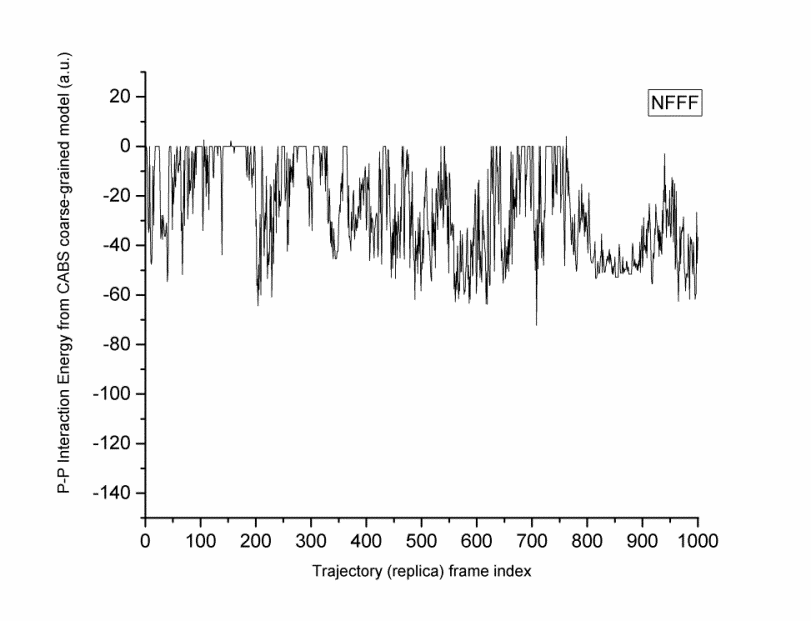

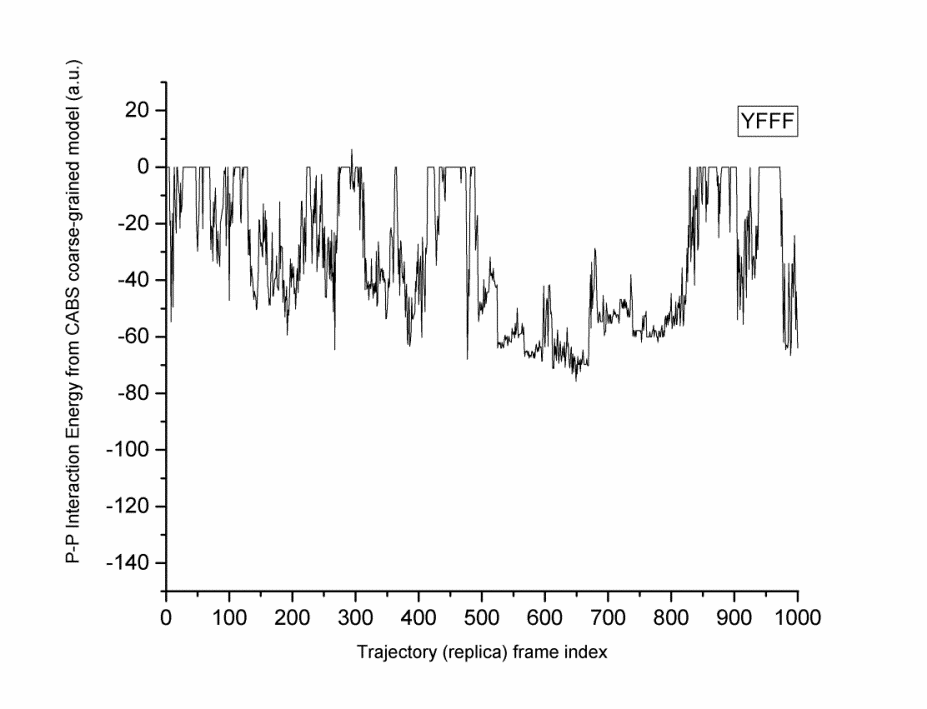

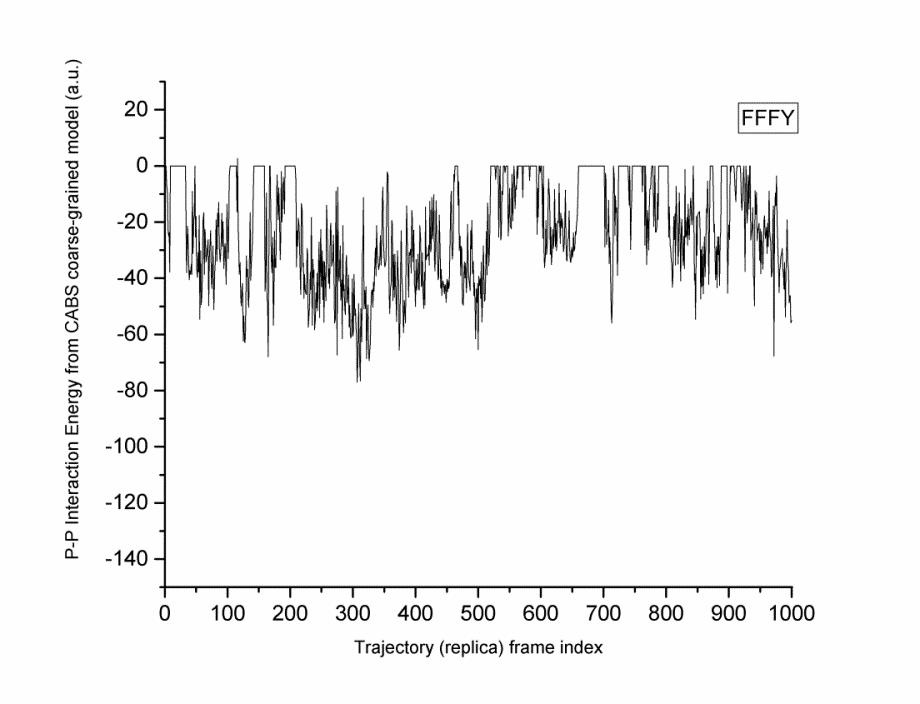

(b)

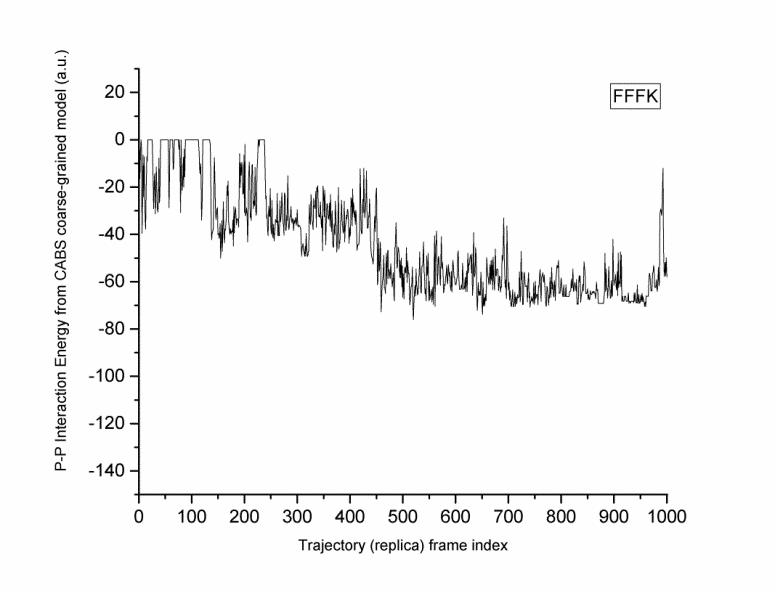

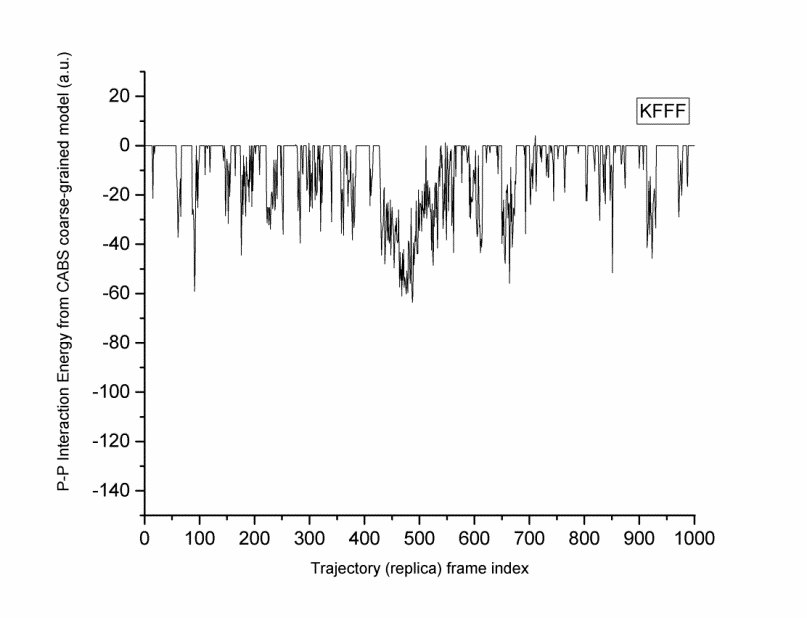

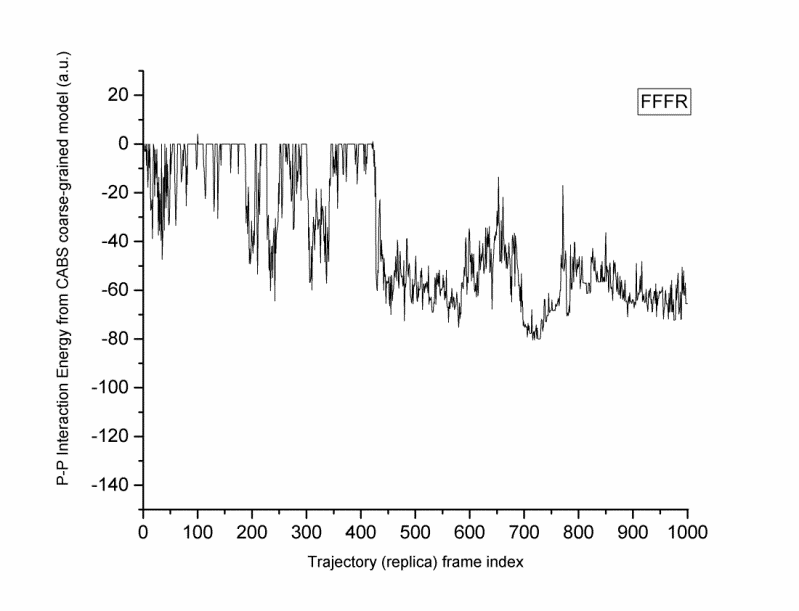

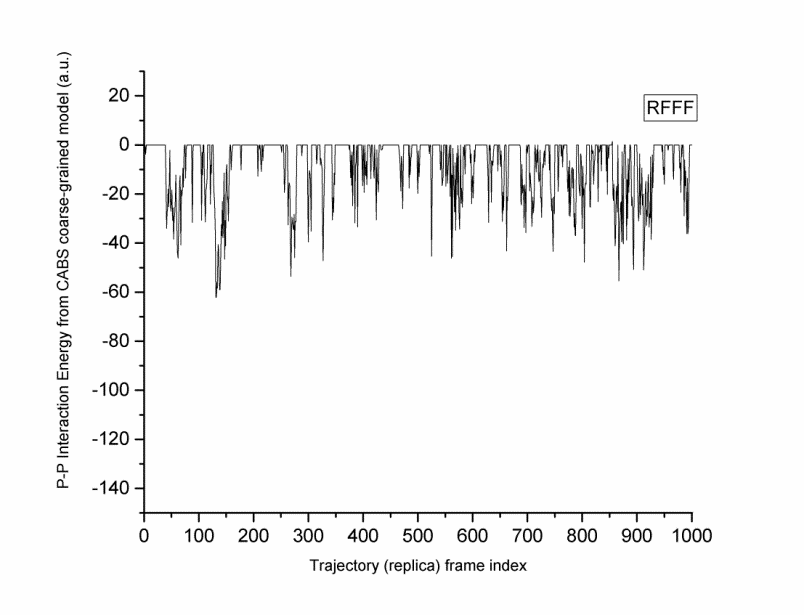

(c)

**Figure S9 (a-c). CABSDOCK trajectory of the phenylalanine-containing analogs belonging to set 10.**

Table S10. **Library I analogs or high binding segments derived from gut-derived parent AMPs**

| Parent AMP | Analog | Design basis | Model 1 Avg. I.E. | All Models Avg. I.E. | Best dock score (Model 1) | MMGBSA of Model 1 | Dock score of best MMGBSA model | Best MMGBSA score |
| --- | --- | --- | --- | --- | --- | --- | --- | --- |
| AMP10 | **5-FNKK-8** | Ligplot | -9.7303 | -16.5484 | -2565.8495 | -29.47 | -1708.15 | -39.98 |
|  | **1-GGFF-4** | Ligplot | -16.9316 | -13.3558 | -1410.50616 | -20.33 | -1020.06 | -20.67 |
|  | **2-GFFFN-6** | Ligplot | -26.204 | -20.2757 | -1664.92496 | -21.71 | -1300.01 | -28.97 |
|  | **5-FNKKNKK-11** | BE | -25.15 | -27.0513 | -2852.41045 | -21.65 | -2789.53 | -39.56 |
|  | **19-IFFS-22** | Ligplot | -11.1993 | -17.8581 | -1397.39799 | -20.42 | -1252.66 | -21.9 |
|  | **3-FFFNKKN-9** | BE | -15.6614 | -32.4537 | -2544.56142 | -31.64 | -2314.84 | -45.87 |
|  | **18-KIFFS-22** | BE | -12.4272 | -21.4024 | -1885.98259 | -29.5 | -1885.98 | -29.5 |
|  | **4-FFNKKNK-10** | BE | -46.5289 | -30.2705 | -2518.94678 | -27.97 | -2441.58 | -39.74 |
| AMP13 | **33-IIFRAR-38** | Ligplot | -43.0497 | -28.5221 | -2630.87406 | -27.26 | -2515.73 | -46.43 |
|  | **35-FRAR-38** | Ligplot | -9.84635 | -18.8469 | -2729.16122 | -38.68 | -2412.05 | -48.5 |
|  | **32-KIIFRAR-38** | BE | -12.5664 | -29.43 | -2741.05653 | -49.24 | -2741.06 | -49.24 |
|  | **27-FYNYKKI-33** | BE | -17.8018 | -30.4676 | -2499.66614 | -24.18 | -2228.96 | -52.86 |
|  | **33-IIFR-36** | BE | -24.4461 | -19.108 | -2022.29028 | -18.78 | -1718.39 | -27.44 |
|  | **11-IFFFFFL-17** | BE | -22.0615 | -34.4525 | -1941.91809 | 45895312 | -1729.68 | -40.72 |
|  | **11-IFF-13** | Ligplot | NA | NA | -1729.26174 | -10.67 | -1143.38 | -19.9 |
|  | **26-FFYNY-30** | Ligplot | -23.3958 | -27.9583 | -1952.57173 | -33.89 | -1952.57 | -33.89 |
|  | **26-FFYNYKK-32** | BE | -50.3474 | -34.5593 | -2572.98141 | -26.2 | -2296.36 | -39.57 |
|  | **25-FFFYNY-30** | Ligplot | -16.9108 | -34.6702 | -2018.81572 | -39.92 | -1742.43 | -39.92 |
| AMP18 | **15-FKNKK-19** | BE | -16.5269 | -20.5455 | -2633.27176 | -37.12 | -1886.66 | -37.52 |
|  | **13-NIFKNK-18** | BE | -44.6829 | -23.7738 | -2353.30768 | -27.97 | -1870.39 | -38.7 |
|  | **20-KGFF-23** | BE | -13.7976 | -18.3011 | -1965.08694 | -31.24 | -1437.49 | -37.99 |
|  | **20-KGFFF-24** | BE | -3.67655 | -22.327 | -1963.01719 | -33.64 | -1673.45 | -44.71 |

**Table S11. Library II containing potential analogs found within analogs (AWA)**

| AMP | Parent Sequence | Analog Sequence | Model 1 Avg. I.E. | All Models Avg. I.E. | Best dock score (Model 1) | MMGBSA of Model 1 | Dock score of best MMGBSA model | Best MMGBSA score |
| --- | --- | --- | --- | --- | --- | --- | --- | --- |
| AMP10 | 2GFFFN6 | **FFFN** | -21.967 | -22.979 | -1681.5 | -27.32 | -1681.46 | -27.32 |
|  | 3FFFNKKN9 | **FFNK** | -15.951 | -16.809 | -1817.6 | -24.04 | -1678.76 | -28.3 |
|  | 3FFFNKKN9 | **FNKK** | -9.7303 | -16.548 | -2565.8 | -29.47 | -1708.15 | -39.98 |
|  | 3FFFNKKN9 | **FNKKN** | -6.7217 | -20.689 | -2322.7 | -25.69 | -1907.32 | -31.05 |
|  | 18KIFFS22 | **IFFS** | -11.199 | -17.858 | -1397.4 | -20.42 | -1252.66 | -21.9 |
|  | 18KIFFS22 | **KIFF** | -10.772 | -17.73 | -1792.3 | -18.65 | -1475.39 | -27.49 |
| AMP13 | 33IIFRAR38 | **IIFRA** | -12.509 | -20.254 | -2358.1 | -33.89 | -2193.87 | -41.46 |
|  | 33IIFRAR38 | **IFRA** | -6.3665 | -16.296 | -1976.8 | -22.41 | -1807.7 | -30.57 |
|  | 32KIIFRAR38 | **IIFR** | -24.446 | -19.108 | -2022.3 | -18.78 | -1718.39 | -27.44 |
|  | 26FFYNY30 | **FFYN** | -9.9736 | -20.642 | -1853.4 | -27.19 | -1505.12 | -32.66 |
|  | 26FFYNYKK32 | **FYNY** | -17.764 | -22.65 | -1941.4 | -19.64 | -1598.51 | -25.16 |
|  | 11IFFFFFL17 | **IFFF** | -45.025 | -21.36 | -1302 | -31.1 | -1301.97 | -31.1 |
|  | 11IFFFFFL17 | **FFFL** | -8.7247 | -19.575 | -1313.2 | -7.85 | -1230.95 | -25.58 |
| AMP18 | 15FKNKK19 | **FKNK** | -19.917 | -17.459 | -2257.7 | -31.18 | -2257.67 | -31.18 |
|  | 13NIFKNK18 | **NIFK** | -24.035 | -16.372 | -2121.3 | -15.18 | -1666.15 | 27.12 |
|  | 13NIFKNK18 | **IFKNK** | -17.536 | -20.229 | -2161.8 | -26.51 | -1991.05 | -32.27 |
|  | 13NIFKNK18 | **IFKN** | -22.107 | -18.189 | -1805.1 | -16.91 | -1766.87 | -27.72 |

**Table S12. Library III of analogs containing custom sequences**

| **Analog** | **Model 1 Avg. I.E.** | **All Models Avg. I.E.** | **Best dock score (Model 1)** | **MMGBSA of Model 1** | **Dock score of best MMGBSA model** | **Best MMGBSA score** |
| --- | --- | --- | --- | --- | --- | --- |
| FYNYKK (Intermediates lying in between FYNYKKI -> FFYNYKK) | -26.8189 | -29.7932 | -2189.08 | -34.56 | -2189.08 | -34.56 |
| YNYKK (Intermediates lying in between FYNYKKI -> FFYNYKK) | -25.3562 | -23.7678 | -2178.23 | -30.91 | -2131.94 | -37.93 |
| FFYNYKKI (Intermediates lying in between FYNYKKI -> FFYNYKK) | -38.0072 | -36.6917 | -2265.54 | -28.81 | -2204.57 | -42.13 |
| KFKFKF | -13.5823 | -27.2175 | -2182.57 | -24.99 | -2021.26 | -38.13 |
| FKFKFK | -39.7546 | -28.146 | -2128.73 | -29.75 | -2128.73 | -29.75 |
| NFNFNF | -12.5419 | -28.0364 | -2185.58 | -18.6 | -1963.92 | -32.17 |
| FNFNFN | -51.7015 | -28.1562 | -1928.43 | -14.38 | -1849.56 | -25.13 |
| FNKFNK | -40.3799 | -25.2319 | -2195.31 | -26.64 | -1669.98 | -32.5 |
| KNFKNF | -14.8934 | -28.1336 | -2418.27 | -21.31 | -2182.32 | -27.6 |
| KFKFKFF | -53.5749 | -32.868 | -2070.34 | -19.76 | -2047.03 | -36.34 |
| FFKFKFK | -33.7979 | -32.3448 | -2403.91 | NA | -2007.59 | -36.19 |
| NFNFNFF | -71.4044 | -33.33 | -2163.35 | -27.9 | -1620.04 | -28.29 |
| FFNFNFN | -61.7073 | -34.0312 | -2411.82 | -29.21 | -2195.76 | -37.28 |
| FFNKFNK | -64.4905 | -31.0497 | -2170.14 | -30.89 | -2051.86 | -41.59 |
| KNFKNFF | -38.8272 | -32.1876 | -2449.94 | -40.39 | -2257.42 | -41.78 |
| FYNYKK | -26.8189 | -29.7932 | -2189.08 | -34.56 | -2189.08 | -34.56 |
| NFKNFK | -11.3546 | -26.5392 | -2627.12 | -22.63 | -1934.4 | -31.21 |
| KFNKFN | -12.9065 | -24.4148 | -2072.27 | -16.61 | -1680.33 | -32.6 |
| LFFF | -24.5054 | -24.4556 | -1305.44388 | -28.3 | -1197.22232 | -39.47 |
| FFFI | -25.4897 | -22.4361 | -1219.171282 | -15.84 | -1191.229568 | -36.18 |
| NFFF | -25.1819 | -21.1696 | -1554.941253 | NA (error) | -1429.910287 | -27.84 |
| YFFF | -31.8679 | -23.8273 | -1691.285642 | -26.52 | -1268.323465 | -32.84 |
| FFFY | -22.6547 | -22.9186 | -1636.009347 | -14.42 | -1304.523904 | -28.2 |
| RFFF | -8.3469 | -22.4436 | -2152.882301 | -35.78 | -1818.026778 | -38.66 |
| FFFR | -38.4891 | -22.1052 | -2041.051777 | -24.46 | -1693.599229 | -30.66 |
| KFFF | -9.70395 | -19.6445 | -1987.621937 | -25.84 | -1317.483089 | -37.01 |
| FFFK | -43.9577 | -21.4863 | -1612.534049 | -20.03 | -1575.696861 | -35.01 |

**Table S13.** Similarity-based sets (SBS) and their analogs

| **Set 1** | FNKK |
| --- | --- |
|  | FNKKN |
|  | FNKKNKK |
| **Set 2** | IIFRAR |
|  | FRAR |
|  | KIIFRAR |
|  | IIFR |
|  | IIFRA |
|  | IFRA |
| **Set 3** | FFYNYKK |
|  | FYNYKKI |
|  | FYNYKK |
|  | FFYNYKKI |
|  | YNYKK |
| **Set 4** | KIFF |
|  | IFFS |
|  | KIFFS |
|  | IFFF |
| **Set 5** | KGFFF |
|  | KGFF |
| **Set 6** | FFNKKNK |
|  | FFFNKKN |
| **Set 7** | FFYNY |
|  | FFFYNY |
|  | FFYN |
| **Set 8** | NIFKNK |
|  | FKNK |
|  | IFKNK |
|  | IFKN |
| **Set 9** | GFFFN |
|  | FFFN |
|  | FFFL |
| **Set 10** | IFFF |
|  | FFFL |
|  | FFFI |
|  | LFFF |
|  | YFFF |
|  | FFFY |
|  | NFFF |
|  | FFFN |
|  | RFFF |
|  | FFFR |
|  | KFFF |
|  | FFFK |
